# ACPC-MAP: A protocol for manually aligning structural T1-weighted Magnetic Resonance Images to the Anterior Commissure – Posterior Commissure plane according to clinical standards

**DOI:** 10.64898/2026.01.12.699070

**Authors:** Ashok K. Selvaraj, Ronald H.M.A. Bartels, Christian F. Beckmann, R. Saman Vinke, Koen V. Haak

## Abstract

Realigning structural MR images to stereotactic AC-PC space is a standard procedure that enhances anatomical consistency both within and across neuroimaging studies, allowing for precise spatial localization and reliable cross-subject comparisons in both clinical and research contexts. However, different versions of AC-PC spaces, stemming from varying definitions of the AC-PC axis as outlined in common stereotactic atlases like Talairach and Schaltenbrand, often lead to discrepancies between the neurosurgical and neuroimaging communities. Manual realignment of structural MR images to the clinically used version of the AC-PC plane is often necessary to validate the results of automated methods or to realign the image when these methods fail. Furthermore, manual realignment provides a critical ‘ground truth’ for the development of new automated tools. However, such manual interventions are typically performed in a non-standardised manner by domain experts who possess the specialized knowledge of neuroanatomy and neuroimaging required to ensure accurate alignment. To address these challenges, we have developed and validated a standardized protocol for manually realigning structural MR images to the clinically used Schaltenbrand AC-PC plane, using a set of visual criteria to ensure accurate realignment. This protocol can be used to manually align images, verify results, validate existing automated methods, or generate ground truth data for developing new automated techniques.

## 1. Introduction

Realigning Magnetic Resonance Imaging (MRI) data to the Anterior Commissure (AC) - Posterior Commissure (PC) plane is a crucial initial step in stereotactic neurosurgical planning and the development of a wide range of neuroimaging applications (Hamilton et al., 2017; Hebb & Miller, 2010; Hyam et al., 2015; Miller et al., 2019; Mottonen et al., 2015; Nowinski et al., 2005; Starr et al., 1999). Such realignment standardizes the overall image representation, presenting individual brain anatomy in a consistent and organized manner (Choi et al., 2013; Park et al., 2010; Weiss et al., 2003). The AC-PC realignment simplifies systematic navigation through the brain, facilitating a clearer and more accurate depiction of brain structure and function (Coenen et al., 2008; Fox et al., 1984; Minoshima et al., 1993; Starr et al., 1999).

The AC-PC space, frequently referenced in the literature as ‘stereotactic space’, denotes the coordinate system in which anatomical data are embedded (Lau, 2022). This space is commonly described using spatial Cartesian (x, y, z) coordinates relative to a fixed origin, defined by either the AC, the PC, or the midpoint between them, the midcommissural point (MCP) (Alaminos Bouza, 2014; Bejjani et al., 2000; Fox & Kall, 1987; Horn et al., 2017; Kandelʹ & Walker, 1989). Atlases created using this principle are known as stereotactic atlases. Two widely used stereotactic atlases in clinical neuroscience are the Talairach and Schaltenbrand atlases.

The widely recognized Talairach atlas, developed by Jean Talairach, served as a crucial resource for targeting deep brain structures through an AC-PC based coordinate system (Talairach & Tournoux, 1988). Although revolutionary at the time, this atlas was prepared from a single cadaveric brain and had important limitations, such as limited orientation (sagittal and coronal) and an assumption of symmetrical hemispheres (Evans et al., 2012; Mazziotta et al., 1995). Another significant contribution came from Georges Schaltenbrand, who developed a groundbreaking stereotactic atlas using 111 brains (Schaltenbrand et al., 1977). This atlas provides comprehensive representation in all three anatomical planes, effectively addressing the limitations of the Talairach atlas (Lau, 2022).

With the advent of MRI, the neuroimaging community sought to develop a common brain template based on structural MR images that represents the entire population rather than a single brain, like the Talairach atlas (Evans et al., 2012). The Montreal Neurological Institute (MNI) 305 template was an attempt to create a common stereotactic space using 305 high-resolution MR images (Evans et al., 2012). The original T1-weighted images were realigned to approximate the Talairach-defined AC–PC stereotaxic space, following the manual identification of the AC, PC and other neuroanatomical landmarks (Evans et al., 1993). These realigned images were then averaged to create an initial template, which served as the basis for registering the original T1 images to develop the MNI 305 template (Collins et al., 1994; Evans et al., 1993; Evans et al., 1992). Since then, there has been continuous development in MNI templates (Evans et al., 2012), and the International Consortium for Brain Mapping (ICBM) has currently adopted the MNI152 image as a standard reference template for human brain mapping (Brett et al., 2002; Chau & McIntosh, 2005).

The concurrent use of three distinct stereotactic spaces Talairach, Schaltenbrand, and MNI152 has led to inconsistencies within the clinical neuroscience community (Horn et al., 2017). Further complicating this issue, the neurosurgical community, which predominantly adopts the Schaltenbrand-defined AC–PC space, has introduced slight modifications to the definition of the AC–PC line. A comprehensive comparison of these AC–PC spaces is presented in Table 1 and Figure 1.

**Figure 1.**
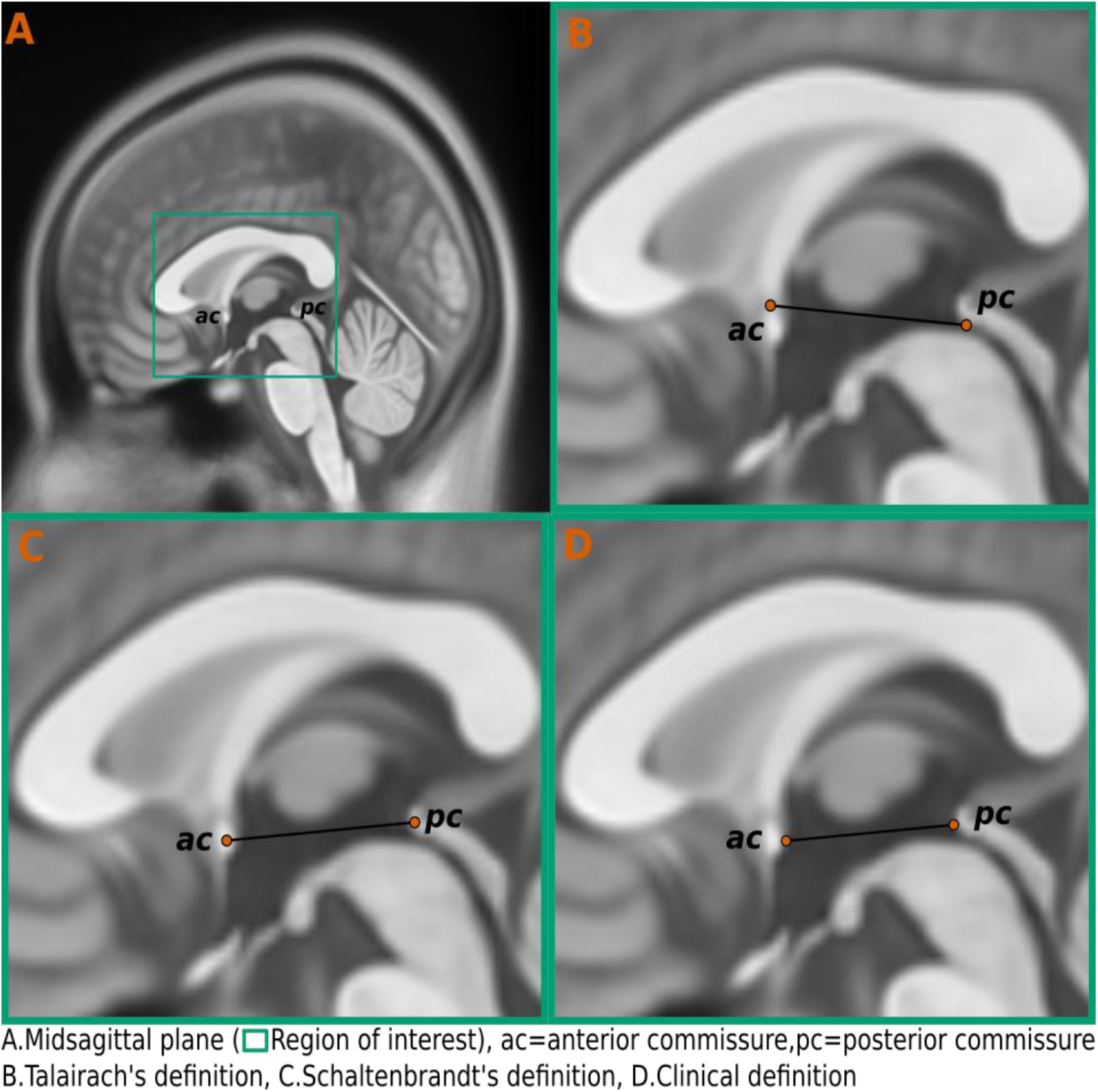
Definitions of the AC-PC line. This illustration presents the various interpretations of the AC-PC line that are commonly used within the neuroscience community. A. Midsagittal plane. The box indicates the region of interest shown in B-D. AC: Anterior Commissure. PC: Posterior Commissure. B. Talairach’s definition. C. Schaltenbrandt’s definition. D. Clinical definition.

**Table 1.** Comparison of the Three Commonly Employed Stereotactic Spaces

| **Stereotactic Space** | **AC-PC line definition** | **Origin** | **Coordinate system** | **Scaling** |
| --- | --- | --- | --- | --- |
| Talairach | From the superior edge of the AC to the inferior edge of the PC (Fig. 1B) | The point where the tangential AC-PC line intersects with the vertical line drawn to the posterior border of the AC. | + x is right,<br>+y is anterior,<br>+z is superior | Piecewise linear scaling |
| Schaltenbrand | From the centre of the AC to the centre of the PC (Fig. 1C) | The point located at the midpoint of the AC-PC line, known as the midcommissural point (MCP). | + x is right,<br>+y is anterior,<br>+z is superior | No scaling |
| MNI | The MNI templates adopted an approximation of Talairach's AC-PC line definition; however, no explicit AC-PC line definition is required to realign the image to MNI space. | The point is located 3.5 mm above the original Talairach origin. | + x is right,<br>+y is anterior,<br>+z is superior | Linear Scaling |

**Table 2.** RMSD values (in mm), computed from rotation matrices, quantifying the deviation of experts manual transformations from the baseline.

| Without guideline |  |  |  |  |
| --- | --- | --- | --- | --- |
| | $R_{min}$ | $R_{avg}$ | $R_{max}$ | $Mean$ |
| Expert 1 | 1.45 | 1.07 | 1.74 | 1.42 |
| Expert 2 | 2.17 | 1.29 | 0.60 | 1.35 |
| With guideline |  |  |  |  |
| Expert1 | 2.04 | 1.22 | 0.85 | 1.37 |
| Expert2 | 0.85 | 1.22 | 1.54 | 1.20 |

The term "AC-PC realignment" fundamentally refers to the process of aligning the structural MR image to a standard orientation. In this framework, the Y-axis is defined by the AC-PC line, running from the AC to the PC. The Z-axis is oriented perpendicular to the AC-PC line, extending from superior to inferior. The X-axis, which runs from left to right, is defined as being perpendicular to the Y and Z axes. Both Y and Z axes are aligned within the longitudinal fissure, or the midline of the brain (Cox, 1996; Lancaster et al., 1995). While these orientations are similar to the aforementioned stereotactic space, the way the AC-PC line is defined and how overall alignment is achieved can vary across different stereotactic spaces (Cox, 1996; Glasser et al., 2013; Horn et al., 2017; Saint-Cyr et al., 2002). This variation leads to differing interpretations of AC-PC realignment, depending on the specific field of application, such as neuroimaging and neurosurgery.

Researchers often use the Talairach coordinate system or atlas-defined AC-PC realignment as a two-stage process known as the Talairach transformation. This process begins with rotating the image to align it with the AC-PC plane. Following this initial alignment, the image is then rescaled to correspond with the anatomical landmarks of the Talairach atlas brain (Brett et al., 2002; Cox, 1996). In stereotactic neurosurgery, it involves rotating the image in the three cardinal (x, y, z) planes so that the AC and PC appear at the same level in the horizontal plane, as defined by the Schaltenbrand atlas (Caire et al., 2011; Saint-Cyr et al., 2002). For the neuroimaging community, "AC-PC realignment" typically refers to the rigid-body registration of a structural MRI image to a standardized stereotactic template, most commonly the MNI152. This process utilizes 6 degrees of freedom (DOF) for rotation and translation, which are derived from an initial 12 DOF registration process (Glasser et al., 2013).

While the majority of researchers in the neuroimaging community have transitioned from Talairach space to MNI stereotactic space for anatomical standardization, the neurosurgical community by and large continues to rely on a Schaltenbrand space (Horn et al., 2017). This framework remains widely used to communicate findings and develop new methods and tools tailored to clinical applications (Bot et al., 2018; Caire et al., 2013; Chakravarty et al., 2006; Duchin et al., 2018; Hyam et al., 2015)

The foundation for successful AC-PC realignment is established through two essential steps. The initial midsagittal realignment involves adjusting the brain along the midsagittal plane to ensure symmetry and proper anatomical orientation. The subsequent AC-PC Realignment, focuses on aligning the desired AC-PC line along the brain’s anteroposterior axis (Lancaster et al., 1995). Both phases of alignment are made possible by the precise definition of two key components: the desired AC-PC line and the midplane (Minoshima et al., 1994). Advances in image transformation techniques now allow the AC-PC line and midplane to be defined by placing points on relevant anatomical structures, such as the AC, the PC, and various midline structures. However, identifying these landmarks demands considerable expertise in brain anatomy and MRI, and this process can be challenging without high-resolution images (Bhanu Prakash et al., 2006; Evans et al., 1992). The potential for discrepancies in landmark identification remains high, even among experts, underscoring the complexity of accurate AC-PC realignment (Pallavaram et al., 2008).

There have been few attempts to develop tools that can automatically realign the MR image to the Schaltenbrand AC-PC plane (Ardekani & Bachman, 2009; Horn & Kuhn, 2015; Horn et al., 2017; Oxenford et al., 2022). However, perhaps surprisingly, there is currently no clear guideline to validate the results of these automated methods or to realign the image manually in case automated methods fail. Hence a standard guideline on how to identify these anatomical structures on structural MR images for manual AC-PC realignment and validate the realignment is an important gap in the literature. Here, therefore, we developed a protocol for the identification of the anatomical landmarks in T1-weighted MR data that are necessary for realigning structural MR images to the AC-PC plane, and for verifying the results of the ensuing AC-PC re-alignment. This proposed guideline has been developed to satisfy key features necessary for accurate data alignment:

1. The guideline adheres to the clinical version of the Schaltenbrand atlas (Fig. 1D), which is widely used in stereotactic and functional neurosurgery to localize subcortical structures (Horn et al., 2017). However, the guideline is still compatible with other AC-PC definitions, including the original AC-PC definitions by Talairach and Schaltenbrand (Fig. 1B and 1C).
2. The guideline does not assume prior anatomical knowledge nor experience with T1-weighted MR imaging and/or AC-PC realignment. These assumptions hold under extreme image rotations, across age groups, and for patients with Parkinson’s Disease.
3. The guideline was developed and validated on commonly used high-resolution structural contrast T1-weighted MR images but can also be used for other structural imaging contrasts such as T2-weighted MR image.
4. The guideline incorporates the results of a validation experiment involving both experts and non-experts.

In the following sections, we first present the proposed final version of the guideline in Section 2. This is followed by a comprehensive justification of key design choices in Section 3. Section 4 provides a concise overview of the evaluation process of the precursor guideline and the results. Finally, in Section 5, we offer a broader discussion, highlighting the practical benefits of the guideline and reviewing past efforts in similar directions

## 2. The Guideline

The proposed protocol is structured into three sections. In the first section, we provide a brief overview of the imaging anatomy relevant to the proposed landmarks. The second section presents a step-by-step guide for selecting the AC-PC points, which define the AC-PC line, as well as the four midline landmarks required for midplane definition. Finally, the third section outlines three visual criteria to verify the successful realignment of the AC-PC space.

### 2.1 Anatomical Planes

Three anatomical planes, which run perpendicular to each other along the X, Y, and Z axes (Fig. 2), are essential for studying the brain in all three orientations:

**Figure 2.**
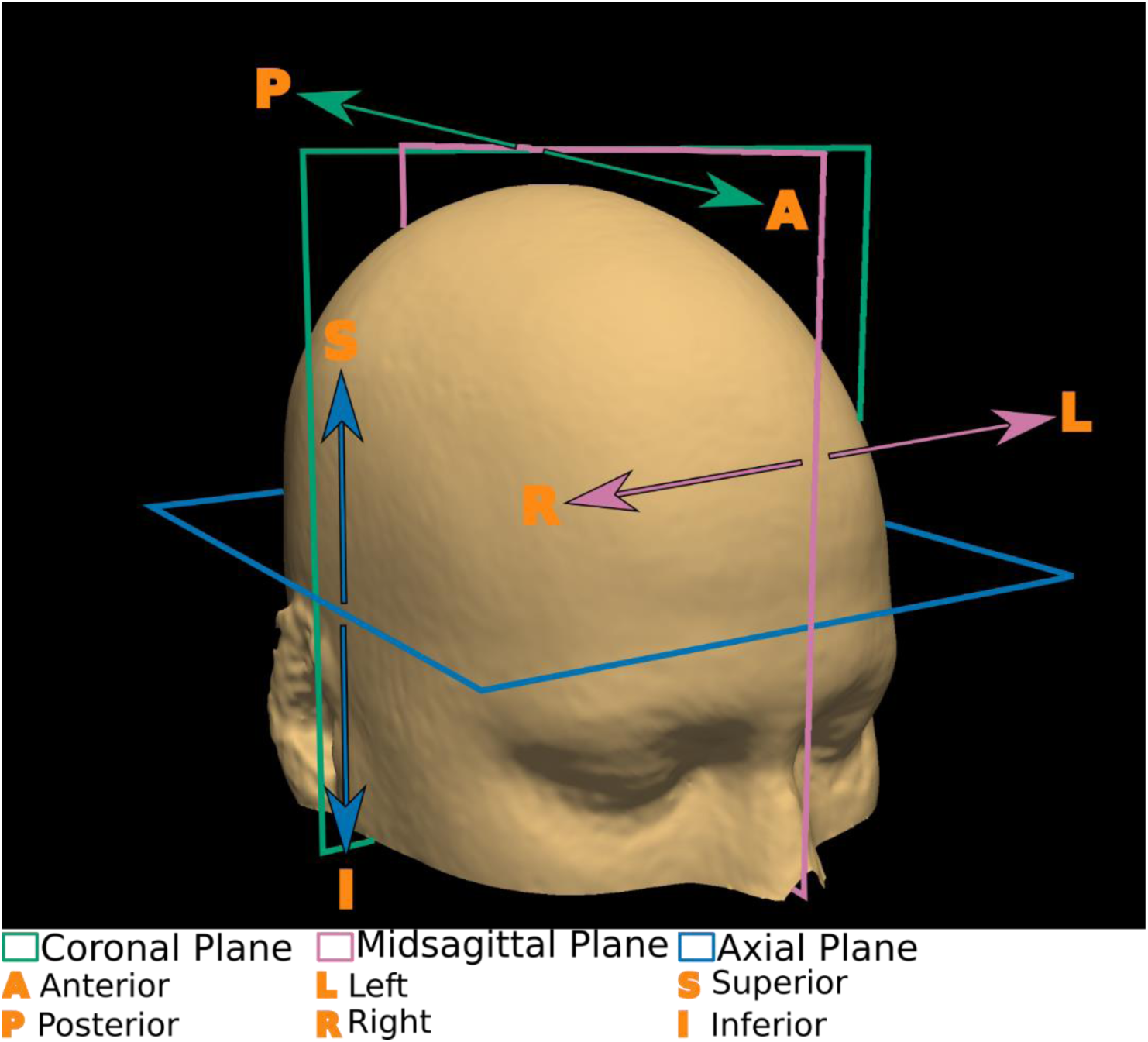
Three-dimensional illustration demonstrating all three cardinal planes.

1. **Midsagittal Plane**: This plane runs along the midline, dividing the brain into two halves—right and left (Fig. 3).

**Figure 3.**
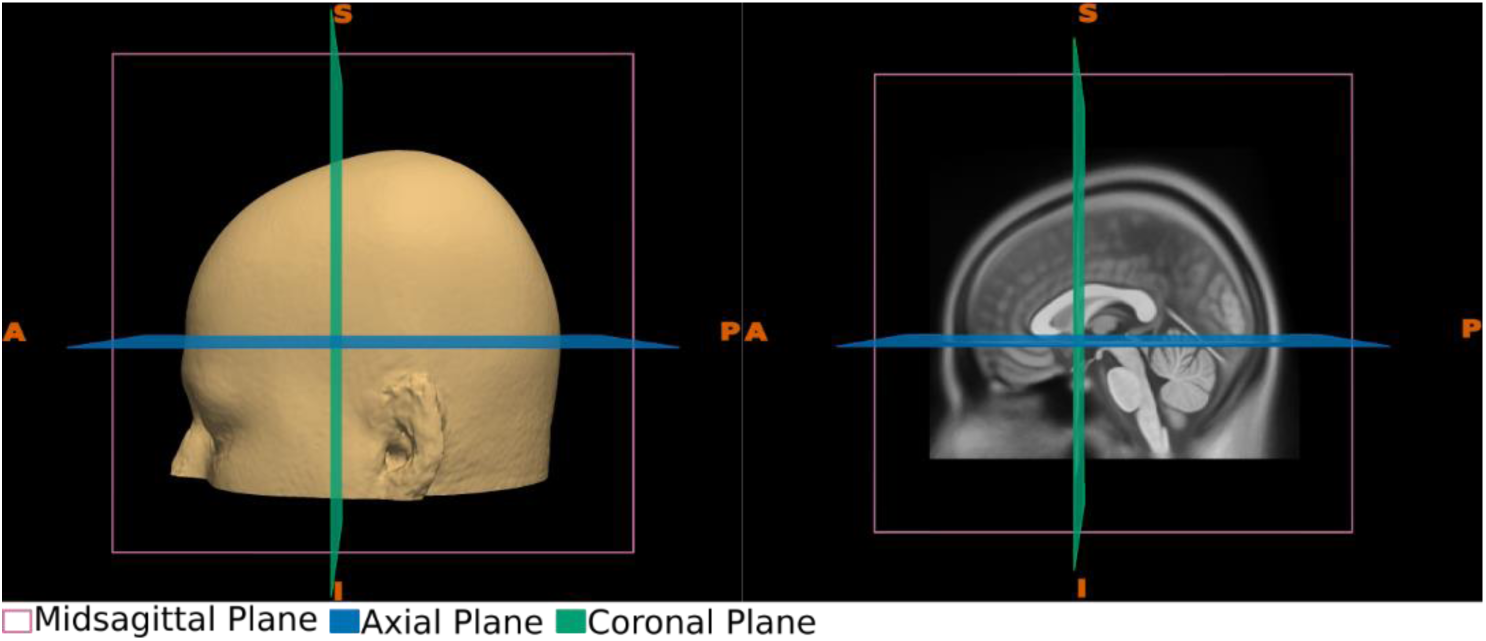
Sagittal plane views. The left panel illustrates a 3D view, while the right panel displays the midsagittal view of a standard MR image. Both panels also depict the coronal and axial planes.

2. **Coronal Plane**: The coronal plane divides the brain into front (anterior) and back (posterior) sections (Fig. 4).

**Figure 4.**
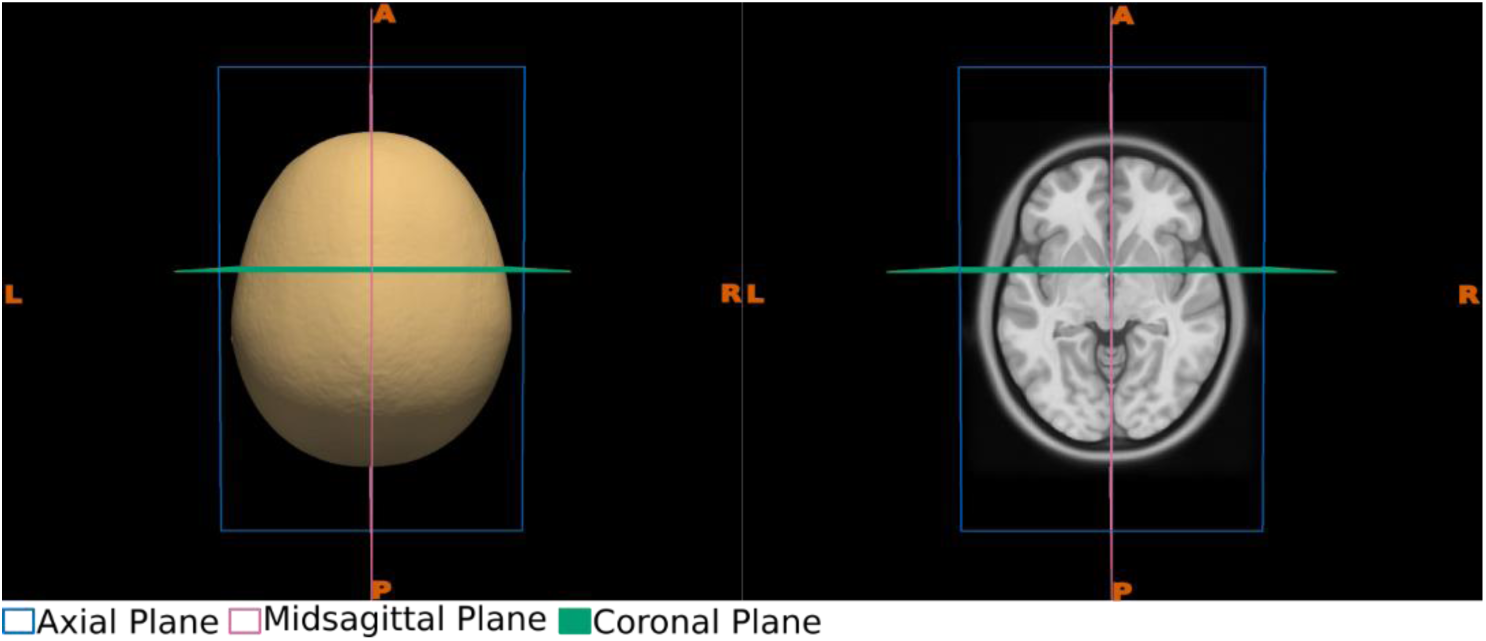
Axial plane views. The left panel illustrates a 3D view, while the right panel displays the axial view of a standard MR image. Both panels also depict the coronal and Sagittal planes.

3. **Axial (Transverse) Plane**: The axial plane divides the brain into upper (superior) and lower (inferior) parts, perpendicular to the longitudinal axis of the body (Fig. 5).

**Figure 5.**
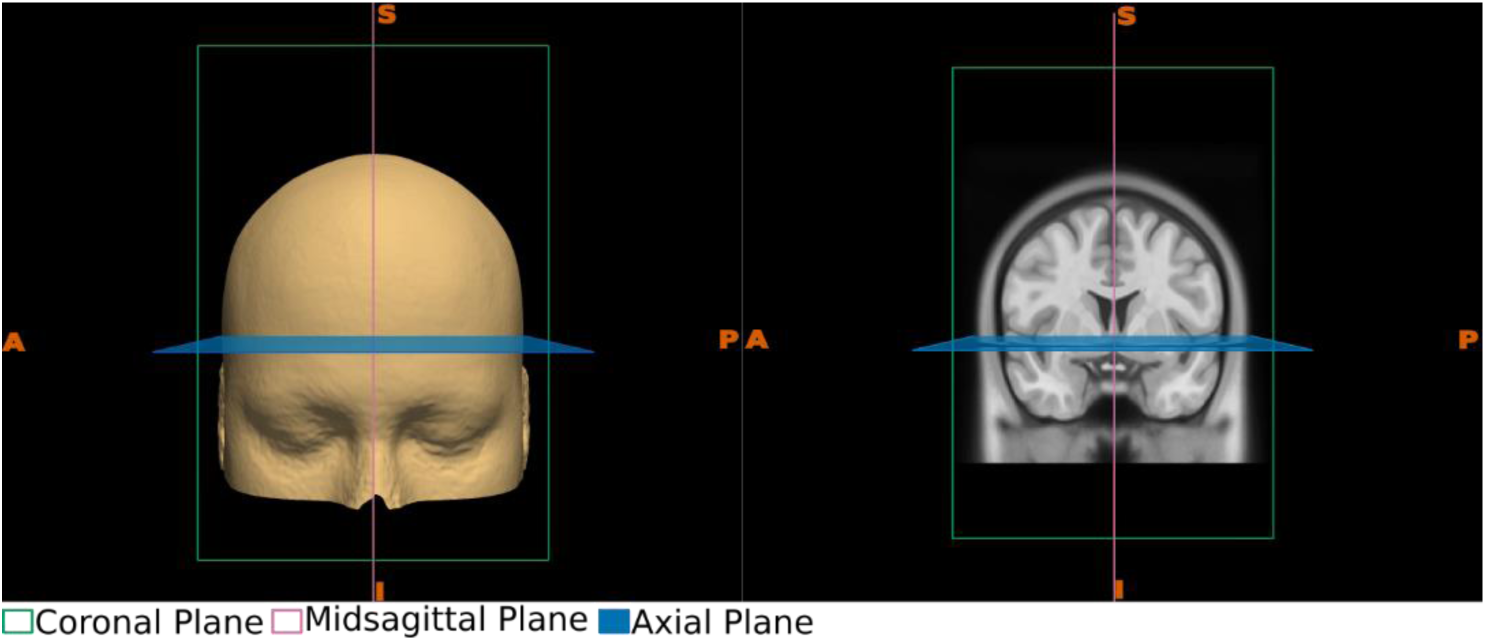
Coronal plane views. The left panel illustrates a 3D view, while the right panel displays the coronal view of a standard MR image. Both panels also depict the axial and Sagittal planes.

### 2.2 Anatomical Landmarks

In this section, we provide a brief overview of the anatomical landmarks used in the guideline, along with instructions on identifying these structures across all three cardinal planes.

#### 2.2.1. Anterior Commissure (AC)

The anterior commissure (AC) is a compact bundle of myelinated fibers that connects various regions across both hemispheres of the brain (Datta, 2013; Standring & Gray, 2008). Due to its consistent and well-defined location, the AC serves as a major landmark for reference in both neurosurgical and radiological practices (Schaltenbrand et al., 1977; Seeger & Zentner, 2002; Talairach & Tournoux, 1988)

##### Mid-sagittal

In the mid-sagittal plane, the AC can be identified as a small protrusion along the anterior wall of the third ventricle (Naidich et al., 1986; Standring & Gray, 2008). It often appears as an extension of the fornix in MRI images (Fig. 6).

**Figure 6.**
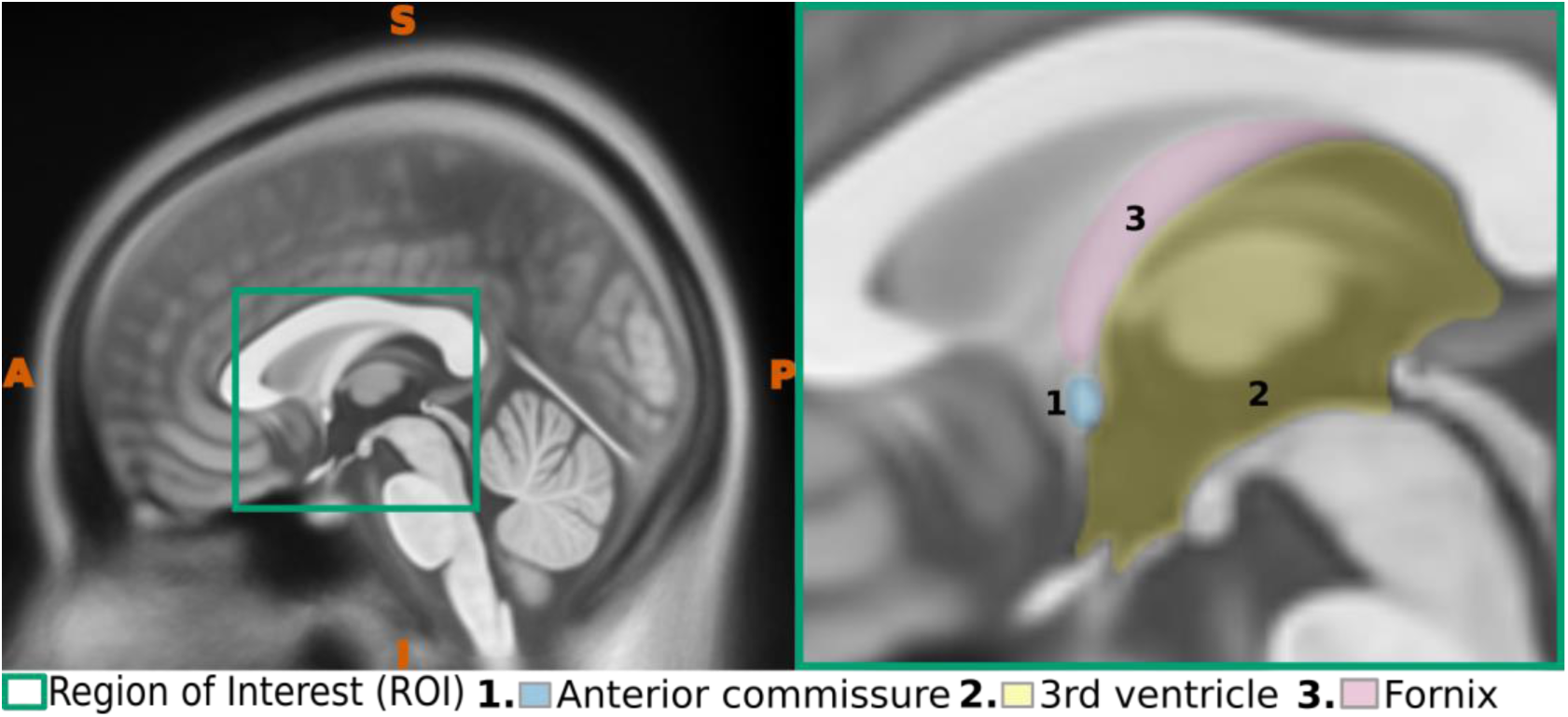
Sagittal view of the anterior commissure. The left panel presents a midsagittal view highlighting the region of interest (ROI), while the right panel offers an enlarged view of the ROI, emphasizing key anatomical structures.

##### Axial

In the axial plane, the AC can be identified in the midline as a thick band of white matter, appearing as a hyperintense structure on T1-weighted images (Fig. 7). It connects both hemispheres of the brain, forming the classic "handlebar" appearance (Naidich et al., 1986)

**Figure 7.**
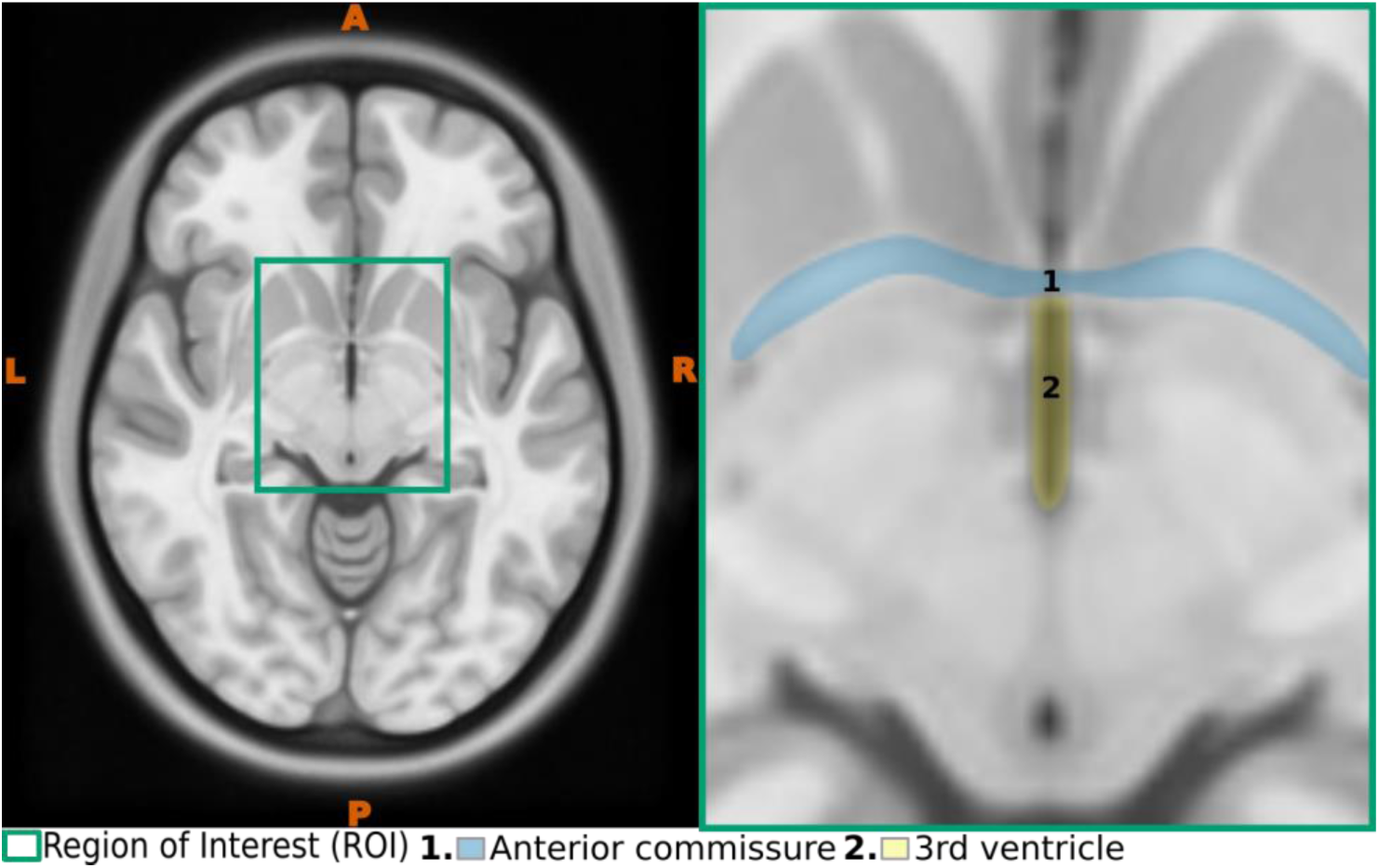
Axial view of the anterior commissure. The left panel displays an axial perspective with the region of interest (ROI) highlighted, while the right panel provides a magnified view of the ROI, emphasizing essential anatomical structures.

##### Coronal

In the coronal plane, AC can be identified as a thick white matter structure extends between the two hemispheres of the brain (Fig. 8), positioned between the lateral ventricles above and the third ventricle below (Naidich et al., 1986).

**Figure 8.**
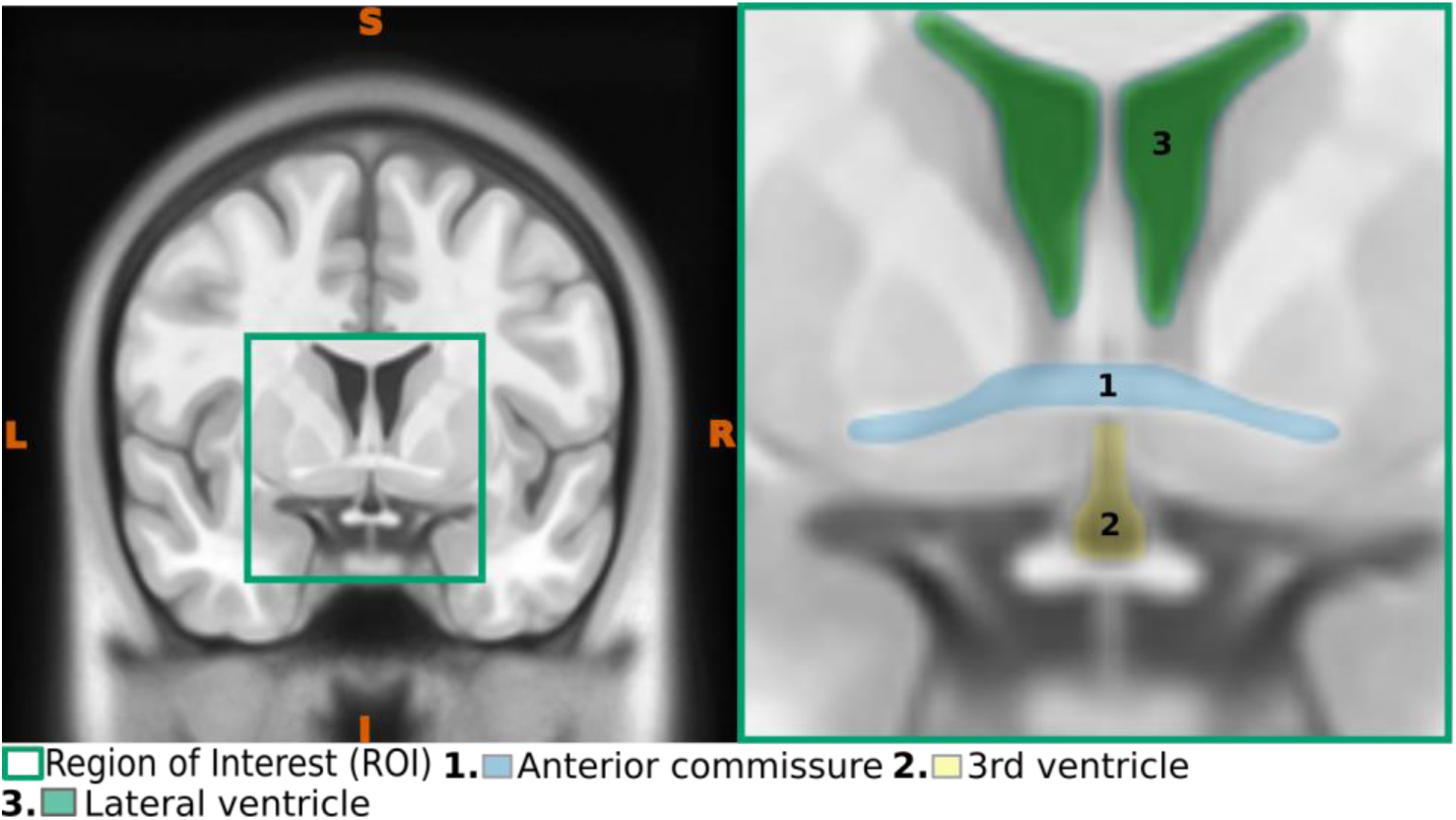
Coronal view of the anterior commissure. The left panel illustrates a coronal perspective highlighting the region of interest (ROI), while the right panel shows an enlarged, view of the ROI, emphasizing key anatomical features

#### 2.2.2. Posterior Commissure (PC)

The posterior commissure (PC) is a complex bundle of fibers that connects various structures in both cerebral hemispheres (Datta, 2013; Standring & Gray, 2008). It is one of the commonly used midline structures in neurosurgical references, owing to its consistent and well-defined location in the brain (Schaltenbrand et al., 1977; Seeger & Zentner, 2002; Talairach & Tournoux, 1988).

##### Mid-sagittal

The posterior commissure (PC) can be identified as a small, "C"-shaped structure located on the posterior wall of the third ventricle, just above the midbrain and at the beginning of the cerebral aqueduct (Fig. 9).

**Figure 9.**
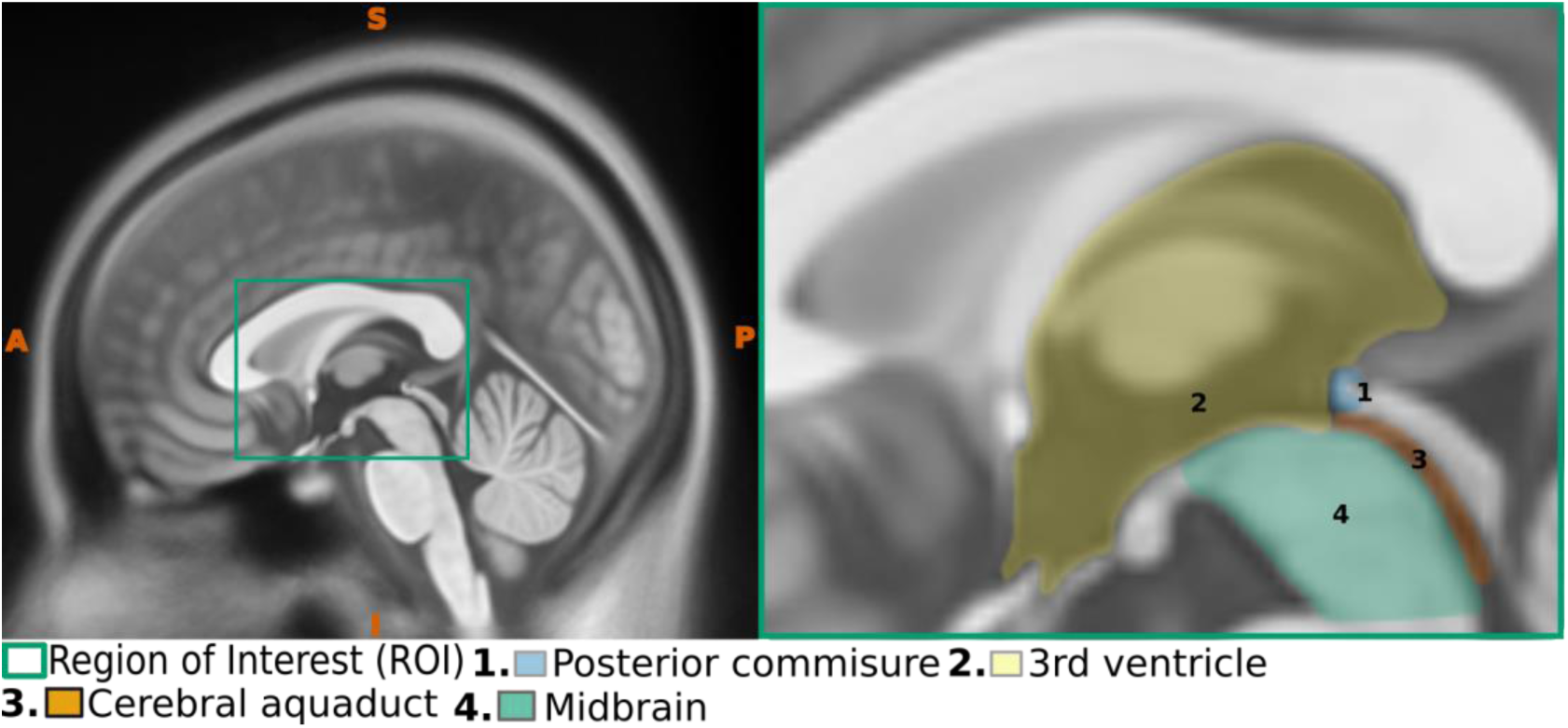
Sagittal view of the posterior commissure. The left panel presents a midsagittal view highlighting the region of interest (ROI), while the right panel provides an enlarged view of the ROI, emphasizing essential anatomical features.

##### Axial

A short band of hyperintense structure can be identified in the midline, connecting both cerebral hemispheres and contributing to the posterior boundary of the third ventricle (Fig. 10).

**Figure 10.**
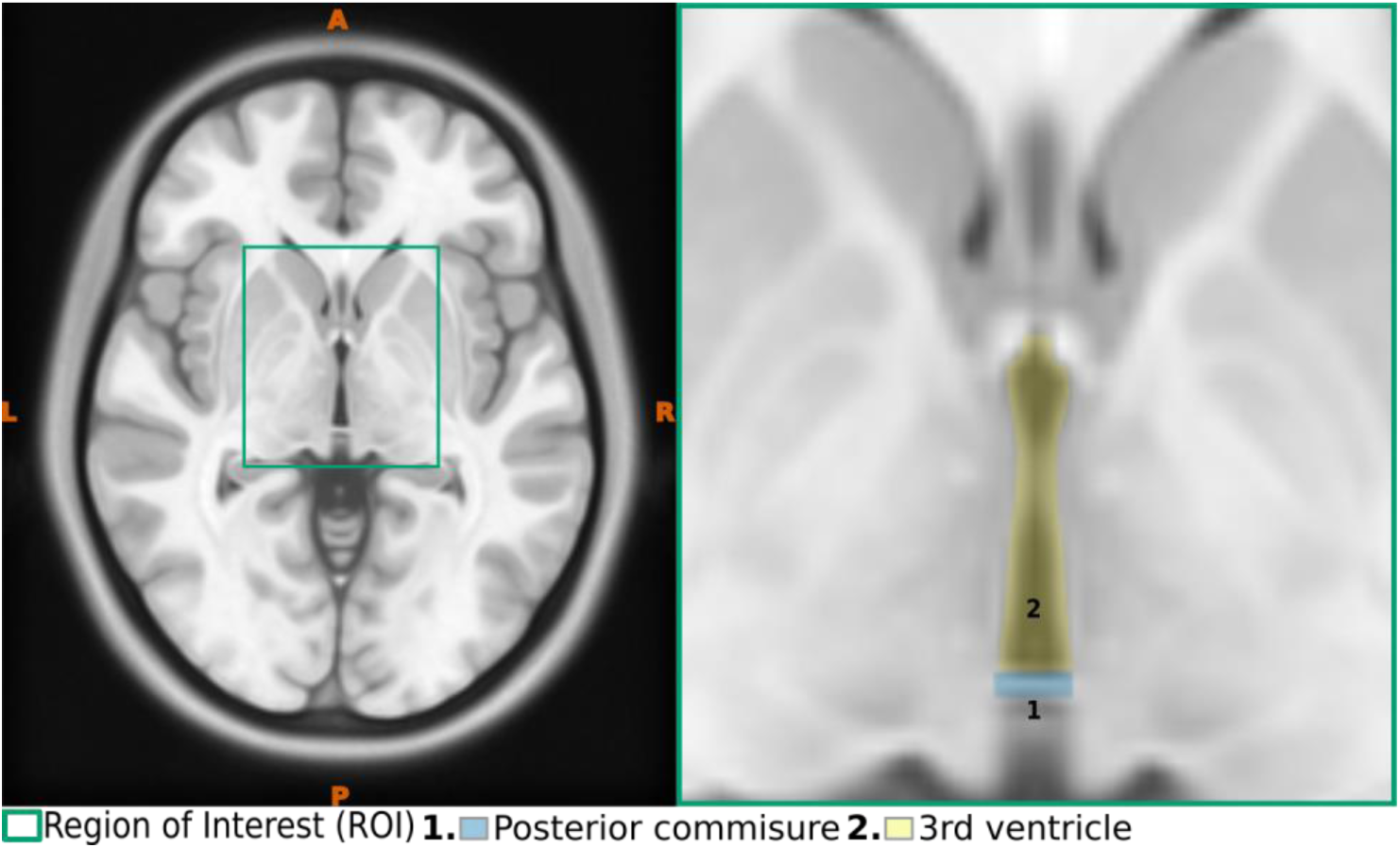
Axial view of the posterior commissure. The left panel presents a axial view highlighting the region of interest (ROI), while the right panel provides an enlarged view of the ROI, emphasizing essential anatomical features.

##### Coronal

PC appears as a short band in the midline, situated between the third ventricle above and the cerebral aqueduct below (Fig. 11).

**Figure 11.**
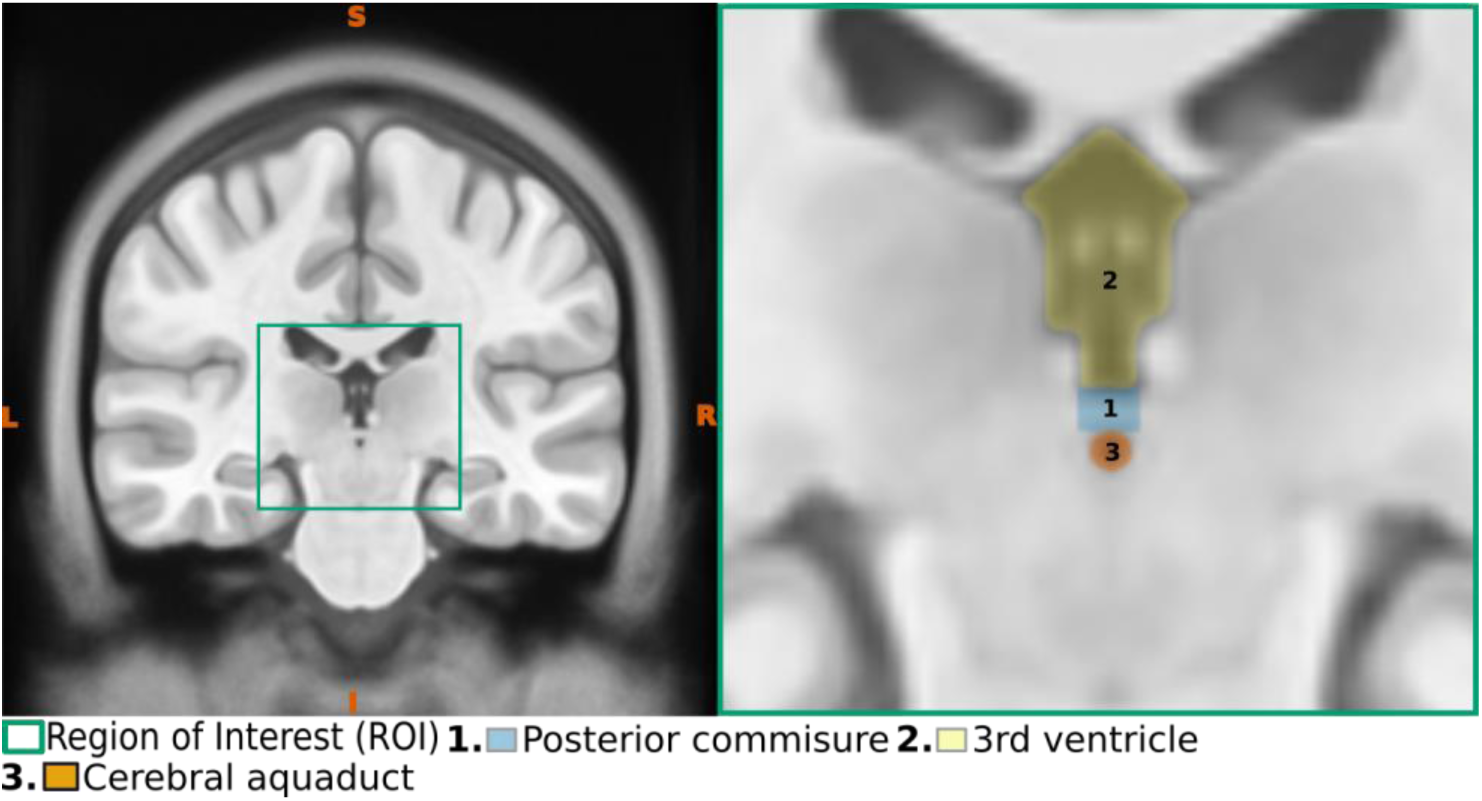
Coronal view of the posterior commissure. The left panel presents a coronal view highlighting the region of interest (ROI), while the right panel provides an enlarged view of the ROI, emphasizing essential anatomical features

#### 2.2.3. Superior Pontine notch

The superior pontine notch is a subtle indentation located along the superior margin of the pons, visible in the sagittal plane. It serves as a key anatomical landmark, often used to demarcate the junction between the pons and the midbrain, also known as the pontomesencephalic junction (Ardekani & Bachman, 2009; Lau et al., 2019), frequently serving as a key imaging landmark in radiological assessments (Oba et al., 2005)

##### Mid-sagittal

The superior pontine notch is easily identifiable in the mid-sagittal plane, situated at the junction between the midbrain above and the pons below, within the interpeduncular fossa (Fig. 12).

**Figure 12.**
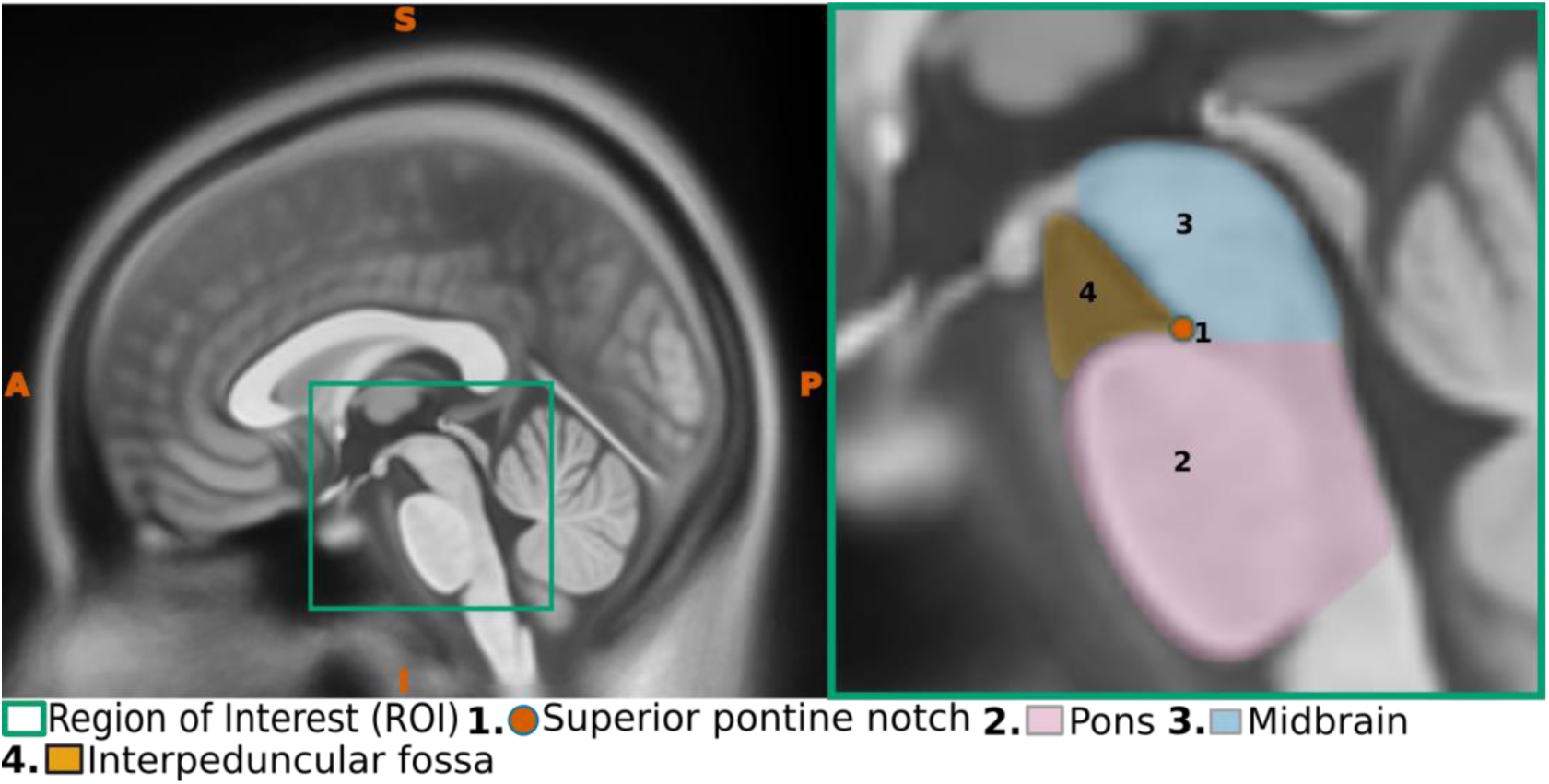
Sagittal view of the superior pontine notch. The left panel displays a midsagittal perspective highlighting the region of interest (ROI), while the right panel provides a magnified view of the ROI, emphasizing critical anatomical structures.

##### Axial

In the corresponding axial slice, the superior pontine notch can be found at the midline, located at the junction where the right and left cerebral peduncles meet (Fig. 13).

**Figure 13.**
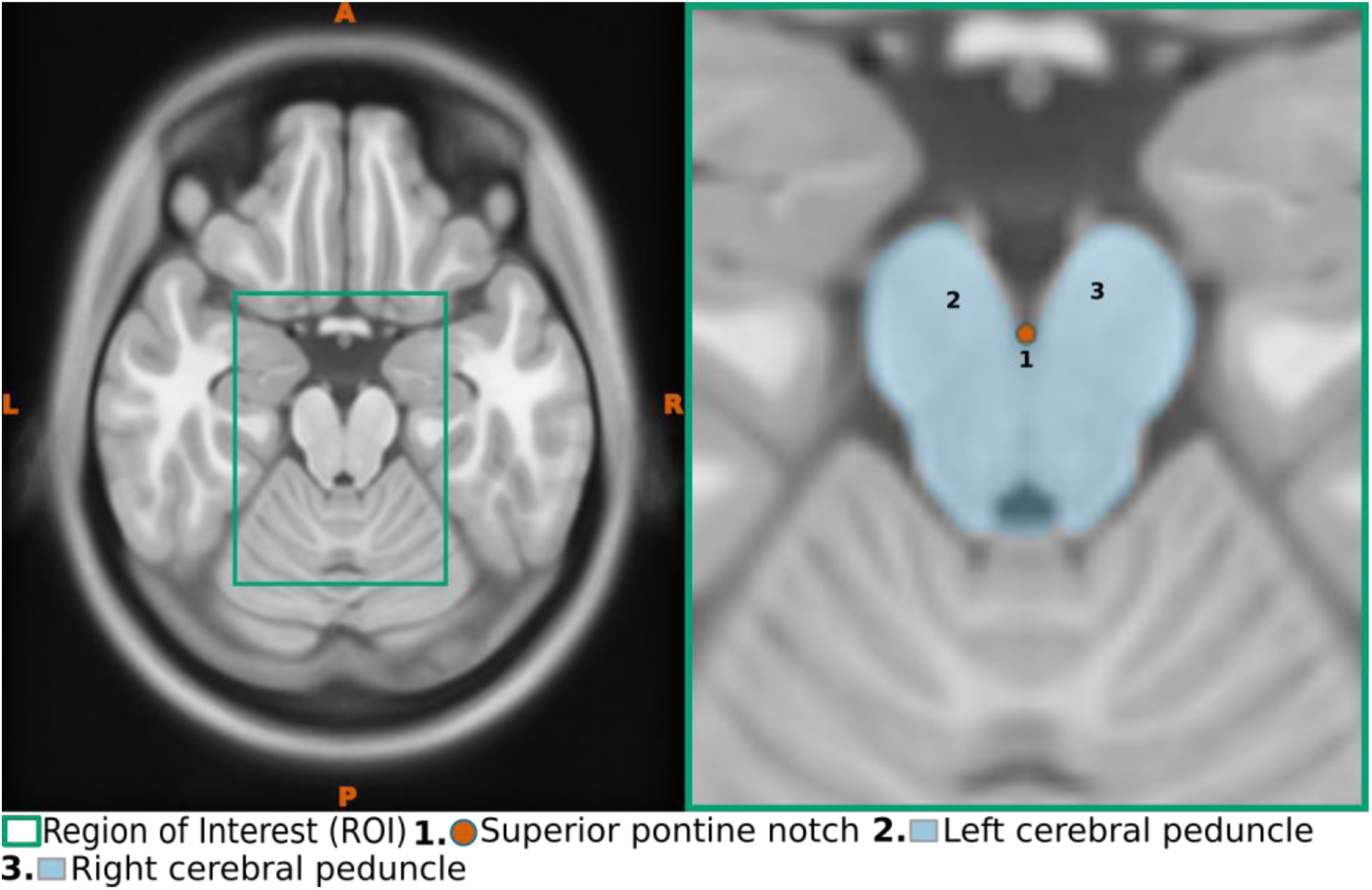
Axial view of the superior pontine notch. The left panel displays a axial view highlighting the region of interest (ROI), while the right panel provides a magnified view of the ROI, emphasizing critical anatomical structures.

##### Coronal

In the corresponding coronal slice, the superior pontine notch appears as a small midline opening at the pontomesencephalic junction, marking the transition zone between the midbrain above and the pons below (Fig. 14).

**Figure 14.**
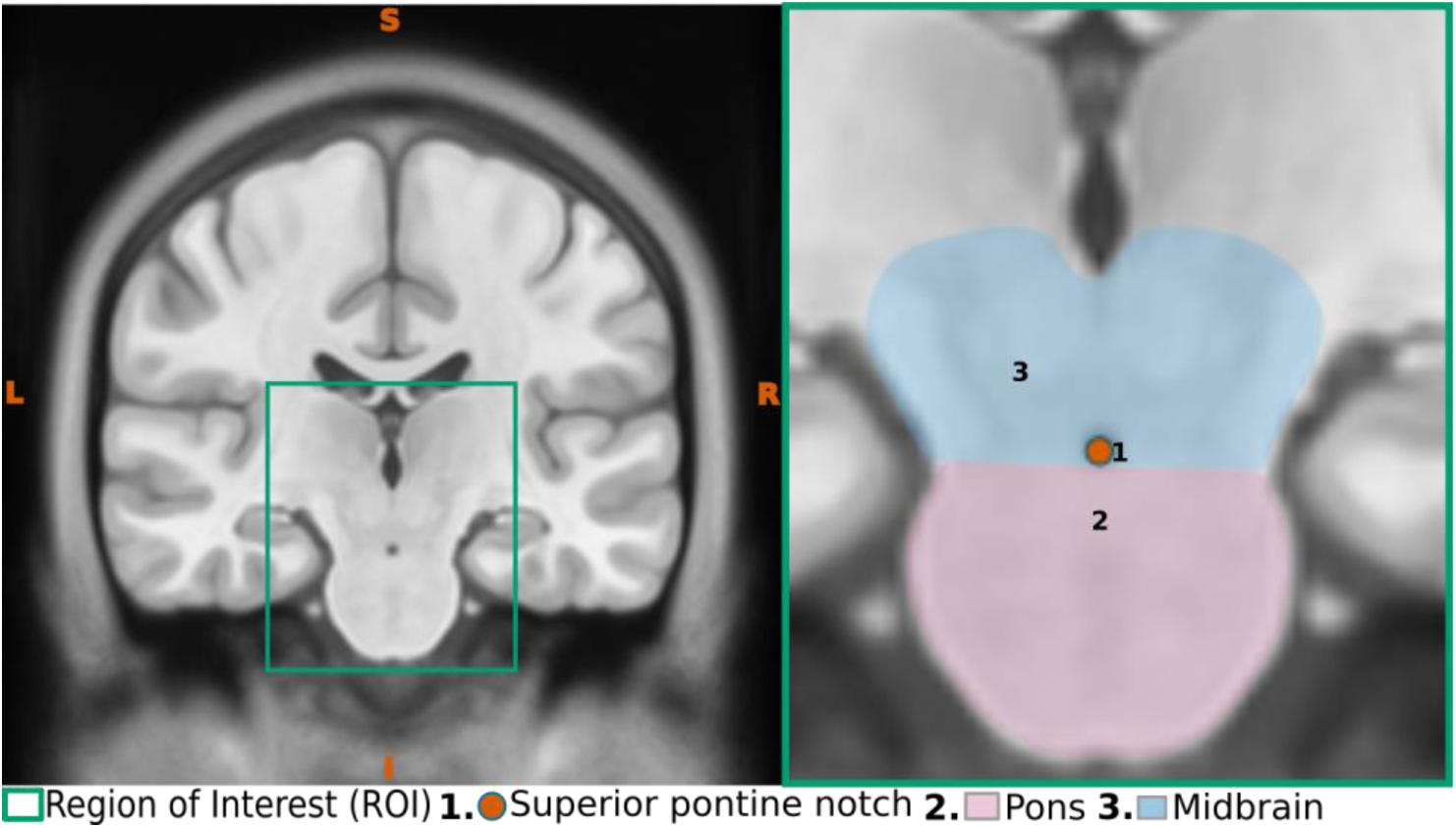
Coronal view of the superior pontine notch. The left panel displays a coronal view highlighting the region of interest (ROI), while the right panel provides a magnified view of the ROI, emphasizing critical anatomical structures.

#### 2.2.4 Fastigium or Apex of the roof of the Fourth Ventricle

The fastigium or fastigium cerebelli is the apex of the tent-shaped roof of the fourth ventricle in the brain (Matsushima et al., 1982; Salma et al., 2013). This apex marks the junction between the superior and inferior parts of the ventricular roof (Morris et al., 2022; Stratchko et al., 2016), serving as a key structural landmark for both the cerebellum and the brainstem.

##### Mid-sagittal

The fastigium is best visualized in the midsagittal plane, where the two parts of the roof of the fourth ventricle converge (Fig. 15). At this junction, it appears as a protrusion into the cerebellum, marking the point where the superior and inferior portions of the ventricular roof meet

**Figure 15.**
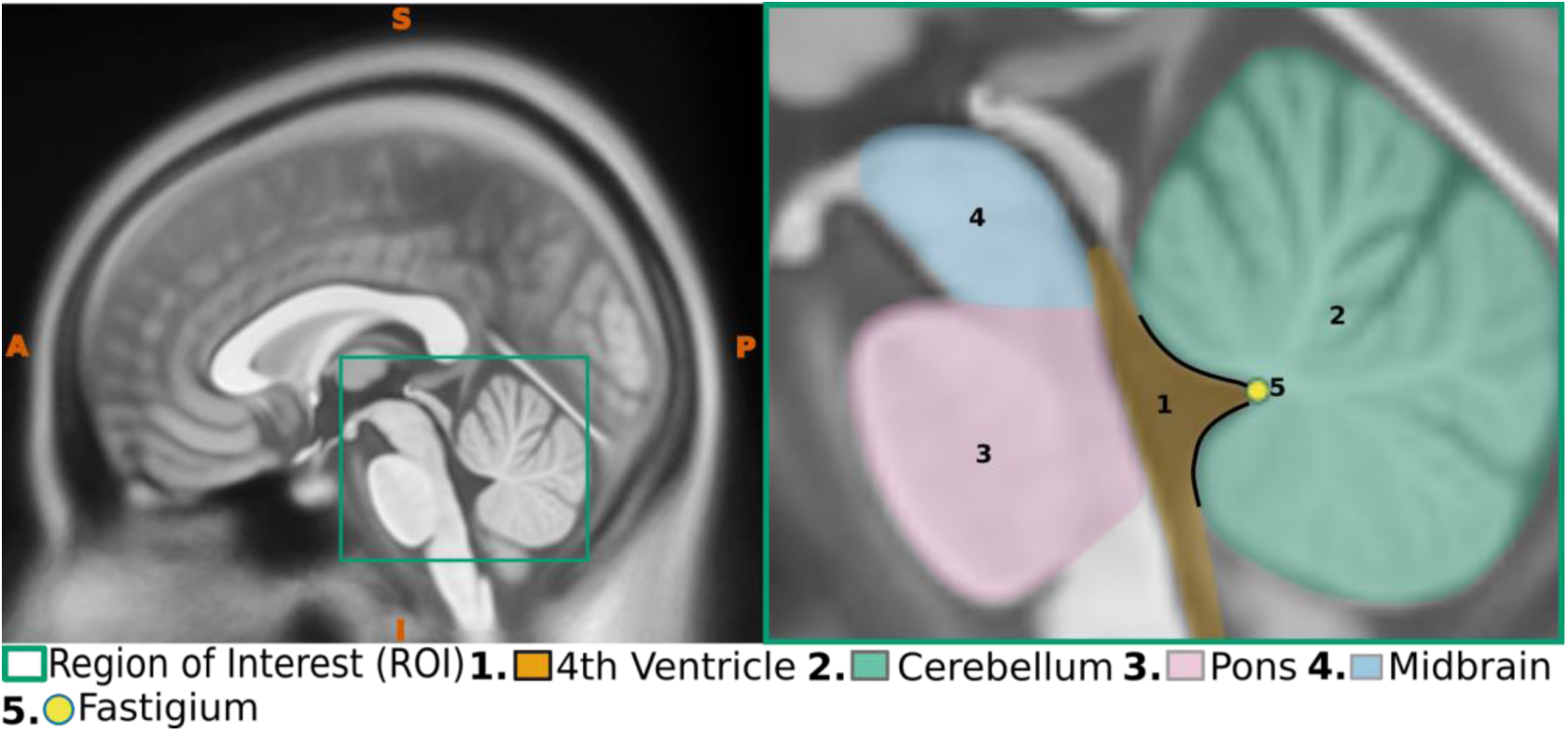
Sagittal view of the fastigium. The left panel illustrates a midsagittal view with the region of interest (ROI) highlighted, and the right panel shows an enlarged depiction of the ROI, emphasizing significant anatomical features.

##### Axial

In the corresponding axial slice, the fastigium appears as a flat surface, forming one of the key boundaries of the fourth ventricle (Fig. 16).

**Figure 16.**
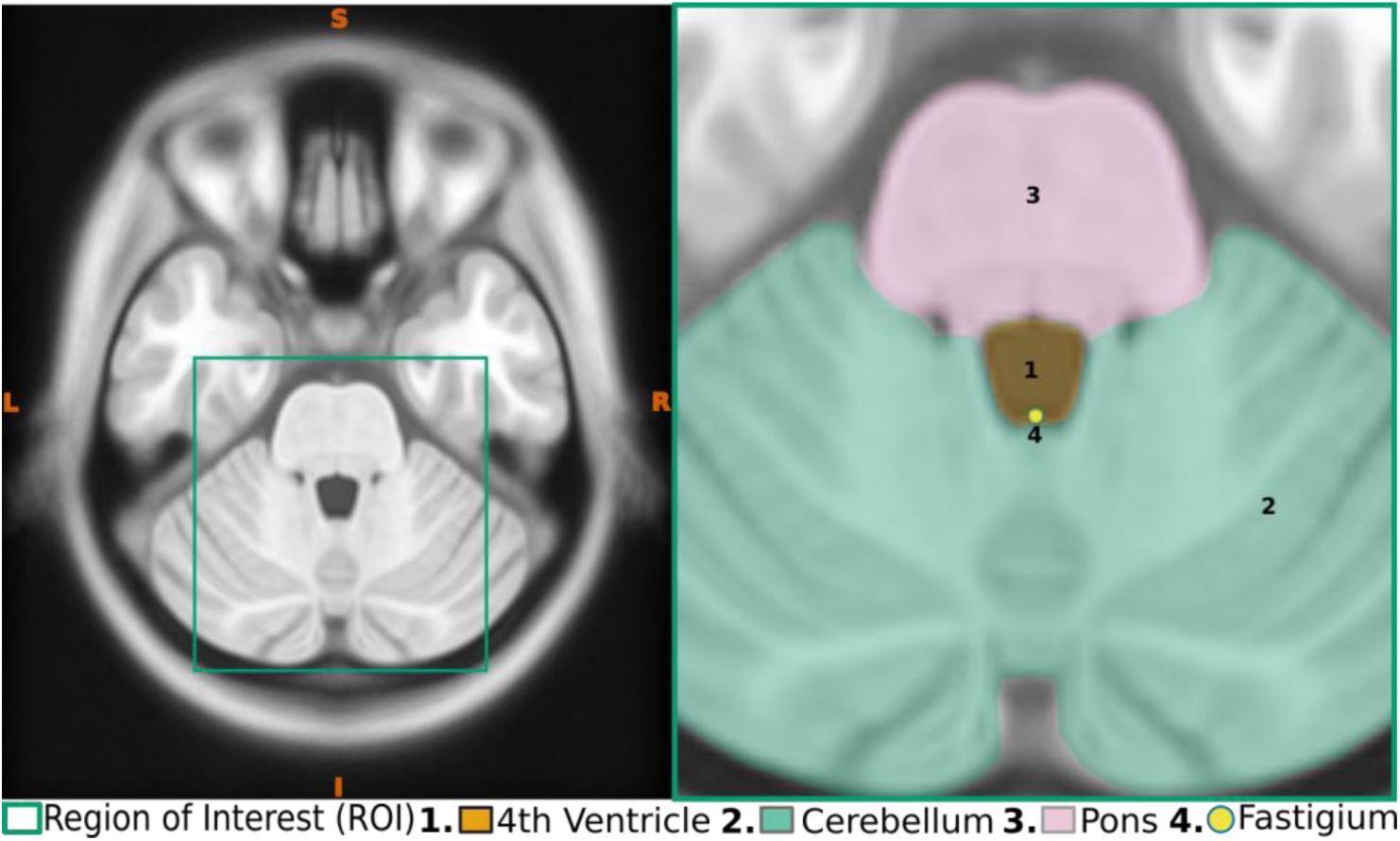
Axial view of the fastigium. The left panel illustrates a axial view with the region of interest (ROI) highlighted, and the right panel shows an enlarged depiction of the ROI, emphasizing significant anatomical features.

##### Coronal

In the corresponding coronal slice, the fastigium appears at the center of the cerebellum, presenting as a curved apex of the roof of the fourth ventricle (Fig. 17).

**Figure 17.**
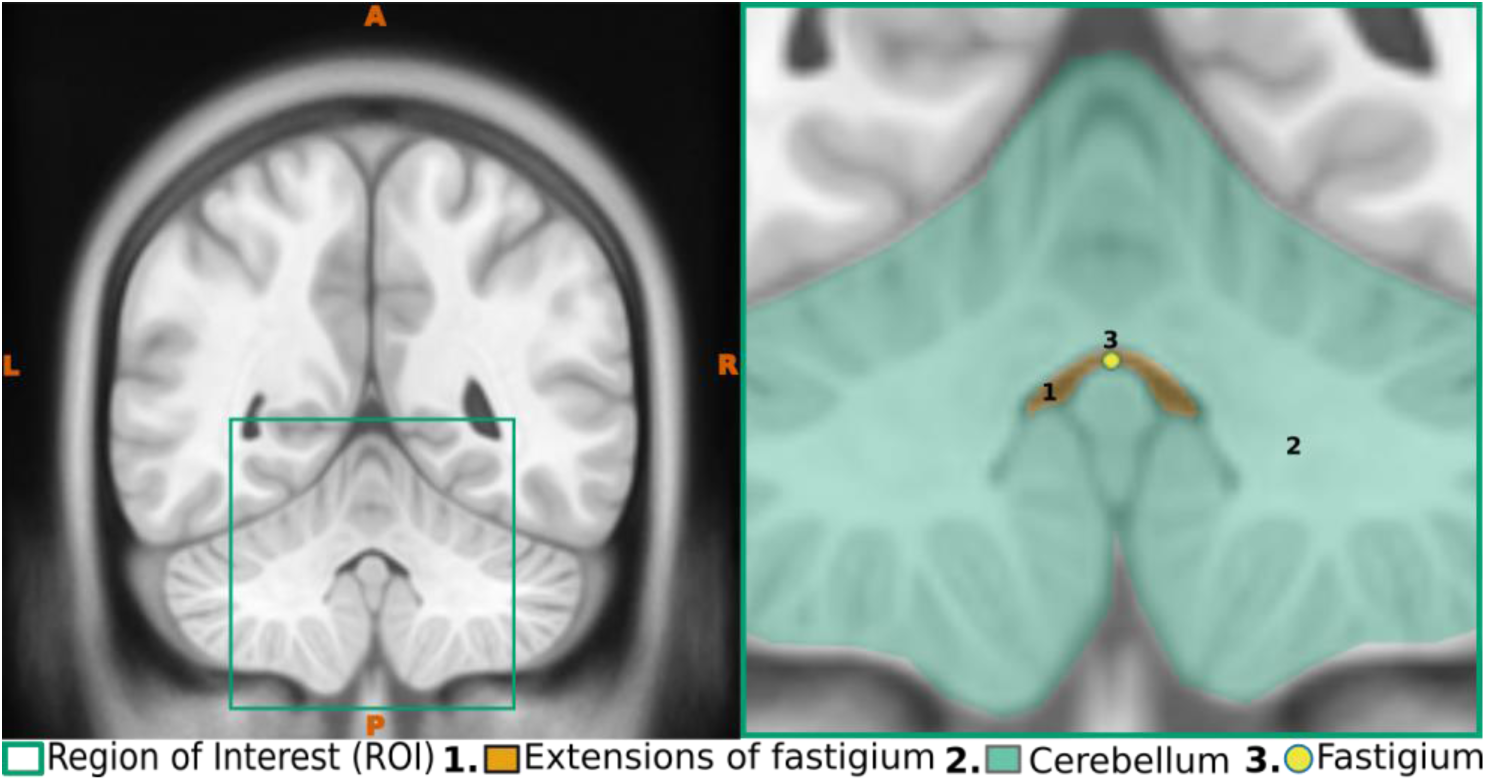
Coronal view of the fastigium. The left panel illustrates a coronal view with the region of interest (ROI) highlighted, and the right panel shows an enlarged depiction of the ROI, emphasizing significant anatomical features.

#### 2.2.5. Corpus callosum (CC)

The corpus callosum is the largest fibre pathway in the brain, linking the majority of the cerebral cortices of both hemispheres (Datta, 2013; Standring & Gray, 2008). It forms a significant portion of the roof of the lateral ventricles. Structurally, the corpus callosum is divided into five distinct parts, with the central and largest portion referred to as the trunk (Musiek, 1986). This trunk can be readily identified across all three imaging planes. usually located in the middle of the brain.

##### Mid-Sagittal

The entire length of the corpus callosum is best visualized in the mid-sagittal plane. On the flat medial surface of each hemisphere, a clearly visible, large, arched band with an upward convexity can be recognized, appearing hyperintense on T1-weighted images (Fig. 18).

**Figure 18.**
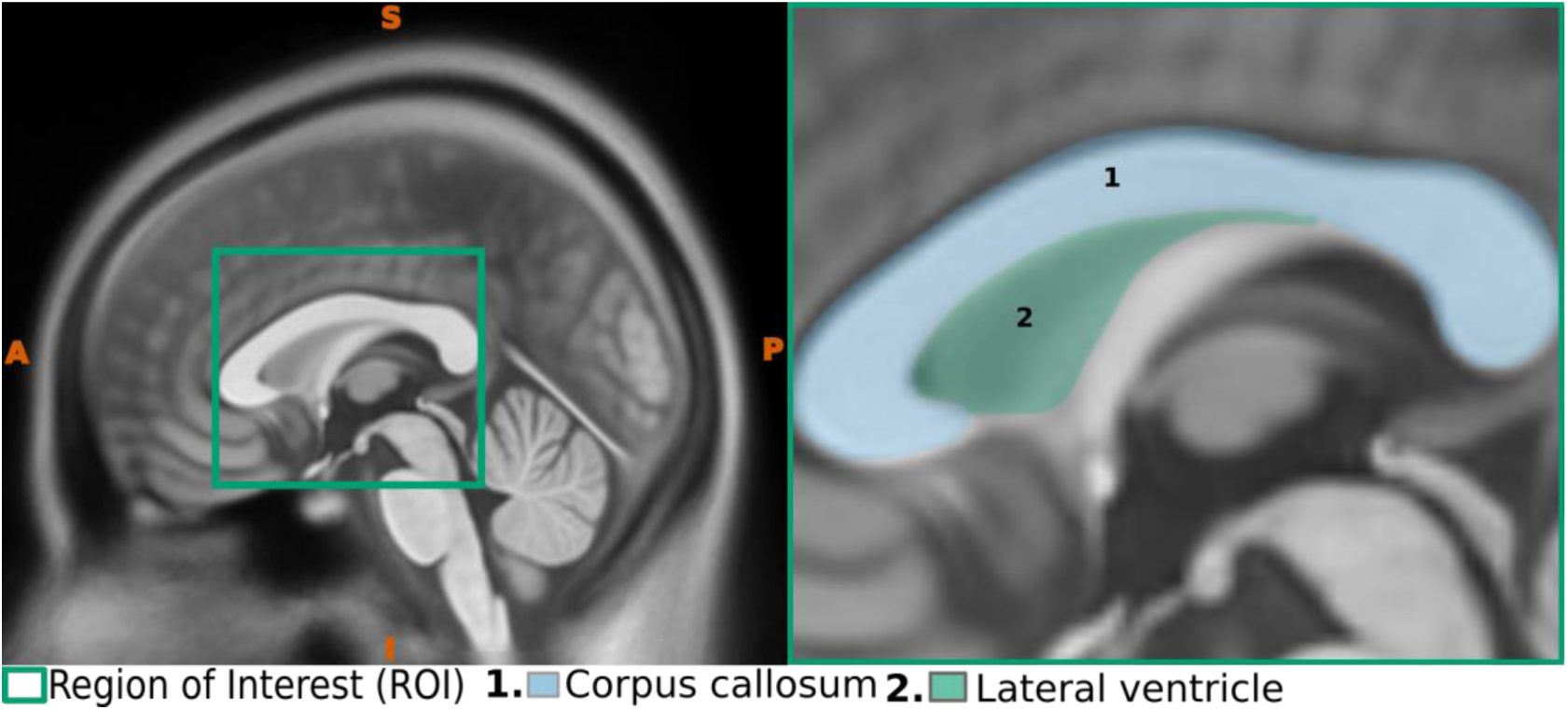
Sagittal view of the Corpus callosum. The left panel illustrates a midsagittal view with the region of interest (ROI) highlighted, and the right panel shows an enlarged depiction of the ROI, emphasizing significant anatomical features

##### Axial

In the axial plane, the middle two-thirds of the corpus callosum appear as a large X-shaped band of white matter in the midline, effectively connecting both hemispheres (Fig. 19).

**Figure 19.**
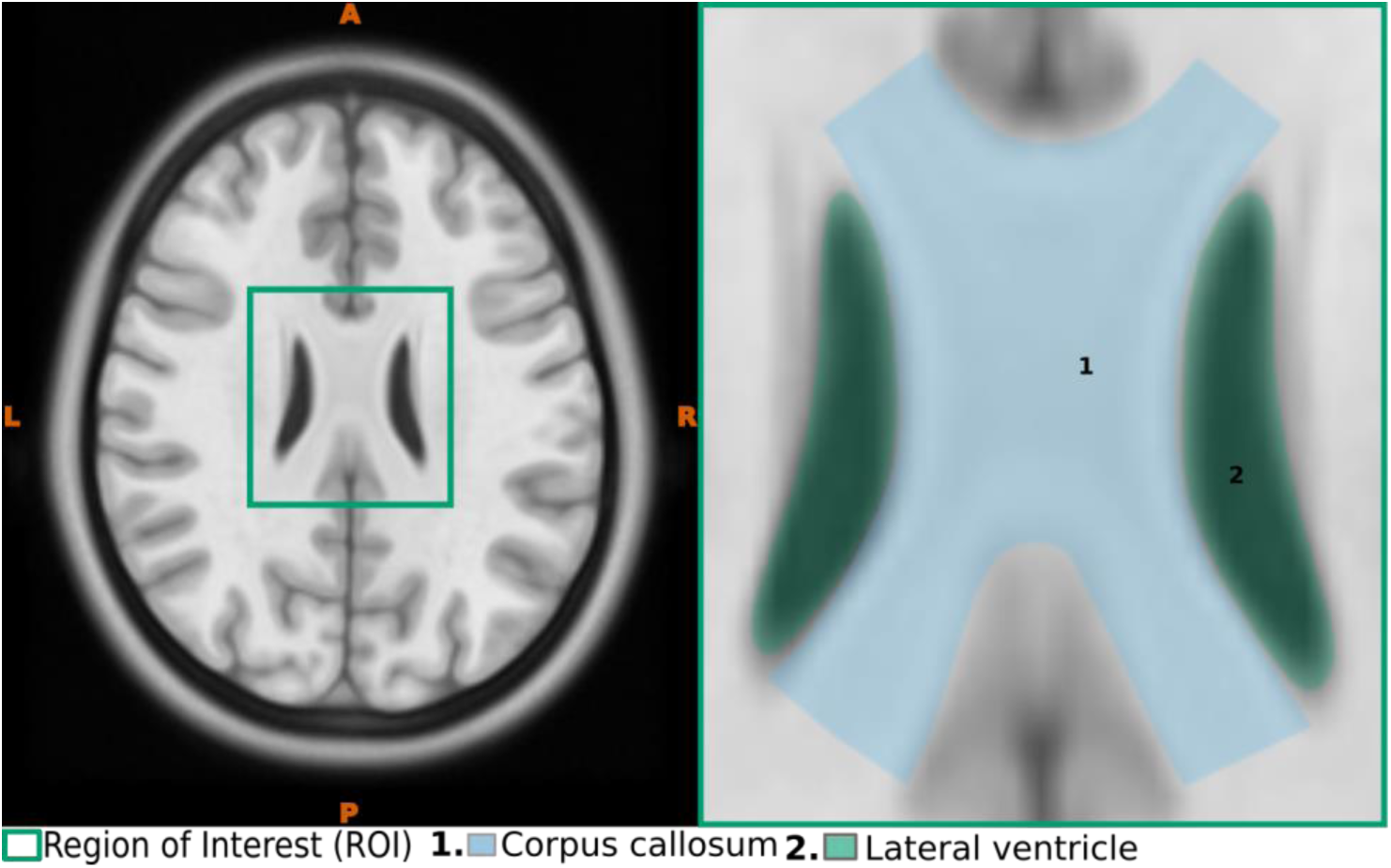
Axial view of the Corpus callosum. The left panel illustrates a axial view with the region of interest (ROI) highlighted, and the right panel shows an enlarged depiction of the ROI, emphasizing significant anatomical features

##### Coronal

In the corresponding coronal plane, the trunk of the corpus callosum appears as a hammock-shaped bridge, extending into both hemispheres just above the lateral ventricles (Fig. 20).

**Figure 20.**
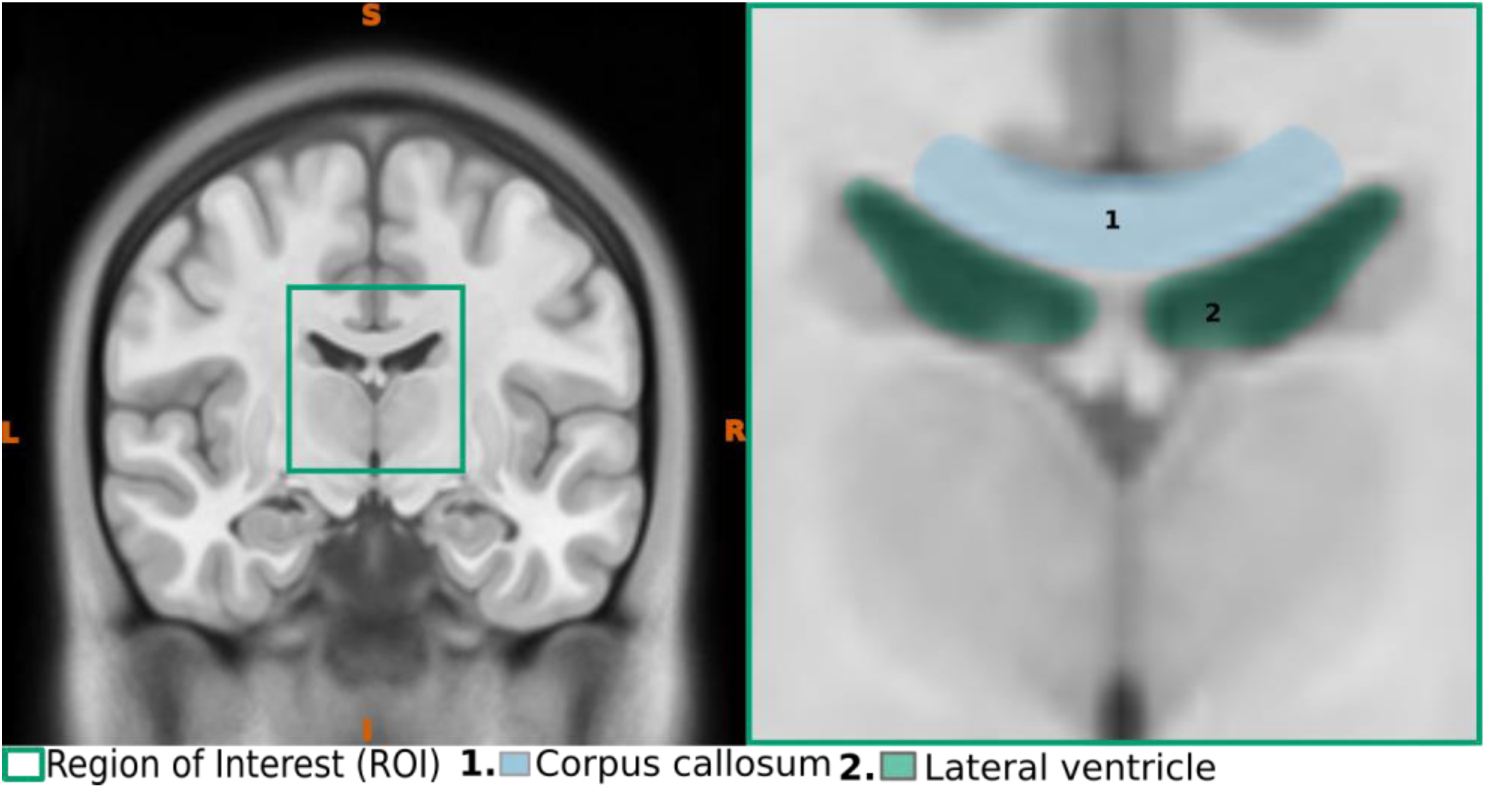
Coronal view of the Corpus callosum. The left panel illustrates a coronal view with the region of interest (ROI) highlighted, and the right panel shows an enlarged depiction of the ROI, emphasizing significant anatomical features

#### 2.2.6. Falx Cerebri

The falx cerebri is the largest infolding of the dura mater, the protective outer layer of the brain (Adeeb et al., 2012; Moore et al., 2014). This thin, crescent-shaped structure extends downward from the inner surface of the skull, running along the midline within the longitudinal fissure, which separates the brain into two cerebral hemispheres (Datta, 2013; Moore et al., 2014; Standring & Gray, 2008). In the sagittal view, the falx cerebri appears as a crescent-shaped structure (Standring & Gray, 2008). Along its upper convex margin runs the superior sagittal sinus, one of the largest venous channels in the brain (Datta, 2013; Moore et al., 2014; Standring & Gray, 2008). This close anatomical relationship between the falx cerebri and the superior sagittal sinus can be a useful landmark when tracing the falx cerebri in medical images

##### Mid-Sagittal

Identifying the falx cerebri in the sagittal plane requires considerable effort. In the midsagittal plane, it presents as a thin, translucent structure with a glassy appearance. This subtlety can make it challenging to trace the falx accurately. However, careful observation allows for identification by following the inferior border of the superior sagittal sinus, which presents as a distinct hyperintense arch on T1-weighted images (Fig. 21).

**Figure 21.**
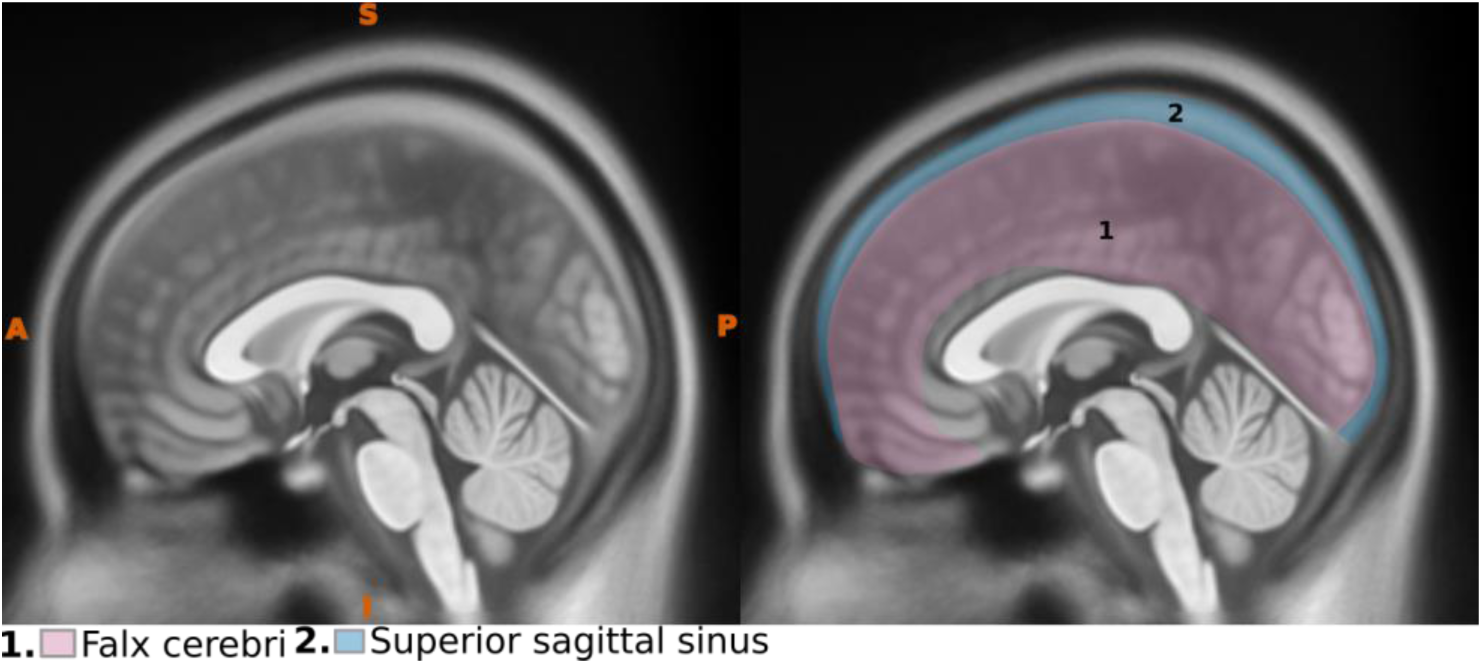
Sagittal depiction of the falx cerebri. The left panel provides a midsagittal view, while the right panel highlights significant anatomical features.

##### Axial

In the axial plane, the falx cerebri can be traced within the longitudinal fissure, extending from the borders of the superior sagittal sinus (Fig. 22). This sinus is characterized by a triangular hyperintense mass located at both the anterior and posterior ends of the brain in the midline (Adeeb et al., 2012; Standring & Gray, 2008), serving as a key landmark for identifying the falx cerebri.

**Figure 22.**
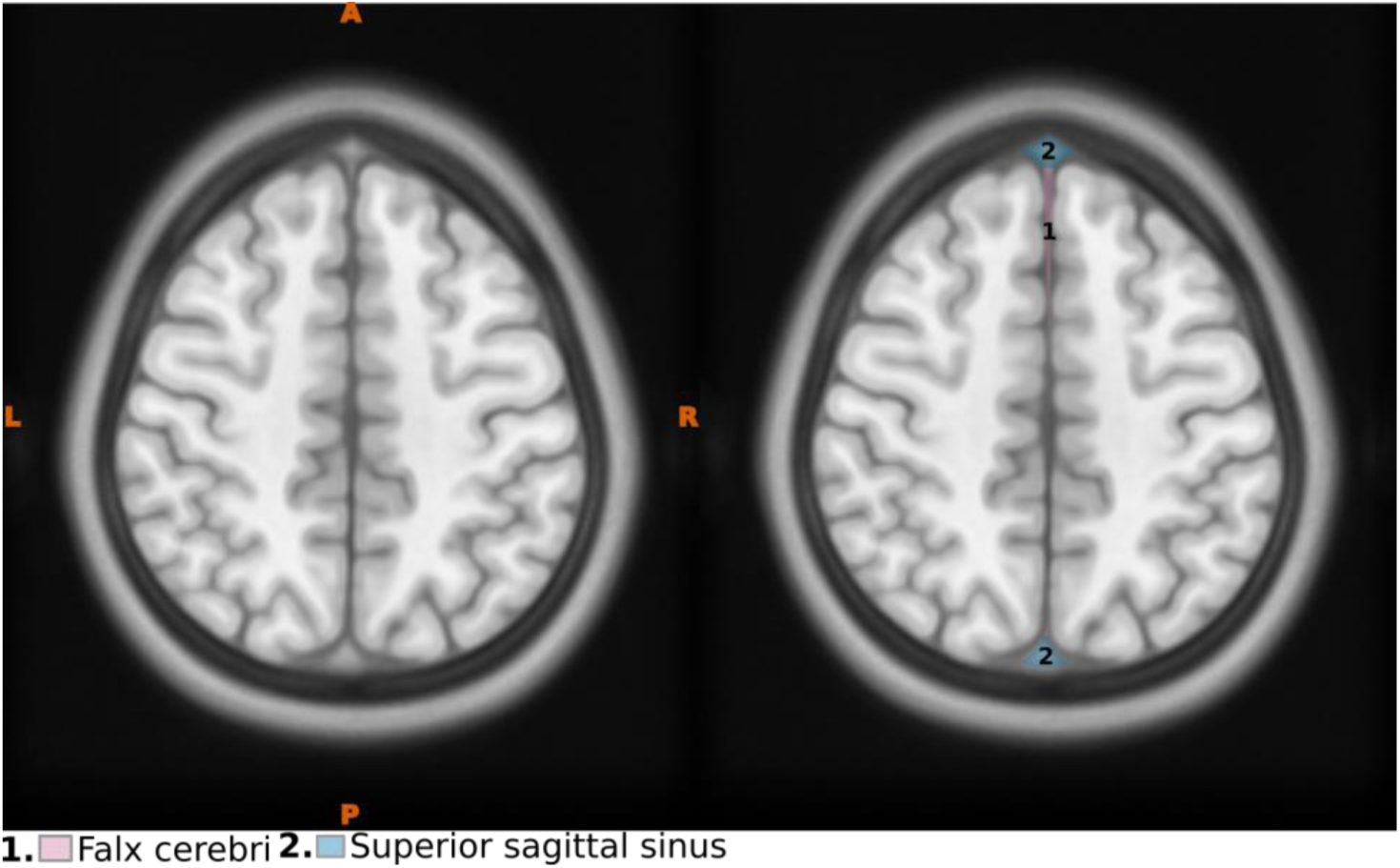
Axial depiction of the falx cerebri. The left panel provides a axial view, while the right panel highlights significant anatomical features

##### Coronal

In the coronal view, the falx cerebri can be recognized as a slender sheet extending from the triangular mass of the superior sagittal sinus in the midline, effectively separating the two hemispheres of the brain (Fig. 23).

**Figure 23.**
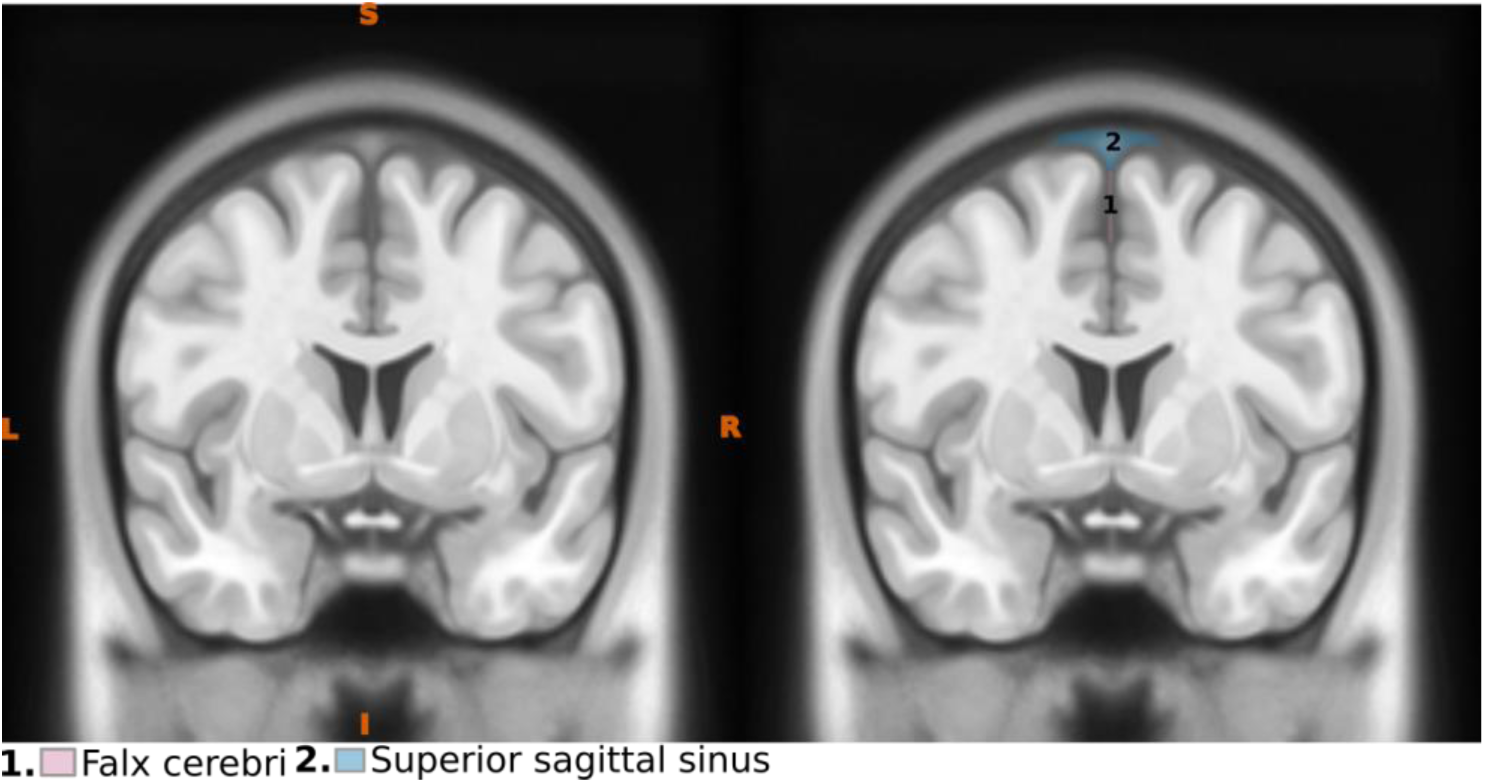
Coronal depiction of the falx cerebri. The left panel provides a coronal view, while the right panel highlights significant anatomical features

### 2.2 The Protocol

Here, we present our protocol in a structured, pseudo-algorithmic format with step-by-step instructions.

#### 2.2.1. Anterior Commissure (AC)

1: Identify the Anterior Commissure (AC; Fig. 24).

**Figure 24.**
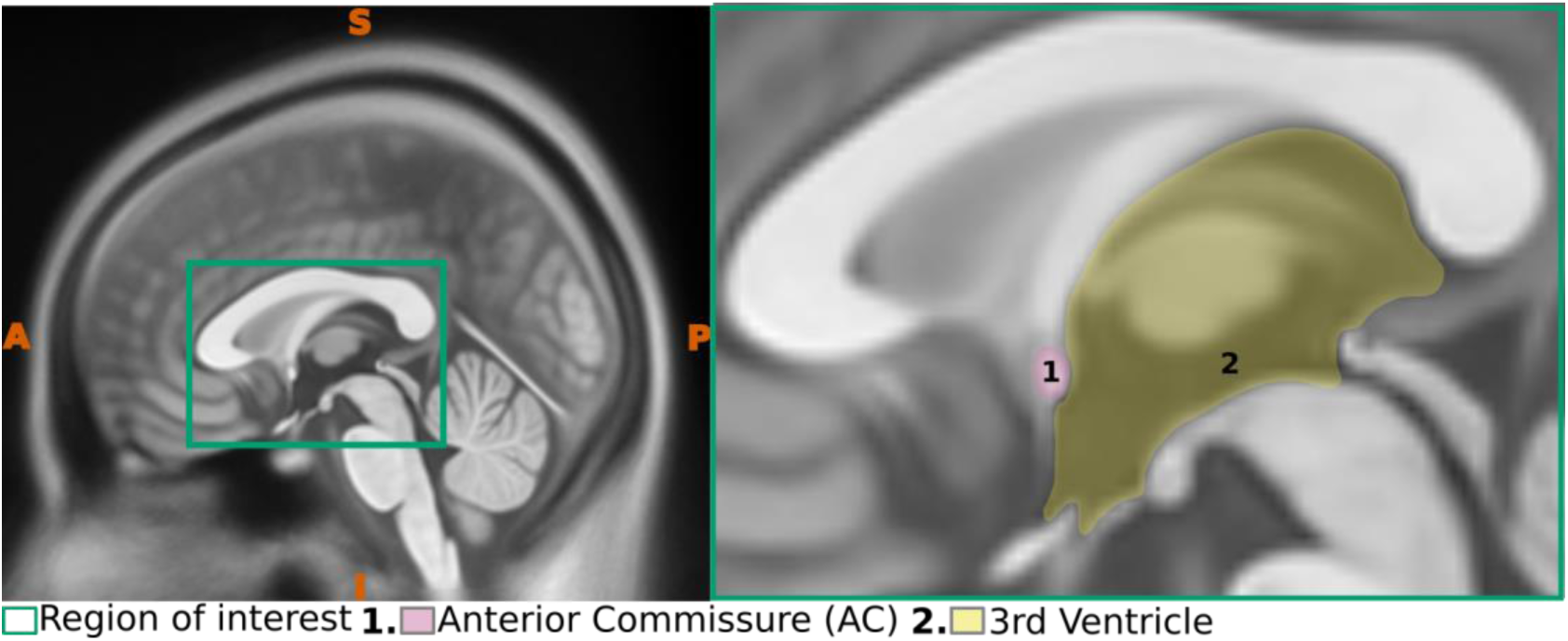
Sagittal view of the Anterior commissure. The left panel presents a midsagittal view with the region of interest (ROI) highlighted, whereas the right panel offers an enlarged view of the ROI, emphasizing critical anatomical structures

2: Select a point at the posterior border of the AC (Fig. 25).

**Figure 25.**
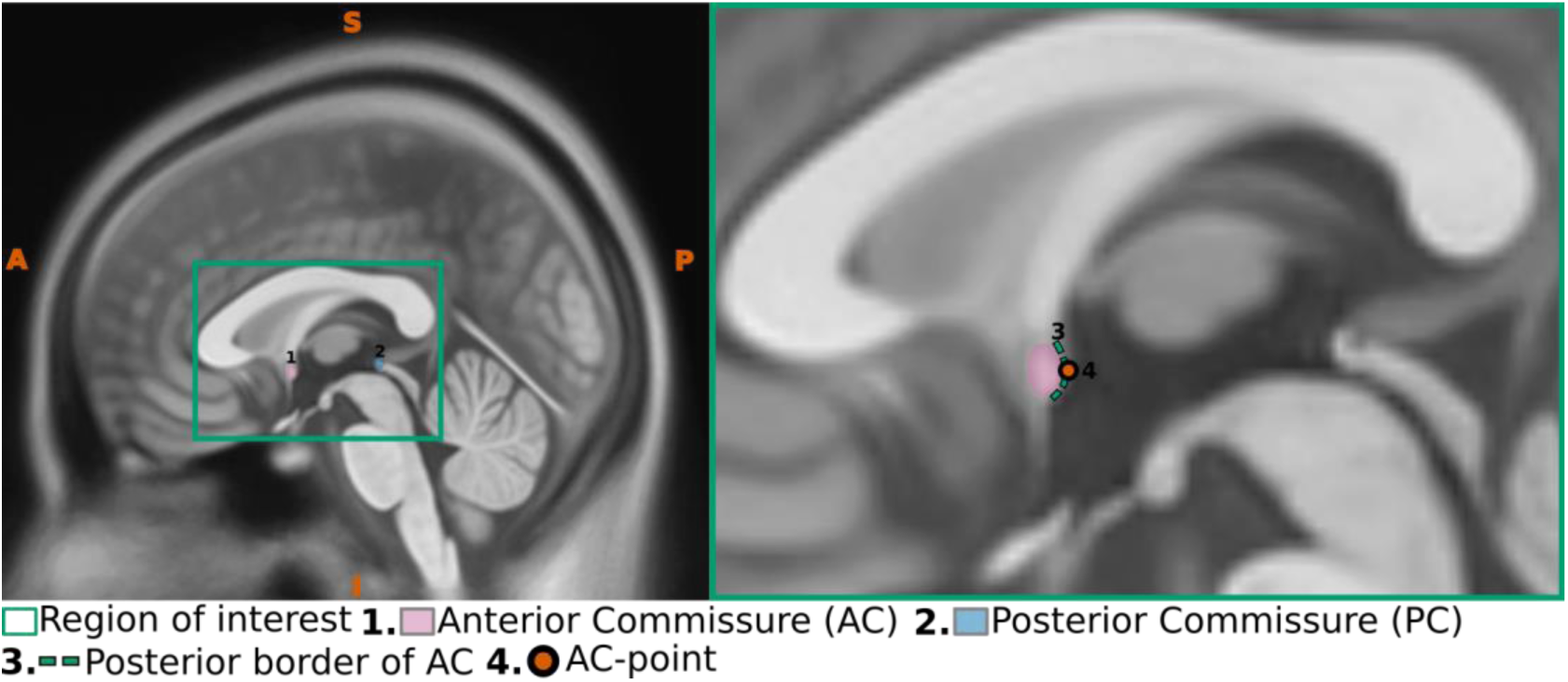
Sagittal view of the Anterior commissure. The left panel presents a midsagittal view with the region of interest (ROI) highlighted, whereas the right panel offers an enlarged view of the ROI, emphasizing critical anatomical structures

3. Use the corresponding axial plane to ensure the annotated point is accurately positioned along the midline (Fig. 26).

**Figure 26.**
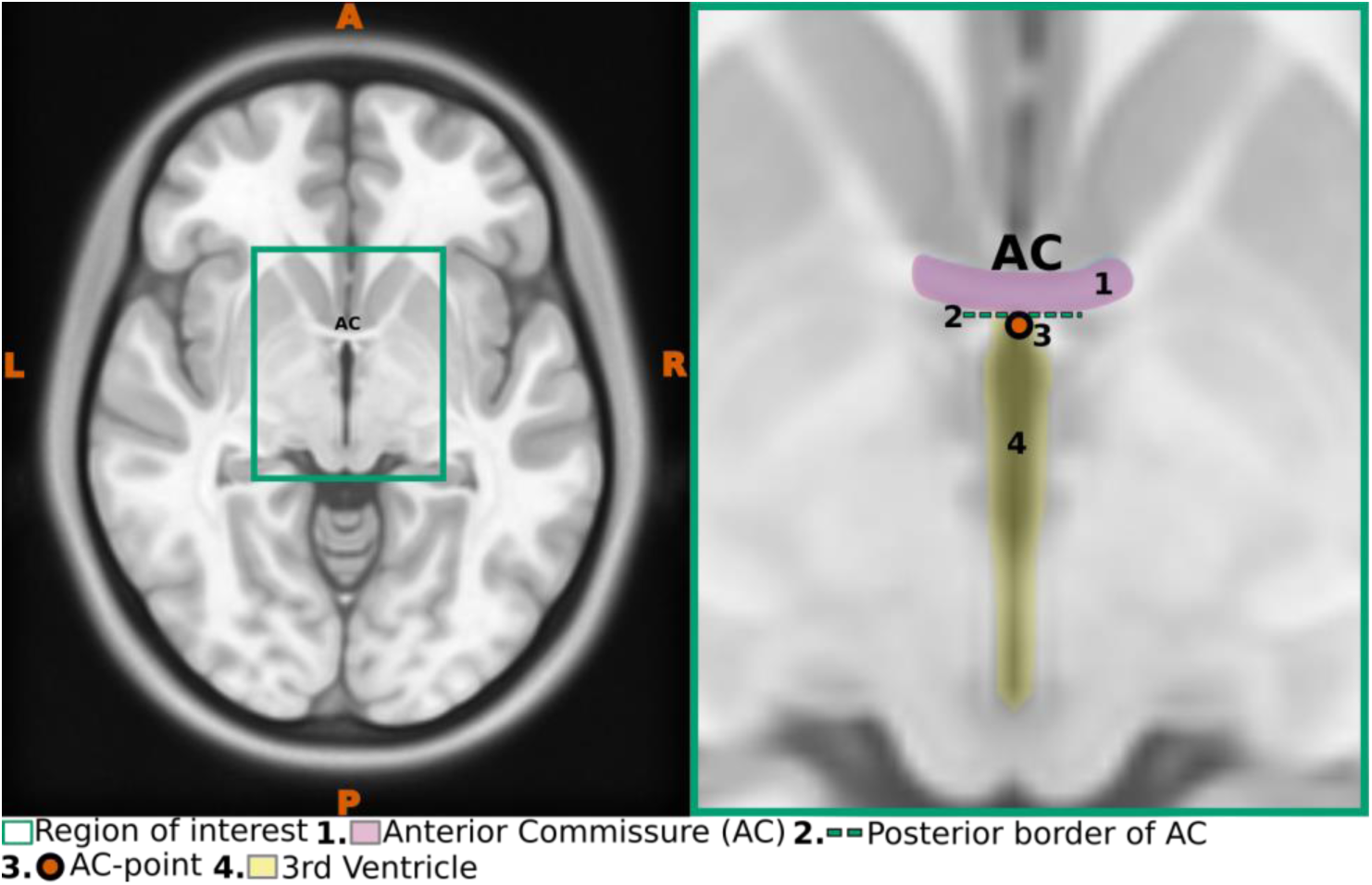
Axial view of the Anterior commissure. The left panel presents a axial view with the region of interest (ROI) highlighted, whereas the right panel offers an enlarged view of the ROI, emphasizing critical anatomical structures

#### 2.2.2. Posterior Commissure (PC)

1. Identify the Posterior commissure (PC; Fig 27).
2. Select a point at the anterior border of the PC (Fig. 28)
3. Use the corresponding axial plane to ensure the annotated point is accurately positioned along the midline (Fig. 29).

#### 2.2.3. AC-PC line

1. Ensure alignment along the AC-PC line, a straight line connecting the posterior border of the AC) to the anterior border of the PC). This line should pass through the midpoints of both the AC and PC in sagittal orientation. Accurate definition of this line is essential for precise AC-PC realignment (Fig. 30).

**Figure 27.**
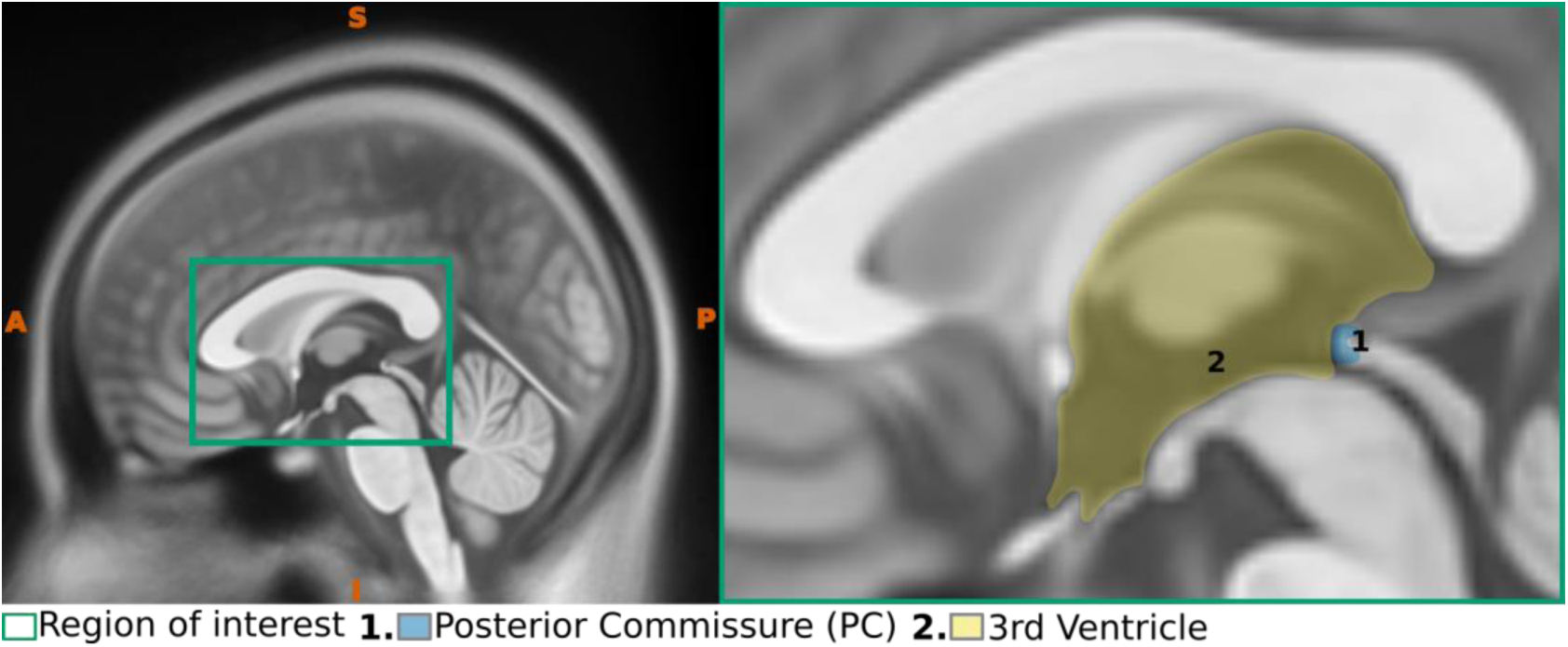
Sagittal view of the Posterior commissure. The left panel presents a midsagittal view with the region of interest (ROI) highlighted, whereas the right panel offers an enlarged view of the ROI, emphasizing critical anatomical structures

**Figure 28.**
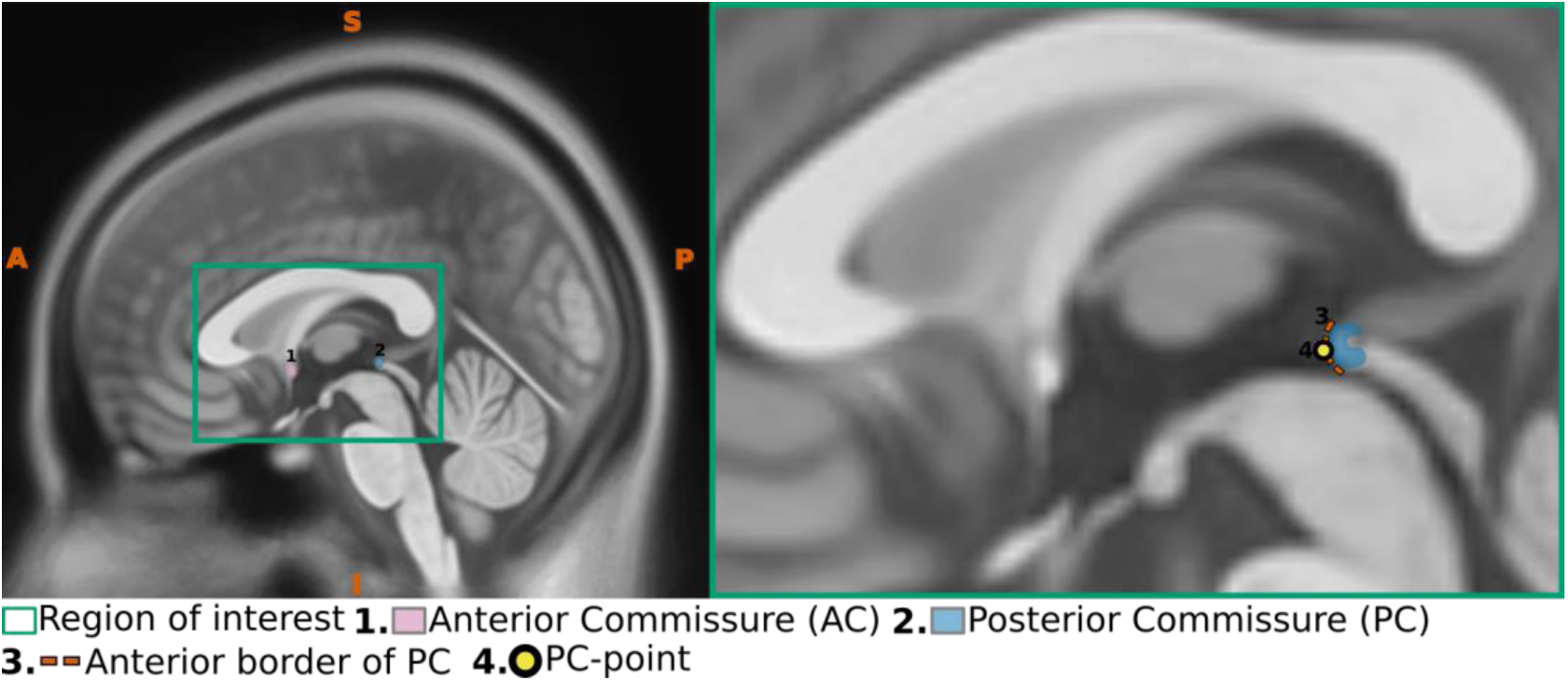
Sagittal view of the Posterior commissure. The left panel presents a midsagittal view with the region of interest (ROI) highlighted, whereas the right panel offers an enlarged view of the ROI, emphasizing critical anatomical structures

**Figure 29.**
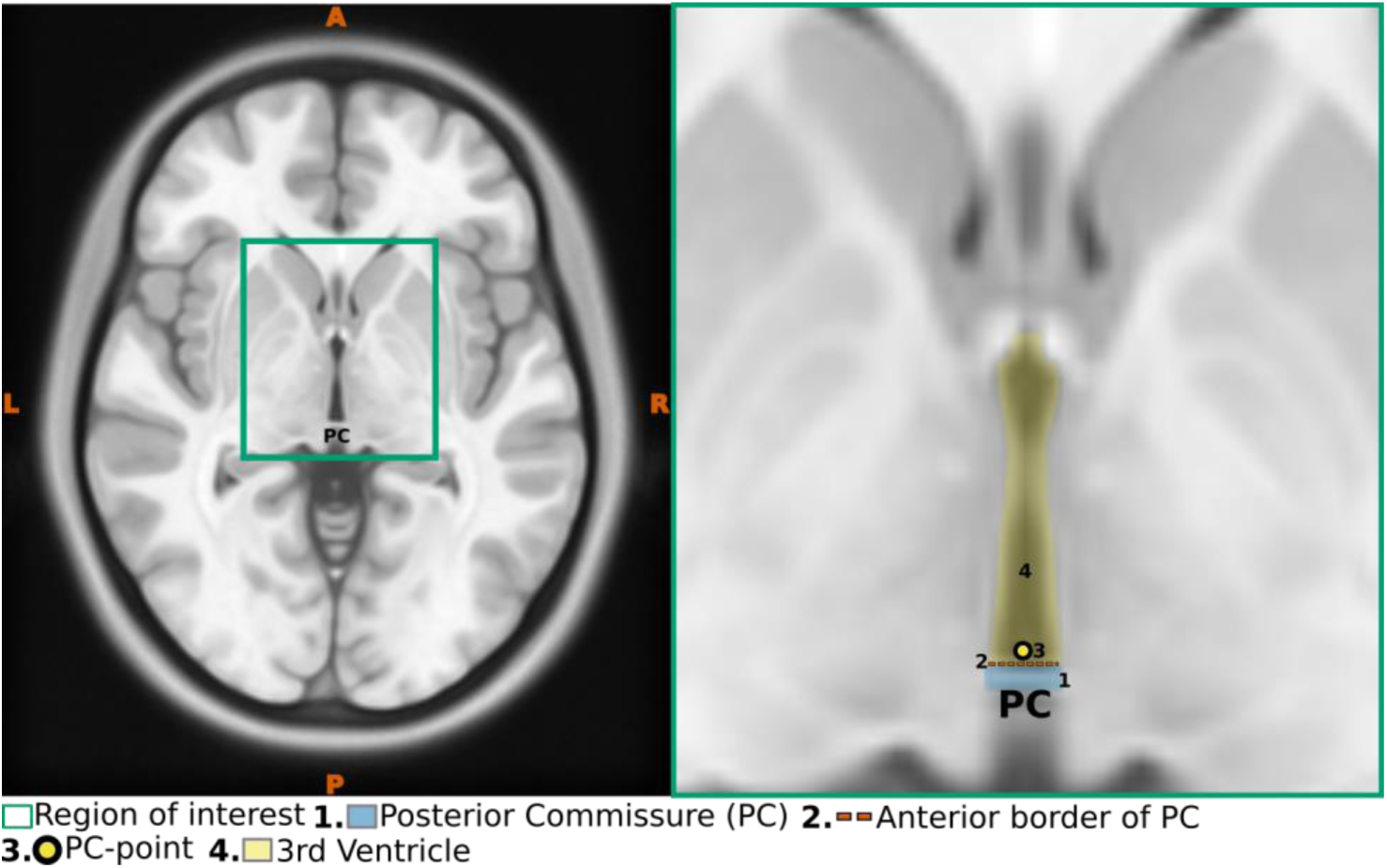
Axial view of the Posterior commissure. The left panel presents a axial view with the region of interest (ROI) highlighted, whereas the right panel offers an enlarged view of the ROI, emphasizing critical anatomical structures

**Figure 30.**
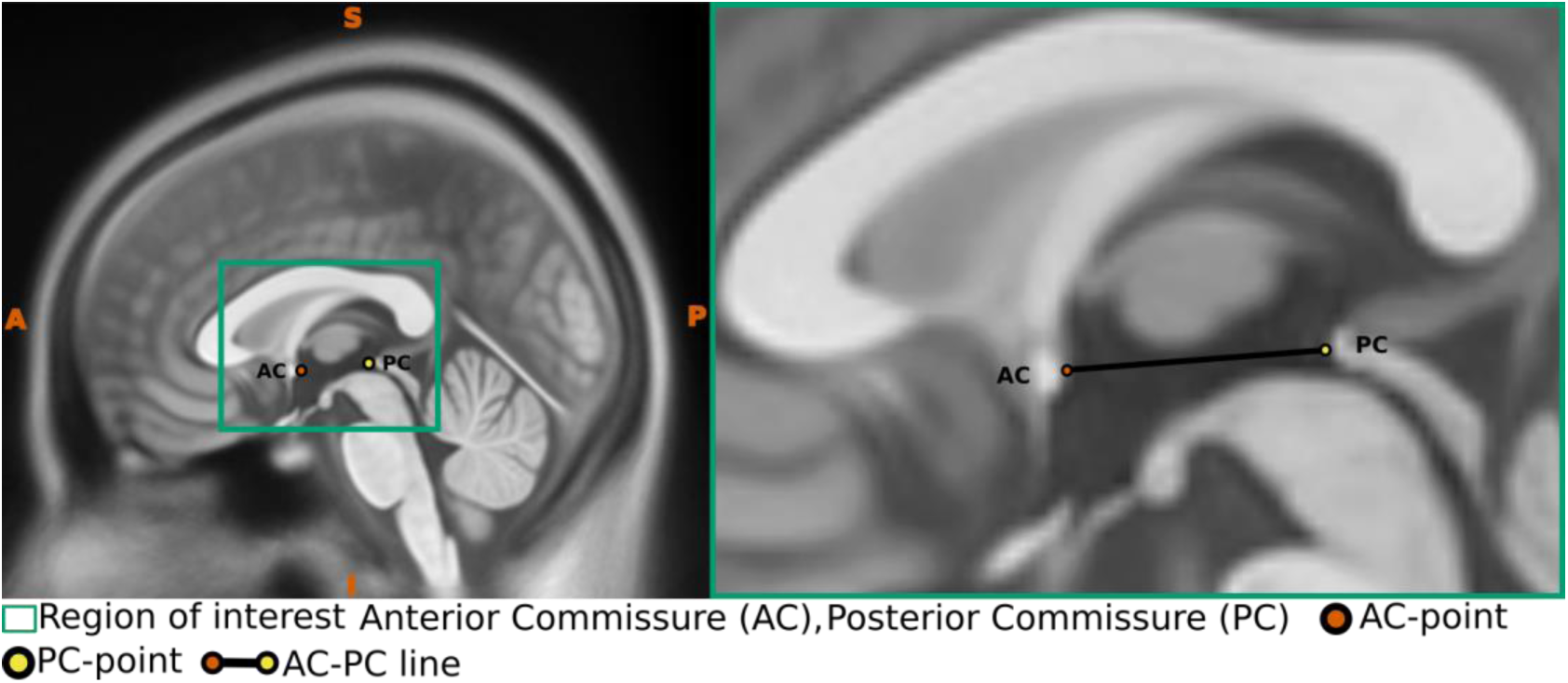
Sagittal view illustrating the AC-PC line. The left panel shows a midsagittal perspective with the region of interest (ROI) highlighted, while the right panel provides an enlarged view of the ROI, clearly emphasizing the AC-PC line.

#### 2.2.4. Superior Pontine notch

1. Identify the superior pontine notch (Fig. 31).

**Figure 31.**
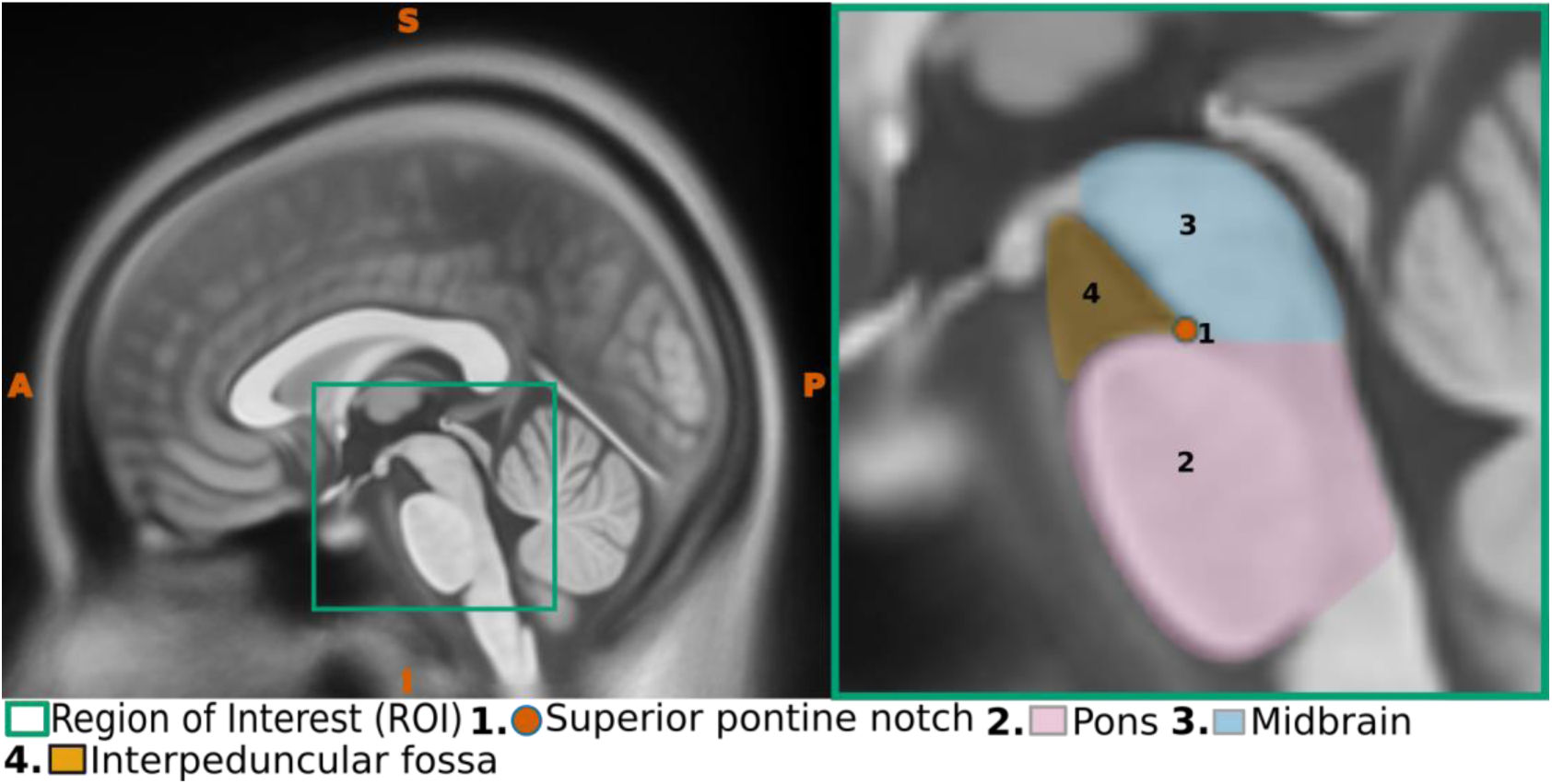
Sagittal view demonstrating the superior pontine notch. The left panel depicts a midsagittal perspective highlighting the region of interest (ROI), while the right panel provides a magnified view of the ROI, clearly emphasizing critical anatomical structures and specifically highlighting the superior pontine notch.

2. Place the first midline point in the superior pontine notch at the junction where the midbrain and pons meet (Fig. 31).

3. Use the corresponding axial slice to ensure the point is correctly positioned in the midline, where the two peduncles join to form the interpeduncular angle (Fig. 32).

**Figure 32.**
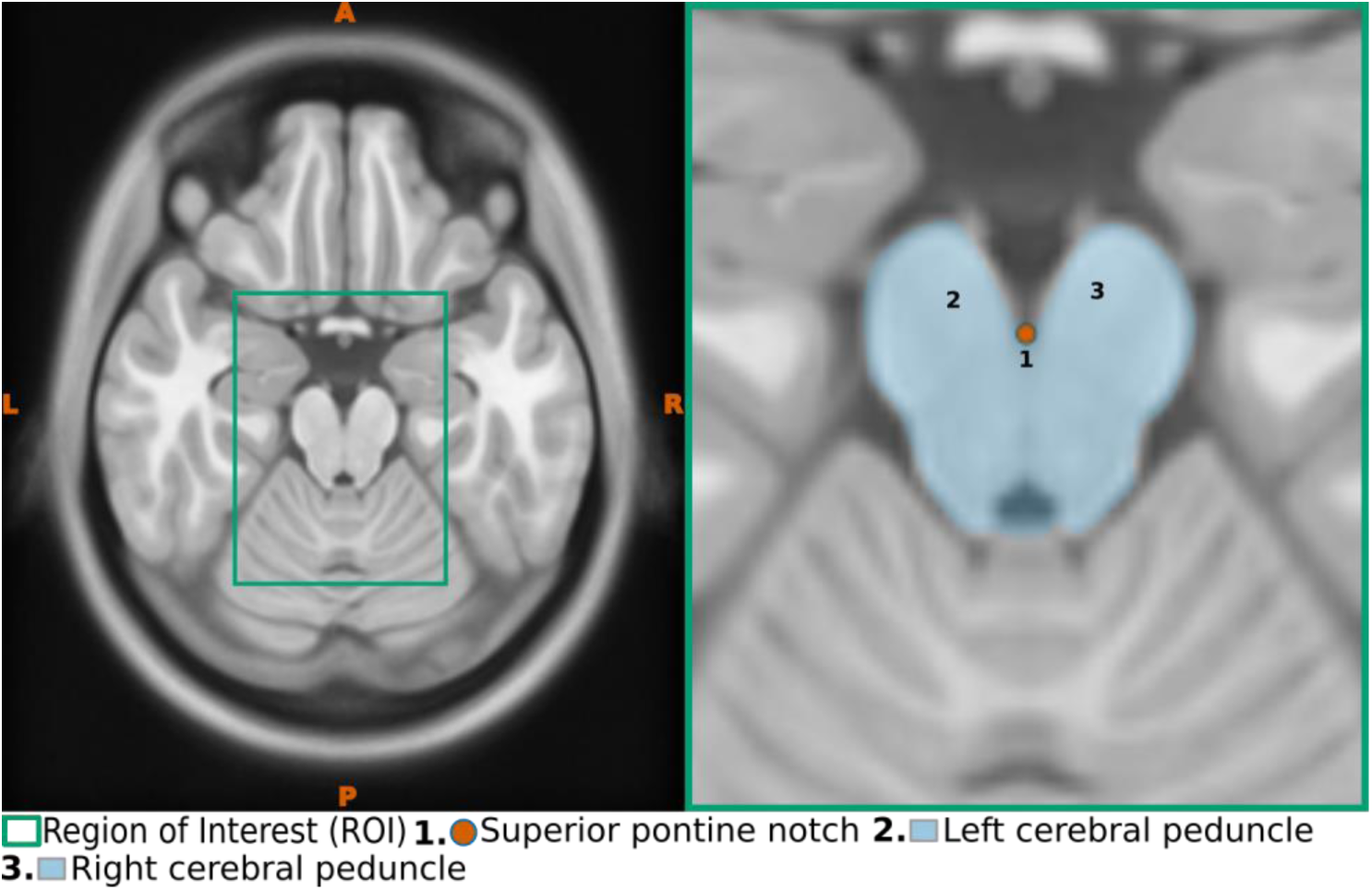
Axial view demonstrating the superior pontine notch. The left panel depicts a axial perspective highlighting the region of interest (ROI), while the right panel provides a magnified view of the ROI, clearly emphasizing critical anatomical structures and specifically highlighting the superior pontine notch

#### 2.2.5. Fastigium or Apex of the Roof of fourth ventricle

1. Identify the fastigium, also known as the apex of the roof of the fourth ventricle (Fig. 33).

**Figure 33.**
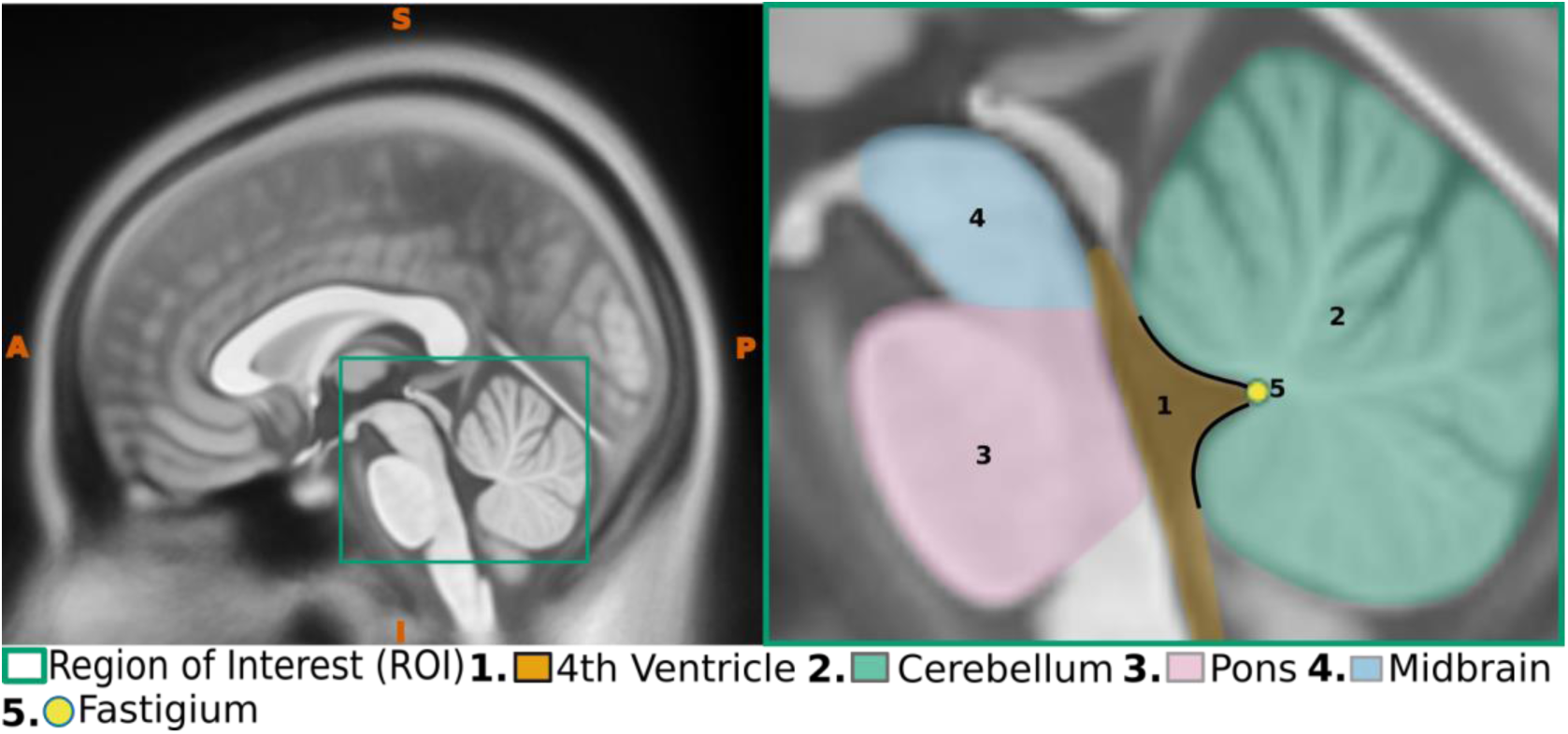
Sagittal view demonstrating the fastigium. The left panel depicts a midsagittal perspective highlighting the region of interest (ROI), while the right panel provides a magnified view of the ROI, clearly emphasizing critical anatomical structures and specifically highlighting the fastigium

2. Place the second midline point in the fastigium (Fig. 33).

3. Use the corresponding axial slice to ensure the point is accurately placed along the midline in the fastigium (Fig. 34).

**Figure 34.**
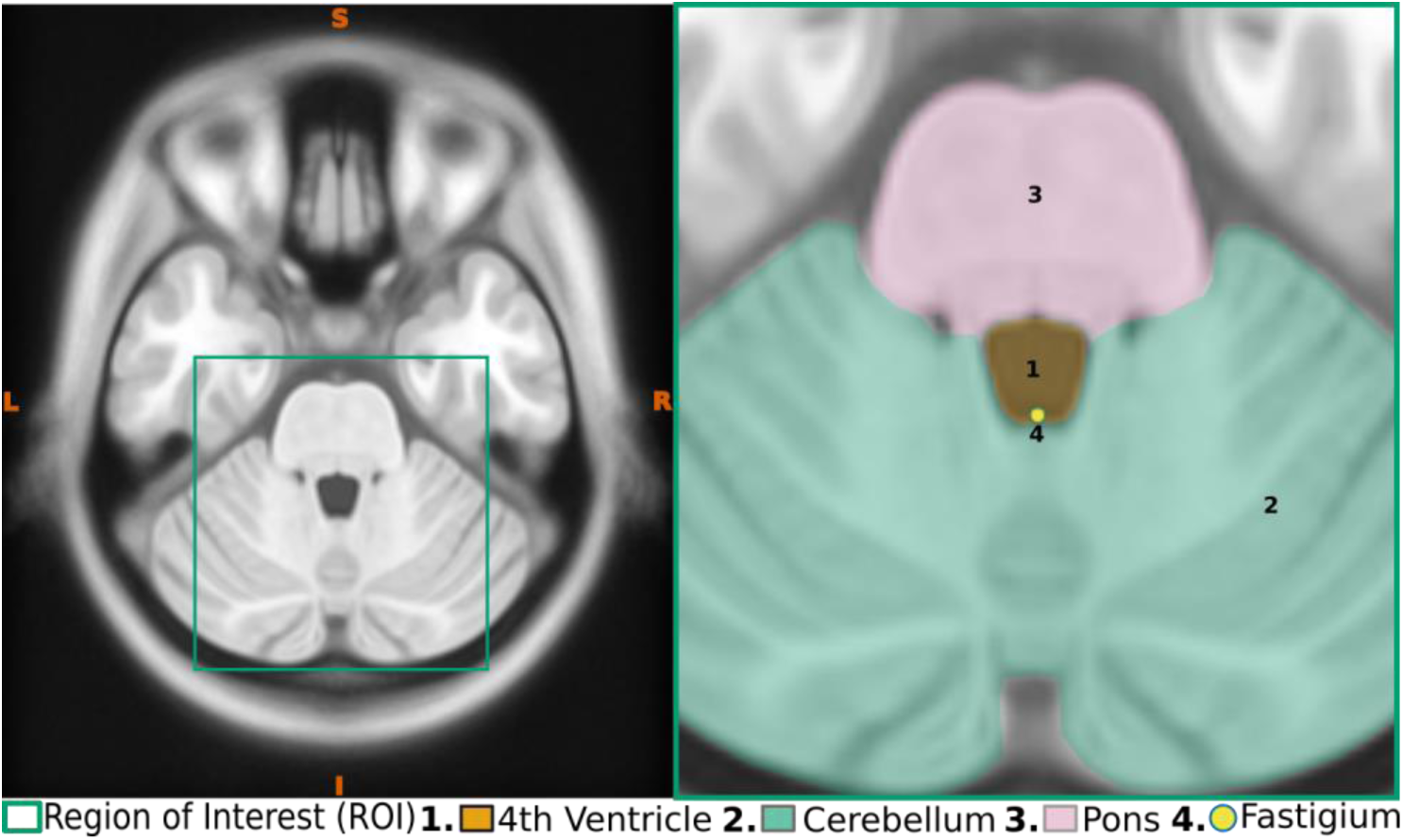
Axial view demonstrating the fastigium. The left panel depicts a axial perspective highlighting the region of interest (ROI), while the right panel provides a magnified view of the ROI, clearly emphasizing critical anatomical structures and specifically highlighting the fastigium

#### 2.2.6. Anterior and Posterior Points in the Falx Cerebri

1. Identify the corpus callosum (Fig. 35).

**Figure 35.**
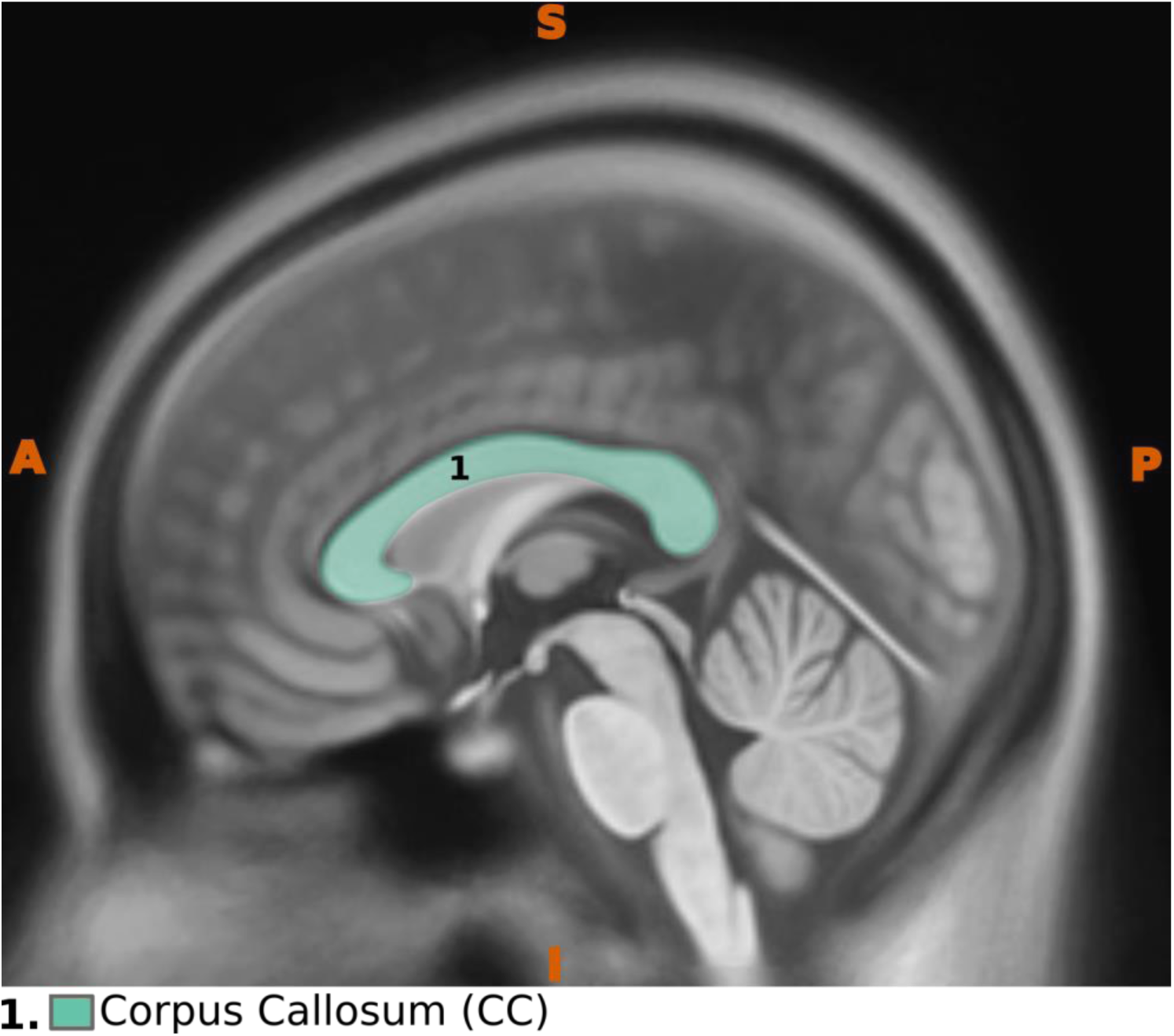
Sagittal view illustrating the corpus callosum

2. To mark its superior-most boundary, draw a line along the upper border of the corpus callosum, ensuring it is aligned with the highest point of this structure (Fig. 36). In conditions of minimal rotation, this should ideally correspond to the central body portion of the corpus callosum where it arches. However, the exact positioning may vary depending on the degree of rotation applied to the image. To maintain proper correspondence of the midpoint 3 and 4, it is essential to verify that this point corresponds to the superior-most section of the corpus callosum in a sagittal view.

**Figure 36.**
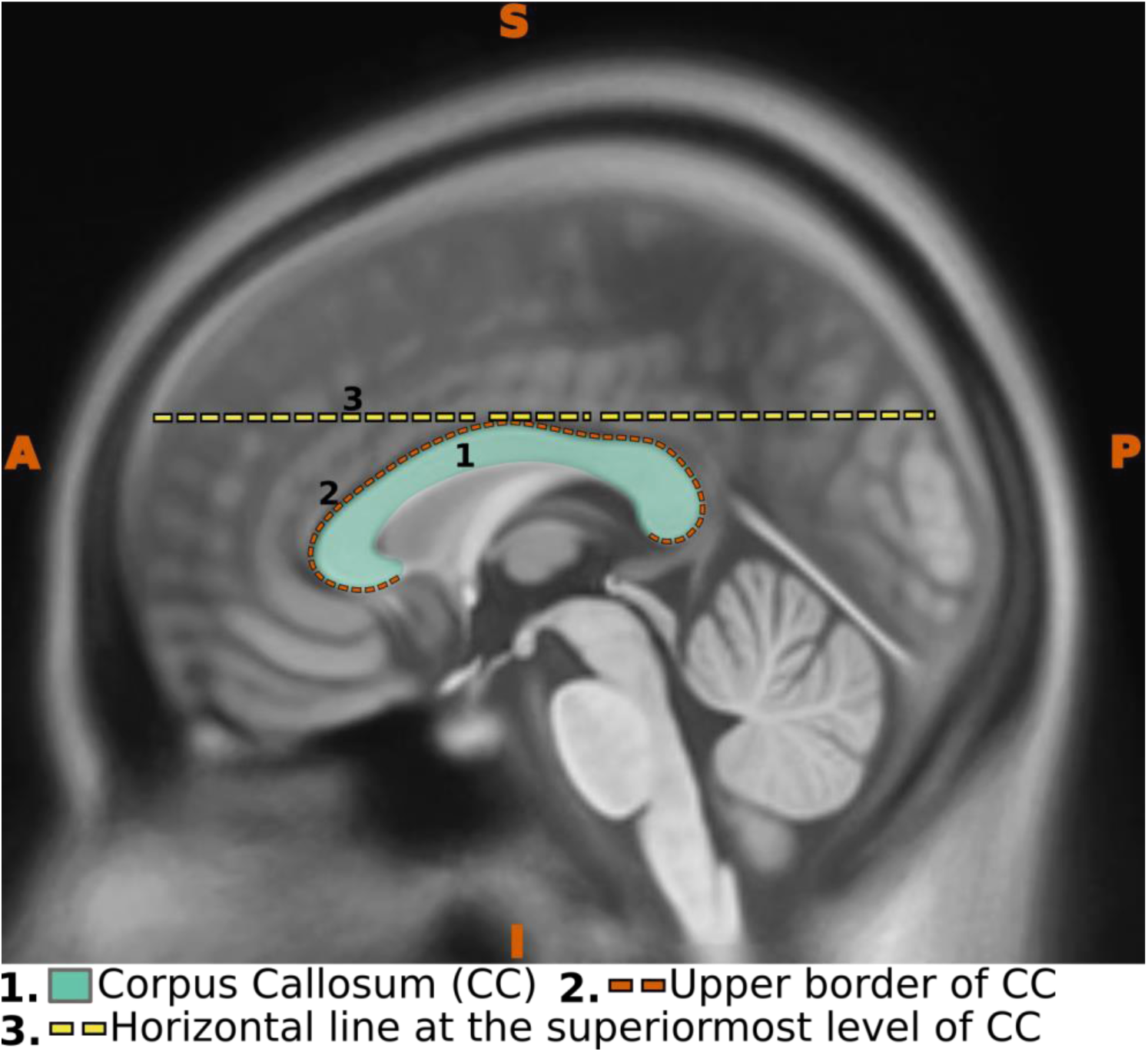
Sagittal view illustrating the corpus callosum with a horizontal line marking its superior-most boundary.

3. Use the corresponding axial plane to place two midline points (the 3rd and 4th points; Fig. 37): one at the anterior end of the falx cerebri (refer to Fig.13.3) and another at the posterior end. Ensure to place the points at the junction between the falx cerebri and the superior sagittal sinus. This junction can be identified as the apex of the triangular appearance of the superior sagittal sinus in a transverse section

**Figure 37.**
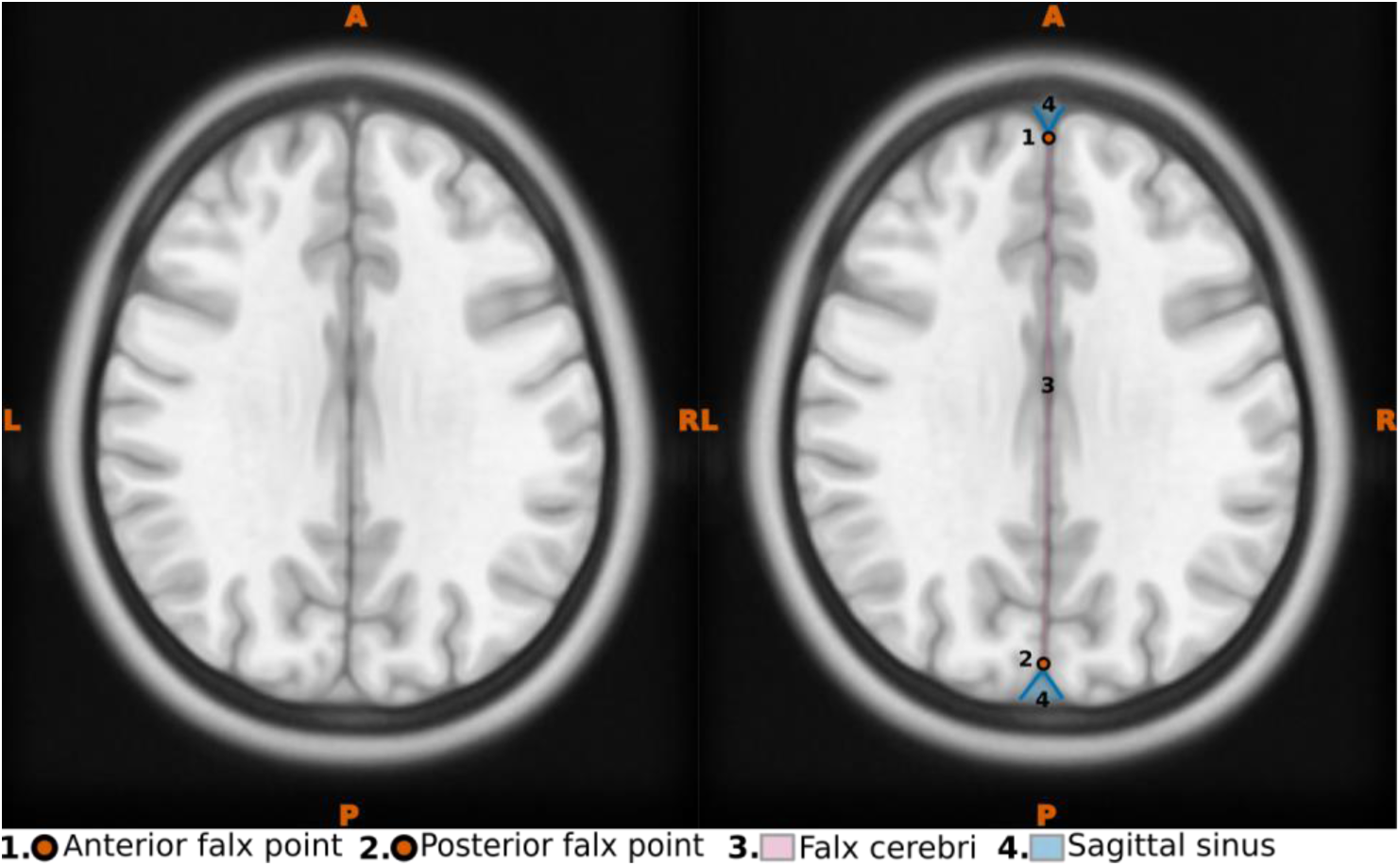
Axial view illustrating the falx cerebri at the midline. The left panel displays an axial perspective at the superior-most level of the corpus callosum, whereas the right panel highlights the falx cerebri and other key anatomical landmarks.

### 2.3 Verification of AC-PC alignment

After realigning the structural image to the AC-PC plane, it is essential to verify that the image is properly aligned. This involves checking for any rotations in the X, Y, and Z directions, commonly referred to as pitch, roll, and yaw.

#### 2.3.1. Verification of pitch

1. After successful realignment, the AC and the PC should be aligned on the same horizontal level. To verify this, draw two parallel lines along the upper and lower borders of the AC in the midsagittal plane. The space between these lines should encompass both the AC and PC, confirming their alignment (Fig. 38).

**Figure 38.**
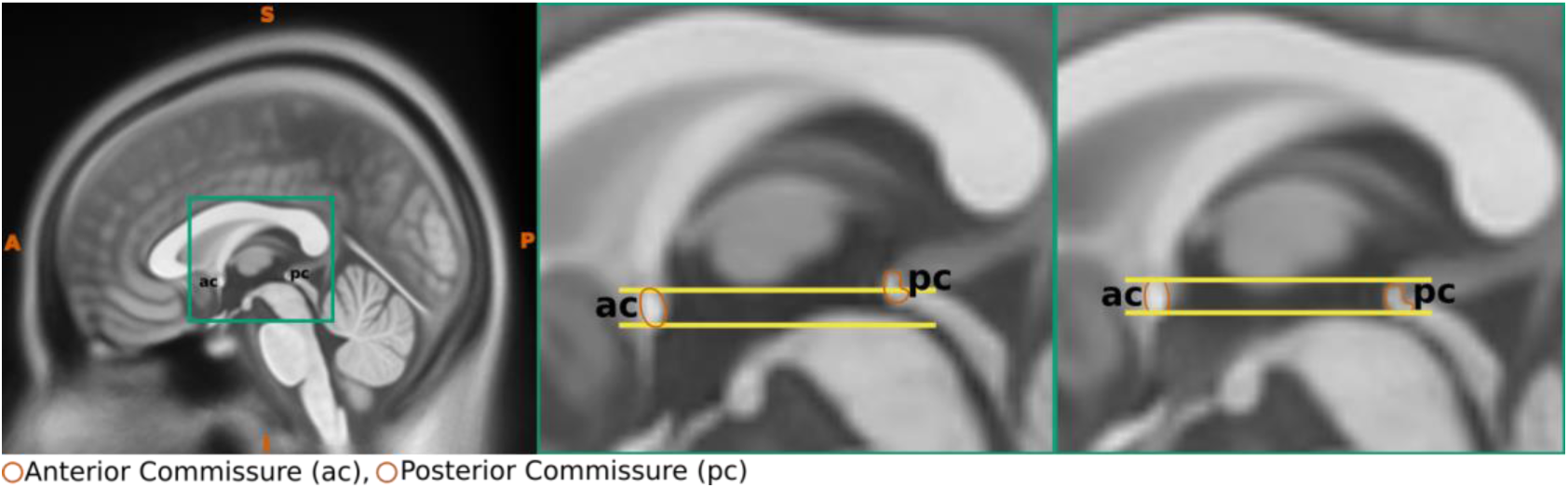
Verification of Pitch. From left to right, the first panel shows a midsagittal plane highlighting the region of interest (ROI). The second and third panels provide enlarged views of the ROI: the second panel illustrates the positions of the AC and PC before realignment, whereas the third panel displays their positions after realignment.

**Figure 39.**
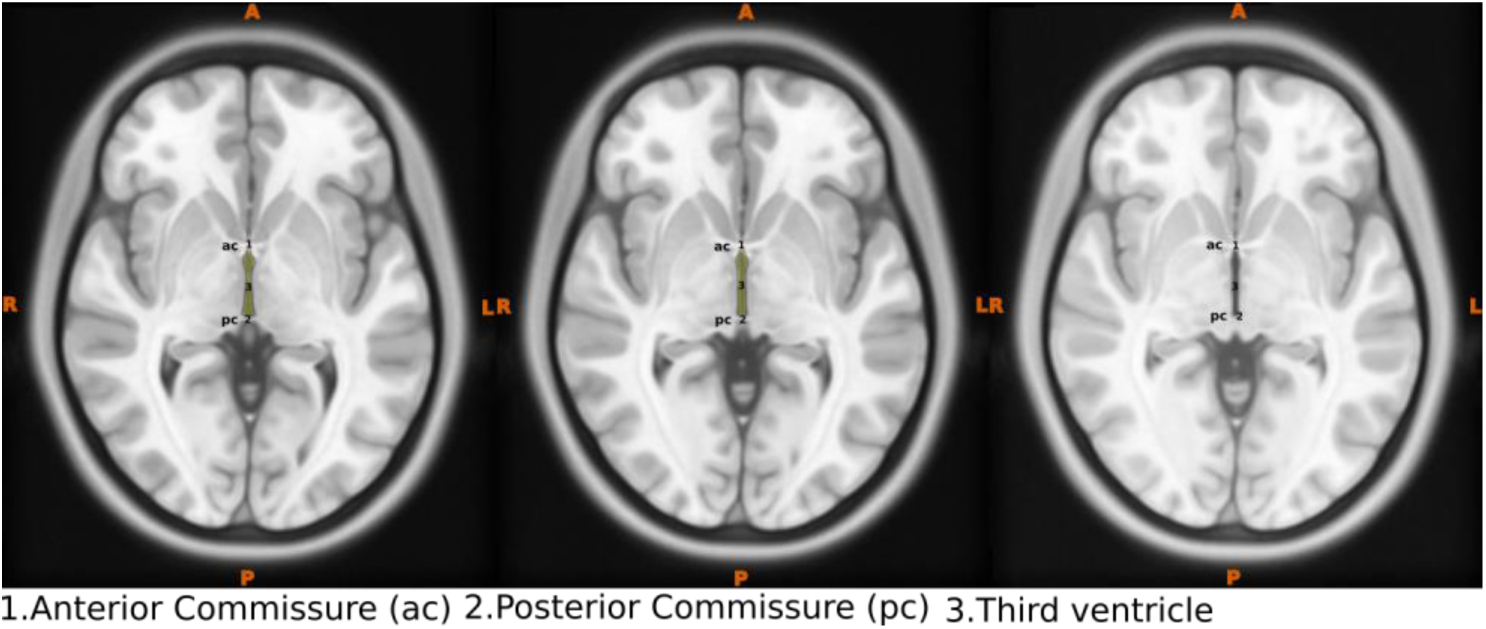
Verification of Pitch. Axial planes demonstrating criteria for verifying rotation around the Y-axis. Note the appearance of AC and PC in consecutive axial slices, as well as the characteristic keyhole shape of the third ventricle in the first two slices.

2. By examining the corresponding axial plane, the AC and PC should be visible within the same slice. For high-resolution images (with 1mm or smaller isotropic voxel size), both the AC and PC should appear across three consecutive slices. The presence of the AC and the PC in the same axial plane slice will create the characteristic keyhole appearance of the third ventricle

#### 2.3.2. Verification of roll

1: To verify the roll alignment, examine the position of the eyeballs in the selected axial slice; they should appear at the same level (Fig. 40). In cases where minor deviations are difficult to discern visually, coronal planes can provide an additional perspective to confirm precise AC-PC realignment (Fig. 41).

**Figure 40.**
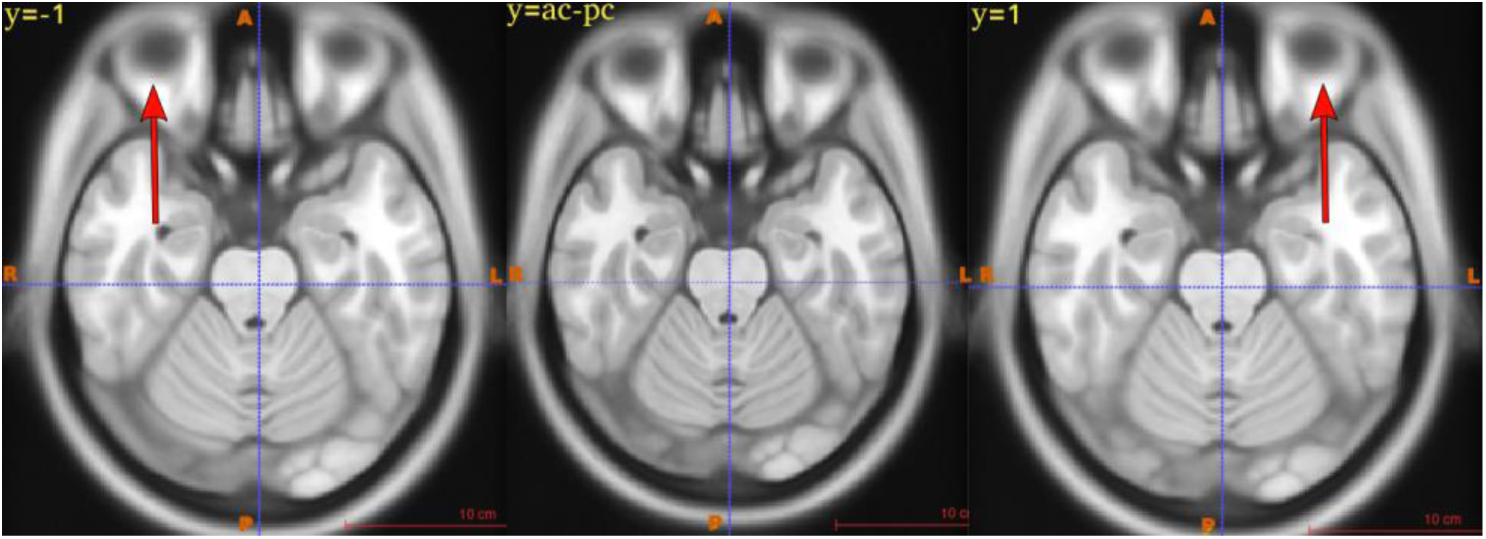
Verification of Roll. Axial planes illustrating the appearance of the eyeballs in three scenarios. From left to right: the first panel shows the right eyeball appearing larger than the left; the middle panel depicts both eyeballs equal in size; and the rightmost panel illustrates the left eyeball appearing larger. Note the subtlety of these differences.

**Figure 41.**
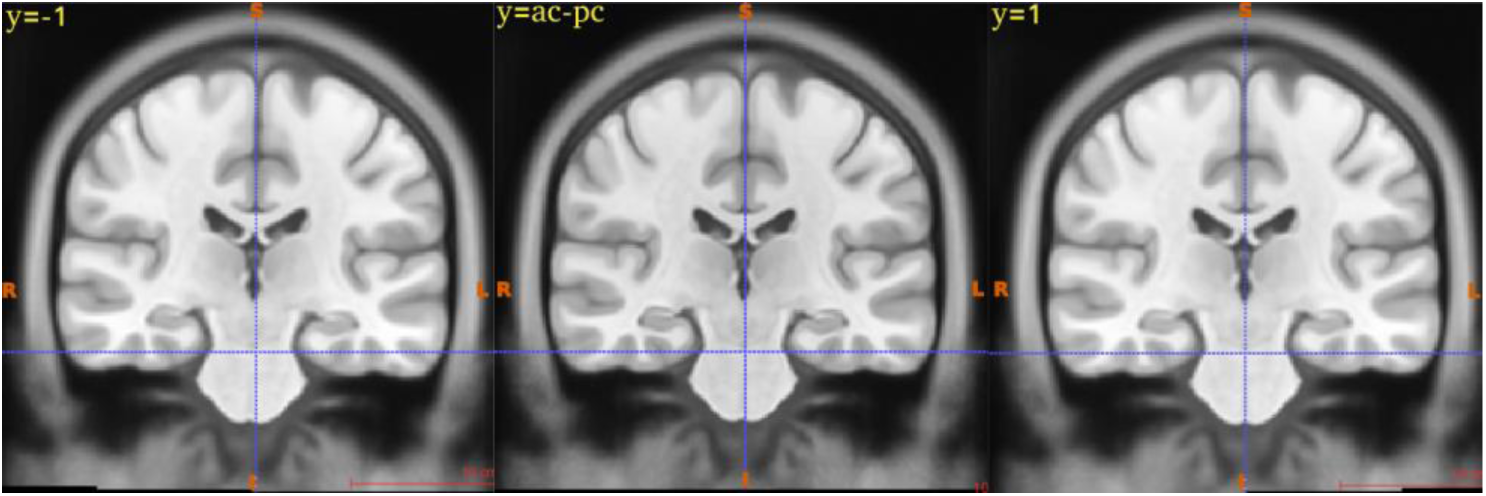
Verification of Roll. Coronal planes demonstrating midline deviations in three cases, marked by a vertical crosshair. From left to right: the first panel illustrates a deviation to the right, the middle panel shows proper midline alignment (no deviation), and the right panel reveals a deviation to the left. Note that these deviations are subtle, making them challenging to visually distinguish.

#### 2.3.3. Verification of yaw

1. The yaw could be verified by placing the vertical crosshair. The vertical line should align with the longitudinal fissure or midline of the brain in the axial plane (Fig. 42).

**Figure 42.**
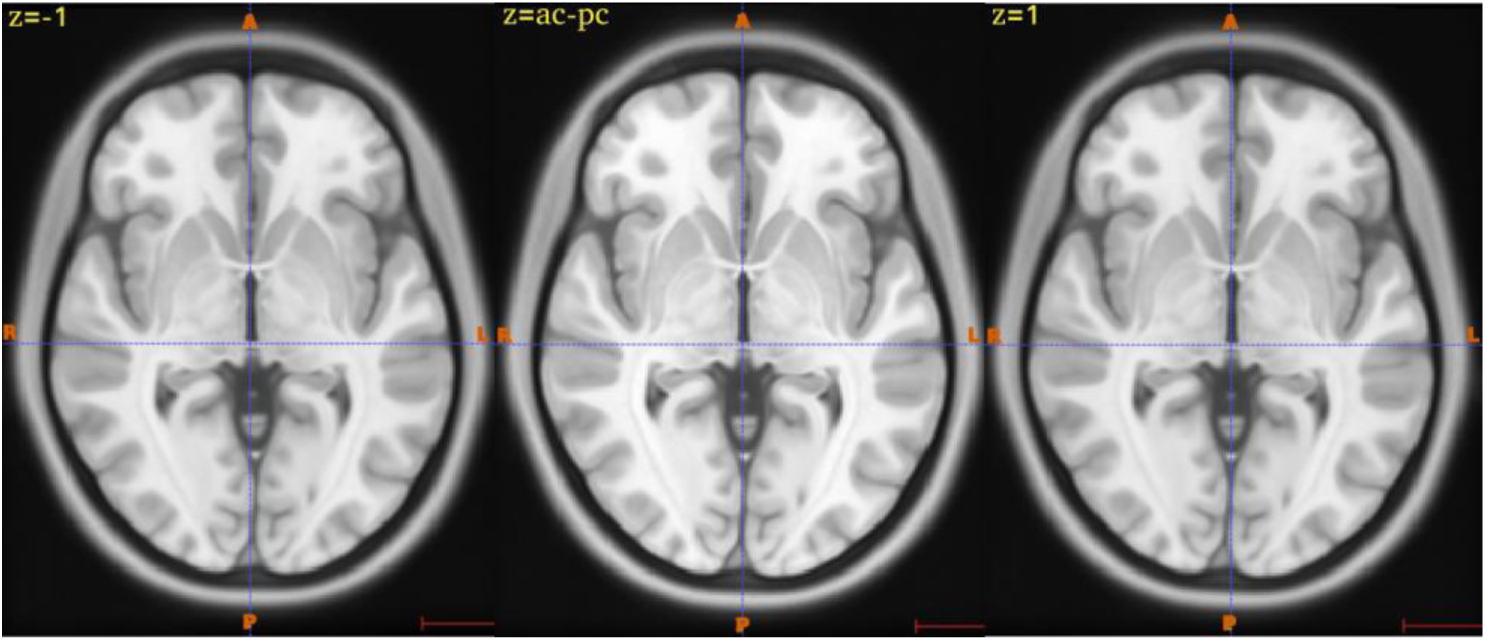
Verification of Yaw. Axial planes illustrating midline deviations in three cases, as indicated by the vertical crosshair. From left to right, the first panel illustrates a deviation to the left, the middle panel shows proper midline alignment (no deviation), and the right panel reveals a deviation to the right.

## 3 Justification

Achieving precise AC-PC alignment is contingent upon two critical factors: (1) an accurately defined AC-PC line and (2) a clearly established midplane. These elements form the foundation of our approach in developing this guideline, with each design choice grounded in careful consideration of identifiable anatomical landmarks that require no prior anatomical expertise. Moreover, these choices have been refined through experimental procedures and feedback from the six manual observers who participated in the experiment

### 3.1 Design Choices

#### 3.1.1. Definition of the AC-PC line

We adopted the clinically used definition of the AC-PC line (Fig.1d), which is defined from the posterior border of the AC to the anterior border of the PC (Acar et al., 2007; Caire et al., 2009; Horn et al., 2017; Patel et al., 2008; Saint-Cyr et al., 2002; Schuurman et al., 1999). The Schaltenbrand atlas is the cornerstone reference for modern stereotactic neurosurgery (Gildenberg et al., 2009; Pallavaram et al., 2009). However, there is a notable variation in the definition of the AC-PC line between clinical practice and the original atlas version. (Horn et al., 2017) (Fig.1C-D). This discrepancy arises from the necessity to adapt a post-mortem atlas for in vivo imaging, a challenge that was particularly prominent before the advent of MRI technology (e.g. for cerebral ventriculography images). Despite advancements in imaging, this adaptation persists in current surgical practices (Lau, 2022; Pallavaram et al., 2009).

The other variations like the tangential inter-commissural line, as used in the Talairach atlas (Fig.1B), are influenced by factors such as the size and shape of the AC and the PC. These factors can significantly affect the angle of the AC-PC line (Choi et al., 2013; Nowinski, 2001). Given that the AC and PC can vary in shape, ranging from round to elliptical, defining the AC-PC line based on the superior and inferior edges becomes unreliable and prone to errors (Park et al., 2010). This issue is further compounded by poor image resolution, making accurate definition even more challenging (Choi et al., 2013).

Besides its status as a surgical convention, defining the AC-PC line from the posterior border of AC to anterior border of PC offers several advantages. First, identifying the borders of the AC and PC is more straightforward, as this can be accomplished by tracing the boundaries of the third ventricle, which is often easier than locating their centers or superior and inferior edges. Second, defining the line from the borders of the AC and PC remains consistent across different imaging modalities and various populations (e.g. healthy vs. atrophied). Moreover, the borders are not influenced by the shape and size of the AC and PC (Choi et al., 2013). Third, reliable identification of the borders of the AC and PC does not require high-resolution images or explicit sagittal slices (Otake et al., 2018). Unlike the centers and superior and inferior edges of the AC and PC, the borders can be identified in axial slices and low-resolution images (Choi et al., 2013; Sandor et al., 1994).

#### 3.1.2. Selection of the midline-points

To robustly define the midplane, we require at least three non-collinear points. In the proposed protocol, we have introduced four midline points to facilitate midplane definition (Fig. 43). The four proposed midline points are:

1. the superior pontine notch
2. the fastigium of the fourth ventricle
3. the anterior point in the falx cerebri
4. the posterior point in the falx cerebri

**Figure 43.**
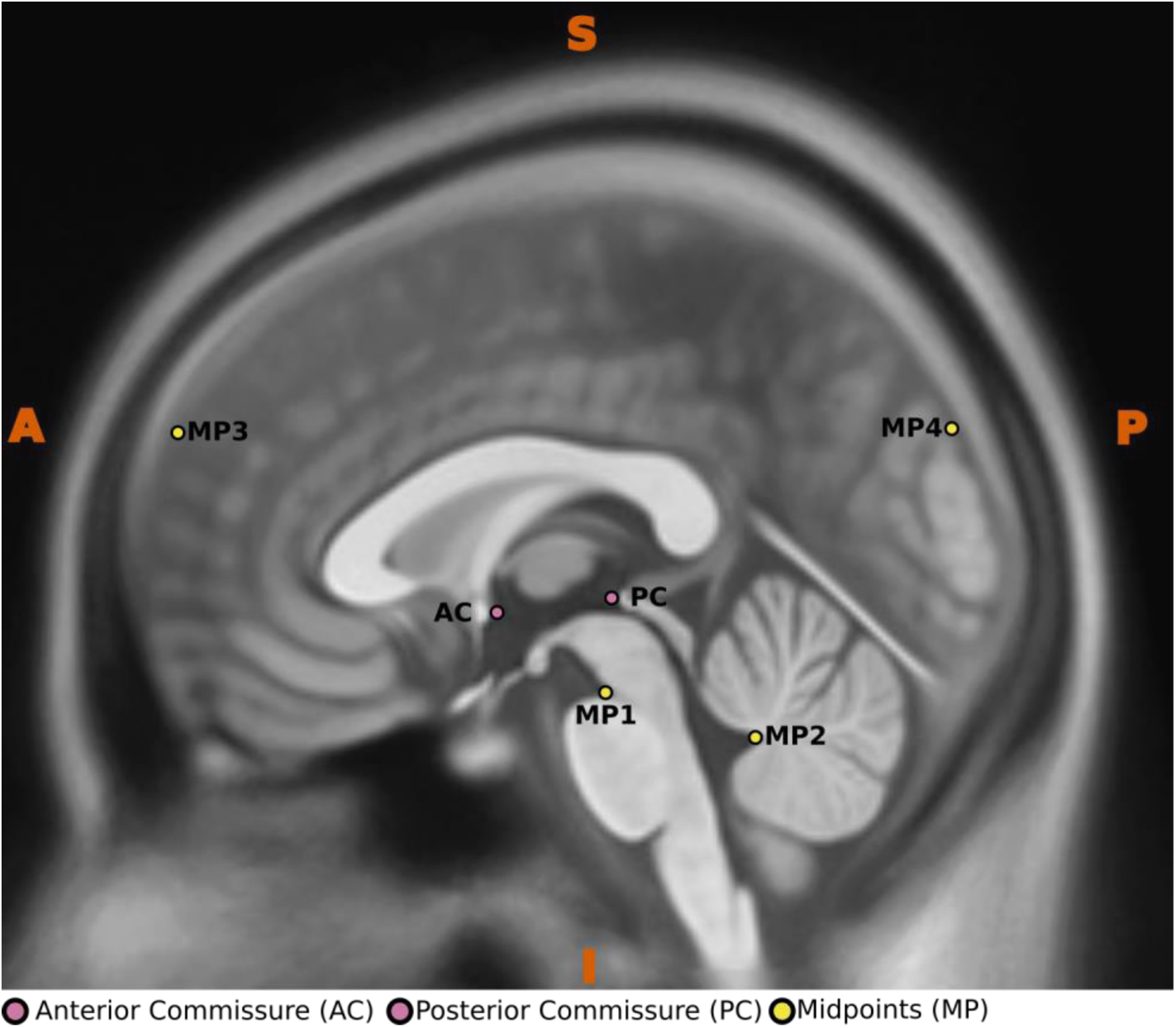
Sagittal view illustrating all landmark points, including the Anterior Commissure (AC), Posterior Commissure (PC), and the four midline reference points (MP).

The selection of these four specific landmarks is based on their extensive use across both clinical and research contexts including ac-pc realignment, which establishes them as highly validated reference structures.(Ardekani & Bachman, 2009; Chen et al., 2010; Horn et al., 2017; Liu & Dawant, 2015; Niemann et al., 1999; Oba et al., 2005; Otake et al., 2018; Zrinzo et al., 2008). Their consistency, reliability, and ease of identification across various imaging modalities, whether assessed manually or through automated computer vision techniques, enhance their versatility, making them invaluable landmarks for a wide range of applications (Ardekani & Alzheimer’s Disease Neuroimaging, 2022; Lau et al., 2019; Roelants et al., 2016; Shah et al., 2013).

Moreover, these landmarks offer a thorough and extensive representation of the brain’s anatomy, with two positioned at the supratentorial level specifically at the falx cerebri and two located at the infratentorial level, namely the superior pontine notch and the fastigium of the fourth ventricle. This distribution helps ensure a balanced representation of the brain’s anatomy, minimizing the impact of any anatomical or imaging anomalies that may arise (Liu et al., 2001; Shah et al., 2013; Yap et al., 2023).

Additionally, we propose to define two distant points lying on the same line to minimize the error in plane fitting and to span a longer plane (Sandor et al., 1994), which is particularly useful when employing algorithms like Principal Component Analysis (PCA) to fit a plane. For instance, the AC-PC transformation model in 3D Slicer utilizes this PCA-based approach to accurately fit a plane.

#### 3.2.3 Selection of verification criteria

As the final step in our protocol, we propose three visualization criteria to verify successful AC-PC realignment. Clinically, AC-PC realignment requires rotating the image around the three cardinal planes (x, y, z) to align the AC-PC at the same level within the horizontal plane, as outlined in the Schaltenbrand atlas (Caire et al., 2011; Saint-Cyr et al., 2002). Consequently, the verification criteria for successful AC-PC realignment are specifically designed to ensure the accuracy of rotations around the X, Y, and Z axes. Among these criteria, verifying the pitch or X-axis rotation is the most critical, as it confirms that the image is properly aligned with the clinically utilized AC-PC plane (Hamilton et al., 2017; Lancaster et al., 1995; Ramirez et al., 2014). Since both the AC and the PC can be clearly visualized in the mid-sagittal and axial planes, we utilized both planes to verify the level of AC and PC to ensure precise X-axis rotation. It is important to note that the criteria to verify pith or X-axis rotation is specifically applicable to verify the Schaltenbrand definition of the AC-PC plane (Horn et al., 2017), encompassing both the clinical and original versions. To verify the roll or Y-axis rotation, we utilized both the axial and coronal planes. In the axial plane, we focused on aligning the level of the eyeballs, while in the coronal plane, we emphasized ensuring midline alignment (Hamilton et al., 2017; Lancaster et al., 1995; Ramirez et al., 2014). Finally, to verify the yaw or Z-axis rotation, we employed the axial plane to assess midline alignment (Hamilton et al., 2017; Lancaster et al., 1995; Ramirez et al., 2014). The criteria addressing roll (Y-axis rotation) and yaw (Z-axis rotation) are essential for maintaining proper alignment and symmetrical positioning of the brain. Consequently, these two criteria can be used to verify all definitions of AC-PC plane alignment, including those established by Talairach.

## 4 Validation procedure of precursor guideline

### 4.1 Data preparation and experimental setup

The proposed guideline integrates essential aspects from a precursor guideline that was refined through a combination of experimental results and observer feedback. A detailed version of the precursor guideline, along with the experimental procedures and part of the results, is provided in the supplementary materials. The evaluation procedure can be summarized as follows: three images representing healthy, aged, and Parkinson’s disease (PD) groups were preprocessed and aligned to the clinically utilized AC-PC plane. These AC-PC realigned images were subsequently rotated at three different rotation values. The rotated images, referred to as *R*_min_, *R*_avg_, and *R*_max_, were used in a two-stage evaluation procedure. In stage 1, six manual observers, two experts, and four non-experts were asked to realign these rotated images to the clinically established AC-PC plane following the precursor guideline. Additionally, the experts performed the realignment independently without using the precursor guideline (prior to the realignment with the guideline). In stage 2, automated AC-PC realignment was conducted using various methods, including the model-based “acpcdetect” algorithm which automatically detects the AC and PC landmarks and realigns the structural T1w MR image to the Schaltenbrand version of the AC-PC plane (Ardekani & Alzheimer’s Disease Neuroimaging, 2022; Ardekani & Bachman, 2009; Ardekani et al., 1997). The template-based alignment involves linearly registering the structural image to template space using two templates: the commonly used MNI152 template and a version realigned to the clinically utilized AC-PC plane (Glasser et al., 2013). Additionally, the points warping method nonlinearly transforms landmark points from the template image to the individual image, defining the AC-PC line and midplane in the subject-specific space for subsequent AC-PC realignment. We tested two methods for defining points in the template image: the first followed the precursor guideline, while the second was based on the approach by Horn et al. (Horn et al., 2017; Oxenford et al., 2022), which employs different midline landmarks than those in our precursor guideline. The experimental results were evaluated against a baseline established by the first author (AS). Key metrics included the spatial differences in the locations of the borders of AC and PC, and four proposed midline landmarks, as well as variations in rotation and root mean squared deviations (RMSD) compared to the baseline. It is noteworthy that not all automated methods provided spatial coordinates for the AC, PC, or midpoint; in such cases, only the rotational errors and RMSD were compared to assess alignment accuracy.

### 4.2 Results

Since rotational errors and Root-mean-square deviation (RMSD) are the two most relevant metrics available across all cases for both automated and manual methods, our discussion here focuses primarily on these key metrics. Comprehensive results detailing the spatial differences in the locations of the AC and PC borders, along with the four proposed midline landmarks are provided in the supplementary material.

#### 4.2.1. Performance of Experts

##### 4.2.1.1 Root-mean-square deviation (RMSD)

RMSD analysis revealed that experts performed better in the *R*_avg_ and *R*_max_ rotation conditions compared to *R*_min_, where the largest deviations were observed. Nevertheless, the overall mean deviation across all three rotation scenarios remained below 1.5 mm, regardless of guideline use. Although adherence to the guideline resulted in slightly improved AC–PC alignment, the difference was marginal and inconsequential.

##### 4.2.1.2 Rotational Errors

The detailed analysis of these rotational errors (Table 3) shows that, in the absence of the guideline, prominent rotation errors were made around the X-axis, with maximum deviations of 1.52°, 1.12°, and 1.79°, for *R*_min_, *R*_avg_, and *R*_max_ respectively. While most rotation errors around the Y- and Z-axes remained below 1°, a few exceptions were noted specifically: deviations of 1.44° for *R*_min_ and 1.39° for *R*_avg._ The only notable Z-axis deviation without the guideline was 0.94°, observed for *R*_min_.

**Table 3.** Rotation angles (in degrees) for all three axes, comparing the baseline with expert assessments under both guideline and non-guideline conditions. Additionally, it shows the absolute difference values (in brackets) between the experts’ measurements and the baseline.

| | $R_{min}$ | | | $R_{avg}$ | | | $R_{max}$ | | |
| --- | --- | --- | --- | --- | --- | --- | --- | --- | --- |
|  | Without guideline |  |  |  |  |  |  |  |  |
| | $x$ | $y$ | $z$ | $x$ | $y$ | $z$ | $x$ | $y$ | $z$ |
| Baseline | -11.65 | -0.35 | 0.09 | 8.54 | 3.48 | 2.26 | 27.27 | -0.74 | -4.58 |
| Expert 1 | -10.15(1.50) | -0.59(0.24) | 0.07(0.01) | 9.66(1.12) | 3.82(0.34) | 2.45(0.18) | 29.06(1.79) | -1.02(0.28) | -4.19(0.39) |
| Expert 2 | -10.13(1.52) | 1.09(1.44) | -0.85(0.94) | 8.80(0.26) | 4.87(1.39) | 2.48(0.21) | 27.22(0.05) | -0.25(0.49) | -4.16(0.42) |
|  | With guideline |  |  |  |  |  |  |  |  |
| Expert 1 | -9.52(2.13) | -0.39(0.04) | 0.33(0.24) | 9.88(1.35) | 3.43(0.05) | 2.44(0.18) | 28.01(0.74) | -0.21(0.52) | -4.67(0.10) |
| Expert 2 | -10.84(0.81) | 0.00(0.36) | 0.11(0.02) | 7.23(1.31) | 3.67(0.19) | 2.48(0.21) | 25.79(1.48) | -0.04(0.70) | -4.68(0.10) |

Similarly, when the guideline was applied, notable rotation errors were predominantly observed around the X-axis across all three rotation scenarios. The maximum recorded deviations were as follows: *R*_min_ = 2.13°, *R*_avg_ = 1.35°, and *R*_max_ = 1.48. In contrast to the no-guideline condition, rotation errors around the Y- and Z-axes consistently remained below 1° with the guideline in place. In summary, although the observed differences were relatively minor, noticeable deviations persisted in both conditions.

#### 4.2.2 Performance of Non-Experts

##### 4.2.2.1 Root-mean-square deviation (RMSD)

RMSD analysis showed that non-experts achieved lower errors in the *R*_min_ and *R*_avg_ rotation conditions than in *R*_max_, where the two largest deviations 3.38 mm and 2.82 mm were recorded (Table 4). These outliers elevated the overall mean RMSD across the three rotation scenarios to 2.13 mm and 1.53 mm, respectively. Even so, the results indicate that, with clear guidelines, non-experts can perform as well as experts.

**Table 4.**
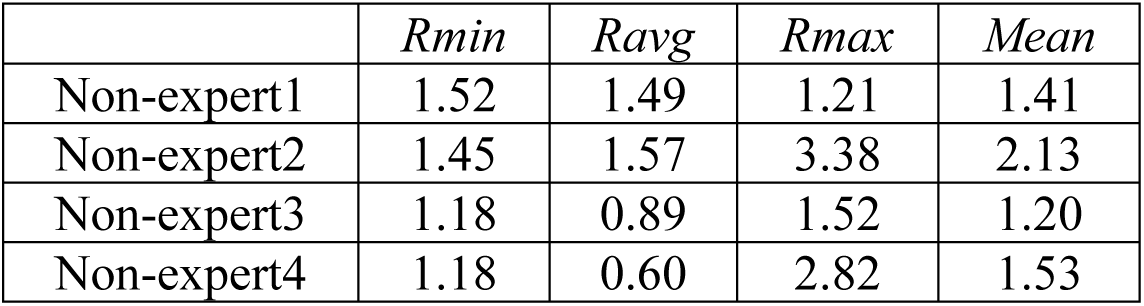
This table presents RMSD values (in mm), computed from rotation matrices, quantifying the deviation of non-expert manual transformations from the baseline.

| | $R_{\min}$ | $R_{\text{avg}}$ | $R_{\max}$ | Mean |
| --- | --- | --- | --- | --- |
| Non-expert1 | 1.52 | 1.49 | 1.21 | 1.41 |
| Non-expert2 | 1.45 | 1.57 | 3.38 | 2.13 |
| Non-expert3 | 1.18 | 0.89 | 1.52 | 1.20 |
| Non-expert4 | 1.18 | 0.60 | 2.82 | 1.53 |

##### 4.2.2.2 Rotational Errors

In the *R*_min,_ condition, the predominant rotational error was observed around the X-axis, ranging from 1.21° to 1.58°, while errors around the Y- and Z-axes remained consistently below 0.5°. For *R*_avg_, X-axis errors ranged from 0.51° to 1.70°, with most Y- and Z-axis deviations staying under 0.5°, except for a single instance where the error reached 0.96°. The non-expert group demonstrated the highest variability in *R*_max_ across all conditions, with X-axis errors fluctuating between 0.70° and 3.51°, and Y-axis errors ranging from 0.82° to 1.31°. Z-axis deviations were minimal, consistently remaining below 0.2°. These results complement the RMSD analysis, indicating that non-experts struggled to achieve proper AC-PC alignment in highly rotated images. However, with few exceptions, their accuracy across all three rotation conditions was generally comparable to that of the expert group (Table 5).

**Table 5.** Rotation angles (in degrees) for all three axes, comparing the baseline with non-expert assessments. Additionally, it shows the absolute difference values (in brackets) between the non-experts’ measurements and the baseline.

| | $R_{\min}$ | | | $R_{\text{avg}}$ | | | $R_{\max}$ | | |
| --- | --- | --- | --- | --- | --- | --- | --- | --- | --- |
| | $X$ | $Y$ | $Z$ | $X$ | $Y$ | $Z$ | $X$ | $Y$ | $Z$ |
| *Baseline* | -11.65 | -0.35 | 0.09 | 8.54 | 3.48 | 2.26 | 27.27 | -0.74 | -4.58 |
| Non-expert1 | -10.07<br>(1.58) | -<br>0.25(0.11) | 0.28(0.19) | 9.95 (1.41) | 3.38(0.10) | 3.22(0.96) | 26.57<br>(0.70) | 0.35(1.09) | -<br>4.55(0.03) |
| Non-expert2 | -<br>13.16(1.51) | -<br>0.34(0.01) | -<br>0.10(0.19) | 10.24(1.70) | 3.79(0.31) | 2.52(0.26) | 23.76(3.51) | 0.08(0.82) | -<br>4.54(0.03) |
| Non-expert3 | -<br>10.44(1.21) | -<br>0.40(0.04) | 0.41(0.32) | 7.62(0.92) | 3.77(0.30) | 2.41(0.15) | 26.32(0.95) | 0.57(1.31) | -<br>4.67(0.09) |
| Non-expert4 | -<br>10.45(1.21) | -<br>0.57(0.22) | 0.25(0.16) | 8.03(0.51) | 3.88(0.41) | 2.34(0.08) | 24.42(2.85) | 0.20(0.94) | -<br>4.71(0.14) |

#### 4.2.3 Performance of automated methods

##### 4.2.3.1 Root-mean-square deviation (RMSD)

RMSD analysis revealed a clear performance hierarchy among the automated methods (Table 6). Template registration that relied on the generic MNI template performed worst, with a mean RMSD of 5.89 mm across the three rotation conditions and errors that grew as the rotational offset increased. Substituting an AC–PC-pre-aligned template substantially improved accuracy, reducing the mean RMSD to 2.15 mm.

**Table 6:**
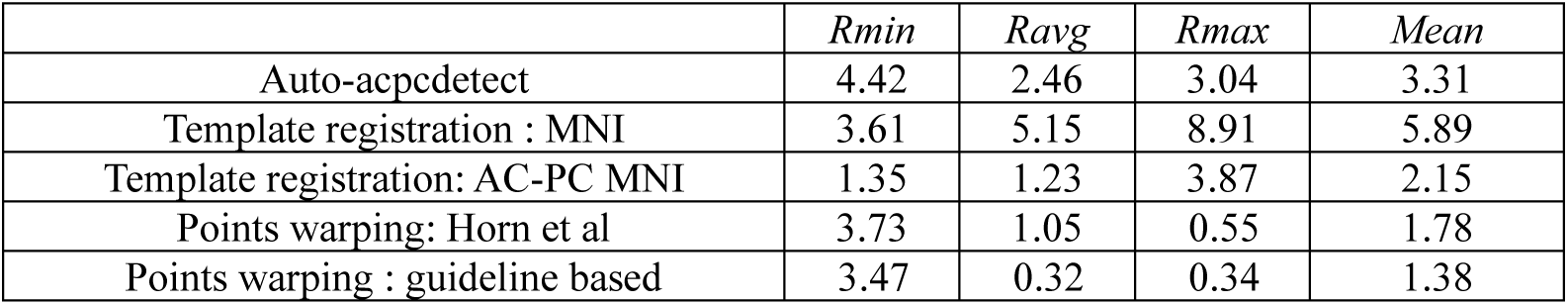
RMSD values (in mm) computed from rotation matrices, quantifying the deviation of automated method’s transformation from the baseline.

| | $R_{min}$ | $R_{avg}$ | $R_{max}$ | $Mean$ |
| --- | --- | --- | --- | --- |
| Auto-acpcdetect | 4.42 | 2.46 | 3.04 | 3.31 |
| Template registration : MNI | 3.61 | 5.15 | 8.91 | 5.89 |
| Template registration: AC-PC MNI | 1.35 | 1.23 | 3.87 | 2.15 |
| Points warping: Horn et al | 3.73 | 1.05 | 0.55 | 1.78 |
| Points warping : guideline based | 3.47 | 0.32 | 0.34 | 1.38 |

The Auto-acpcdetect algorithm achieved intermediate results (mean = 3.31 mm), but its error peaked at 4.42 mm in the *R*_min_ condition. Among the fully automated techniques, the point-warping method delivered the most precise AC–PC realignment overall. Even so, both the Horn et al. algorithm and the guideline-based procedure performed nearly identically, with only minor differences. Notably, the guideline-based approach emerged as the top performer of all automated methods, attaining the lowest mean RMSD of 1.38 mm.

##### 4.2.3.2 Rotational Errors

For the automatic “acpcdetect” method, substantial rotational errors were observed around the X-axis, with deviations of 4.31°, 2.50°, and 3.01° for *R*_min_, *R*_avg_, and *R*_max_ respectively (Table 7). Errors were also notable around the Y-axis, with deviations of 1.66° and 1.23° for the *R*_min_ and *R*_max_ conditions, respectively, and around the Z-axis, with a deviation of 1.24° for *R*_min_ indicating broader alignment inaccuracies across all axes.

**Table 7.** Rotation angles (in degrees) for all three axes, comparing the baseline with automated methods. Additionally, it shows the absolute difference values (in brackets) between the automated measurements and the baseline.

|  | *Rmin* |  |  | *Ravg* |  |  | *Rmax* |  |  |
| --- | --- | --- | --- | --- | --- | --- | --- | --- | --- |
|  | *X* | *Y* | *Z* | *X* | *Y* | *Z* | *X* | *Y* | *Z* |
| **Baseline** | **-11.65** | **-0.35** | **0.09** | **8.54** | **3.48** | **2.26** | **27.27** | **-0.74** | **-4.58** |
| Auto<br>acpcdetect | -15.96(4.31) | -2.02(1.66) | 0.65(0.56) | 6.04(2.50) | 3.57(0.10) | 1.02(1.24) | 24.26(3.01) | -1.97(1.23) | -4.14(0.44) |
| Template<br>registration : MNI | -15.41(3.76) | -0.21(0.14) | 0.53(0.44) | 2.93(5.61) | 4.41(0.93) | 2.29(0.03) | 17.86(9.41) | -0.13(0.60) | -2.95(1.62) |
| Template<br>registration: AC-PC<br>MNI | -11.15(0.50) | -1.11(0.76) | 1.25(1.17) | 7.28(1.26) | 3.53(0.05) | 2.72(0.45) | 23.57(3.70) | -1.14(0.40) | -2.53(2.04) |
| Points warping:<br>Horn et al | -7.93(3.72) | 0.14(0.49) | -1.08(1.16) | 8.87(0.34) | 4.54(1.06) | 1.93(0.34) | 27.41(0.14) | -1.30(0.57) | -4.66(0.08) |
| Points warping :<br>guideline based | -8.01(3.64) | -0.41(0.06) | 0.01(0.08) | 8.67(0.14) | 3.80(0.32) | 2.28(0.02) | 27.11(0.16) | -0.85(0.12) | -4.24(0.33) |

In comparison, the template registration method using the MNI 2007 template initially resulted in the highest rotational errors among the automated approaches. Specifically, the X-axis rotation errors were 3.76°, 5.61°, and 9.41° for *R*_min_, *R*_avg_, and *R*_max_ respectively. While most Y-and Z-axis errors remained below 1°, an exception was noted in the Rmax condition where the Z-axis error reached 1.62°. However, when the MNI 2007 template was first realigned to the AC–PC plane, these rotational discrepancies were significantly reduced. Post-realignment, X-axis errors dropped to 0.50°, 1.26°, and 3.70° for *R*_min_, *R*_avg_, and *R*_max_ respectively. Y-axis deviations remained consistently under 1° across all conditions, while minor Z-axis deviations of 1.17° and 2.04° were observed in the *R*_min_ and *R*_max_ conditions, respectively.

The point warping methods demonstrated the lowest rotational errors overall. The method based on Horn et al. yielded an X-axis error of 3.72° and a Z-axis error of 1.16° in the *R*_min_ condition. In the *R*_avg_ condition, the only notable deviation above 1° occurred around the Y-axis (1.06°), while all *R*_max_ errors remained below 1°. The guideline-based point warping method achieved the highest accuracy among the automated AC–PC realignment techniques. The only rotational error exceeding 1° occurred around the X-axis in the *R*_min_ condition where the errors reached to 3.64°. These rotational errors reflecting the results of RMSD values, demonstrating that the guideline-based point warping method outperforms all other automated technique

## 5 Discussion

We have proposed a comprehensive guideline for realigning structural MR images to the clinically accepted definition of the AC-PC plane. Our contribution introduces anatomical and imaging characteristics of the AC, PC, and four midline landmarks (i.e. the superior pontine notch, the fastigium of the cerebellum and two points, anterior and posterior, in the falx cerebri). Our guideline involves a step-by-step approach tailored for T1-weighted structural images, ensuring accurate alignment to the AC-PC plane. Additionally, we outline criteria to assess the quality of AC-PC alignment, which are included as the final step in our guideline but could also be used separately to evaluate the results of existing realignments.

We present our protocol in a structured format that guides users through each step required to define the AC–PC line and the mid-sagittal plane, ensuring an accurate AC–PC realignment in 3D Slicer. Although optimized for that software environment, these instructions can be applied in any setting: they describe how to (i) define the clinically accepted AC–PC line, (ii) establish the midplane using 4 midline points, and (iii) verify the resulting AC–PC alignment.

This guideline represents an enhanced iteration of the precursor version used during the evaluation phase. Drawing on insights gained from that initial assessment, it incorporates several refinements. A detailed account of all modifications can be found in Supplementary Material, Section 4 (“Amendments”). In summary, the current edition provides clearer instructions on precise AC–PC line definition and on verification of the final alignment.

The results from the rotational error analysis indicate that manual observers were generally accurate, though minor deviations were observed. Notably, the most significant errors among non-experts were recorded by non-expert 2 and non-expert 4, with rotational deviations around the X-axis measuring 3.51 and 2.85 degrees, respectively. Similarly, in the expert group, Expert 1 showed the largest errors, with deviations of 2.13 and 1.79 degrees around the X-axis. These discrepancies primarily stem from subtle differences in interpreting the AC-PC line relative to the baseline. Given that the AC and the PC are only approximately 23 mm apart on average (Schaltenbrand et al., 1977), even minor misalignments in defining the AC-PC line can lead to noticeable alignment deviations (Ardekani & Alzheimer’s Disease Neuroimaging, 2022; Evans et al., 1992). Highlighting the importance of accurate identification of AC and PC. (Supplementary figures 21-24)

The overall results reveals that the experts performed comparably in AC-PC realignment, both with and without the guideline, highlighting the parallels between clinical practice and the proposed guideline. Additionally, non-experts consistently achieved expert-level accuracy in AC-PC realignment across various rotational conditions, with only a few exceptions. This highlights the potential to be highly beneficial in training non-expert individuals with little to no experience in neuroanatomy and neuroimaging, enabling them to achieve expert-level accuracy in realigning and validating structural MRI images to the clinically utilized AC-PC plane. This capability of our proposed guideline can bridge the gap between expert knowledge and non-expert application, making high-precision alignment attainable for a broader range of users. Furthermore, the guideline can serve as a standard protocol for experts, helping to minimize inter-observer variability, which often poses a challenge (Pallavaram et al., 2008), especially when high-quality imaging is not achievable. For example, during the validation process of the precursor guideline, experts performed AC-PC realignment in two scenarios: one with the guideline and one without. In the condition without the guideline, when experts were required to select at least three midline points, there was no correspondence among their selections, as their choices varied randomly along the midline. By providing a systematic approach, the guideline effectively addresses this issue, promoting accuracy and consistency in the identification process.

A comparative analysis of automated methods revealed that warping landmark points from the template to individual space, followed by realignment within that space, outperformed all other automated approaches. Among the two point-warping techniques, the proposed guideline-based approach proved to be more effective than the method adopted from Horn et al., although the differences were minimal and practically negligible. Furthermore, the findings from the template registration method underscored the importance of selecting an appropriate template to accommodate variations in AC-PC line definitions. The largest deviations occurred in template-based registration method using the MNI template were, X-axis *R*_min_ =3.76, *R*_avg,_ = 5.61, *R*_max_ = 9.41 degrees. evidently these largest errors were reduced to *R*_min_ =0.50, *R*_avg,_ = 1.26, *R*_max_ = 3.70 degrees when the MNI template were realigned to the AC-PC plane. These results clearly demonstrate that registering an image to the standard MNI152 template is not a suitable approach if the objective is to align the image with the clinically relevant AC-PC plane or the Talairach version of the AC-PC plane. Finally, the results from fully automated AC-PC realignment methods highlighted the limitations of model-based approaches. For instance, the observed rotational errors *R*_min_ = 4.31°, *R*_avg_ = 2.50°, and *R*_max_ = 3.01° likely due to the inaccurate localization of the AC and PC landmarks. This inaccuracy, in turn, compromised the precision of the final image realignment (Supplementary figures 18-20).

Despite being the preferred choice for large datasets, automated methods can exhibit a wide range of variation in both how they achieve AC-PC realignment and how they define the AC-PC space itself. It is crucial to recognize these differences, particularly regarding the definition of the AC-PC line across various atlas definitions (Horn et al., 2017; Nowinski, 2001). As demonstrated in our precursor guideline validation, even a small deviation in the AC-PC line definition can lead to significant errors, potentially resulting in false interpretations. Additionally, it is important to acknowledge the limitations of these automated methods, which include algorithm-specific constraints inherent to both registration-based and model-based approaches, such as registration failures and inaccuracies in identifying landmarks (Klein et al., 2009; Liu & Dawant, 2014). Our proposed guideline can be beneficial in such instances to validate results from automated methods, manually realign images when these methods fail, or prepare templates by either realigning the template to the AC-PC plane or identifying landmarks in the initial template for warping to individual space when using a registration approach. This capability is extremely advantageous for developing new automated AC-PC realignment tools, which rely on extensive ground truth images typically provided by experts. Additionally, it can be further extended to create ground truth for any proposed landmarks, such as the fastigium or superior pontine notch, thereby facilitating the development of new automated tools for both clinical and research purposes (Lau et al., 2019; Oba et al., 2005; Roelants et al., 2016).

To the best of our knowledge, no published literature currently provides a comprehensive guide on the anatomical landmarks required for AC-PC realignment and a step-by-step process for aligning structural MR images to the AC-PC plane. It is worth noting that while many software packages that utilize AC-PC alignment offer a brief introduction and visual aids to help identify the AC, PC, and, in some cases, the midline points, these resources often lack comprehensive detail. For example, the neuroanatomical and neuroimaging properties of proposed landmarks and variations in the definition of the AC-PC line.

Previous efforts related to AC-PC realignment protocols are noteworthy. One significant contribution comes from (Ramirez et al., 2014) who introduced a standardized, video-guided protocol for semi-automatic brain region extraction that includes a step for aligning images to the AC-PC plane. Although AC-PC realignment was not the primary objective of their study, the accompanying video and text instructions can be effectively used for manually realigning structural images to this plane. Additionally, their work outlines a set of validation criteria that we initially adopted in our precursor guideline. Another important study is that of (Hamilton et al., 2017) where the authors provide brief instructions for manual AC-PC realignment, which they used to standardize individual brain images, for subsequent processing and analysis. A final example is the study by Lau et al (Lau et al., 2019) where the authors developed a method for evaluating the correspondence of brain images using 32 anatomical fiducials, including the AC, PC, and various midline structures. While AC-PC realignment is not the primary focus of their work, the method can still be employed to realign images to the AC-PC plane.

While we acknowledge these efforts in providing instructions for manual AC-PC realignment, we also recognize their limitations, particularly in assuming prior knowledge of anatomical and imaging characteristics of the AC, PC, and the various definitions and purposes of AC-PC realignment. This can present a significant barrier for novice observers. The challenge arises because AC-PC realignment is not the primary focus of these studies but rather a minor component within a broader workflow. Our guideline addresses these gaps by providing a standardized approach to identifying anatomical structures crucial for AC-PC transformation in structural imaging, ensuring clarity and reliability in the process.

## Acknowledgement and competing interest

KVH gratefully acknowledges funding from the Dutch Research Council (NWO; VIDI grant no. 09150171910043). RSV is an independent consultant for Boston scientific and Abbott and received Research grants from Boston scientific. C.F.B. gratefully acknowledges funding from the Well-come Trust Collaborative Award 215573/Z/19/Z and the Netherlands Organization for Scientific Research Vici Grant 17854. C.F.B is director and shareholder in SBGneuro Ltd.

While preparing this article, the authors used ChatGPT (OpenAI) to refine the style and grammar of portions of the text and to assist with code formatting/cleanup for Python scripts used in the data analysis. The tool was not used to generate new scientific content, results, or conclusions. All suggested edits were reviewed and verified by the authors, who take full responsibility for the content of the publication.

