## Supplementary Material for "ACPC-MAP: A protocol for manually aligning structural T1-weighted Magnetic Resonance Images to the Anterior Commissure – Posterior Commissure plane according to clinical standards"

#### Contents


1.Precursor Guideline

2.Verification Criteria for the Precursor Guideline

3.Comprehensive Evaluation of the Precursor Guideline

4.Amendments

5.Examples of Minor Rotational Deviations

### 1.Precursor Protocol

#### 1.1.Anterior Commissure(AC):

1: Identify the Anterior Commissure (AC)

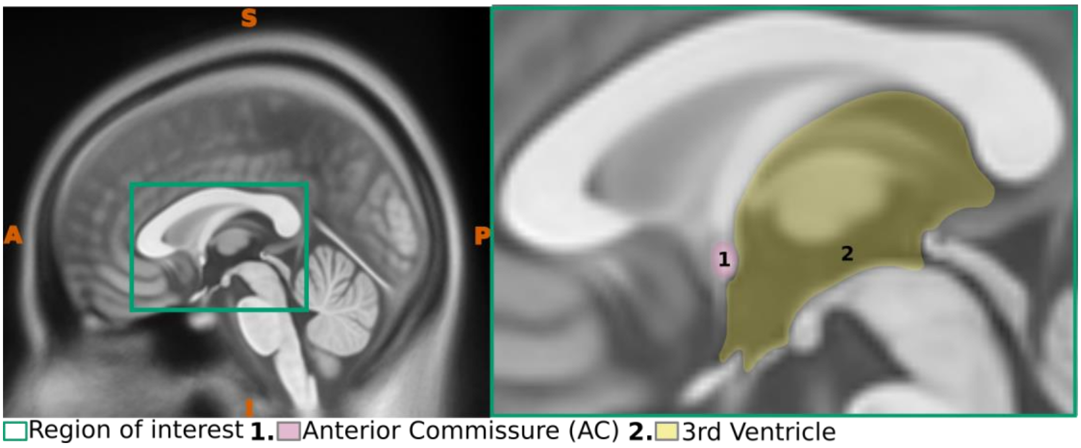


**Figure. 1** Sagittal view of the Anterior commissure. The left panel presents a midsagittal view with the region of interest (ROI) highlighted, whereas the right panel offers an enlarged view of the ROI, emphasizing critical anatomical structures

2: Select a point at the posterior border of the Anterior Commissure (AC)

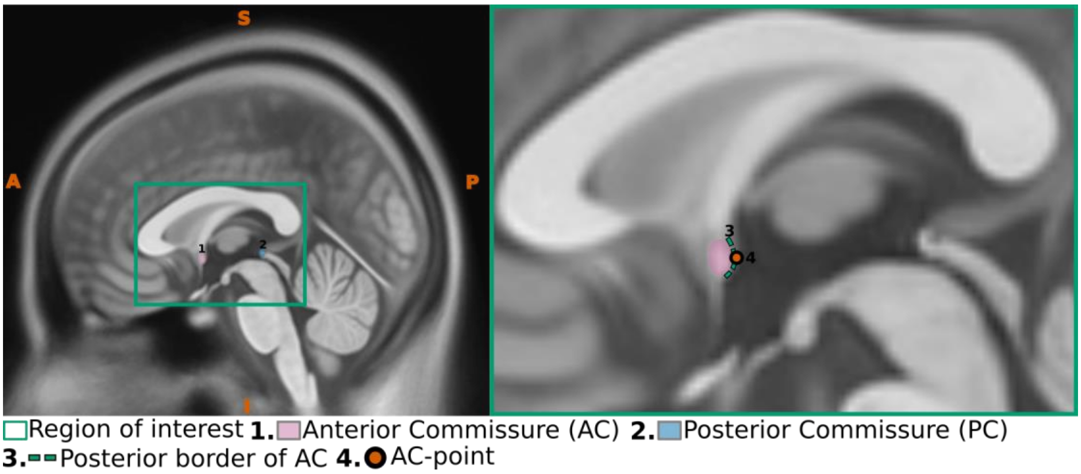


**Figure. 2** Sagittal view of the Anterior commissure. The left panel presents a midsagittal view with the region of interest (ROI) highlighted, whereas the right panel offers an enlarged view of the ROI, emphasizing critical anatomical structures

3. Use the corresponding axial plane to ensure the annotated point is accurately positioned along the midline.

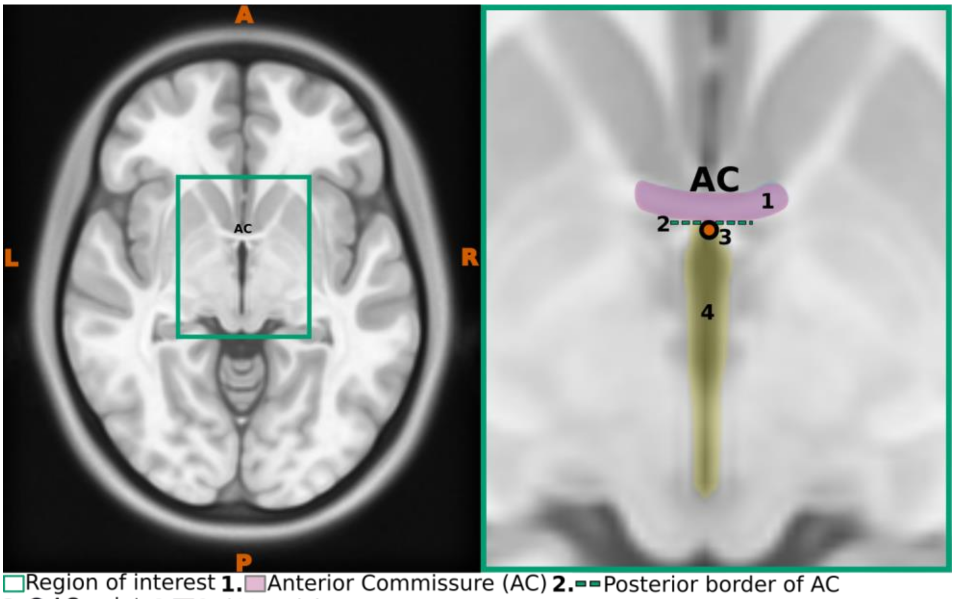


□Region of interest **1.**□Anterior Commissure (AC) **2.**▬▬Posterior border of AC
**3.**●AC-point **4.**□3rd Ventricle

**Figure. 3** Axial view of the Anterior commissure. The left panel presents a axial view with the region of interest (ROI) highlighted, whereas the right panel offers an enlarged view of the ROI, emphasizing critical anatomical structures

#### 1.2.Posterior Commissure(PC):

1.Identify the Posterior commissure (PC)

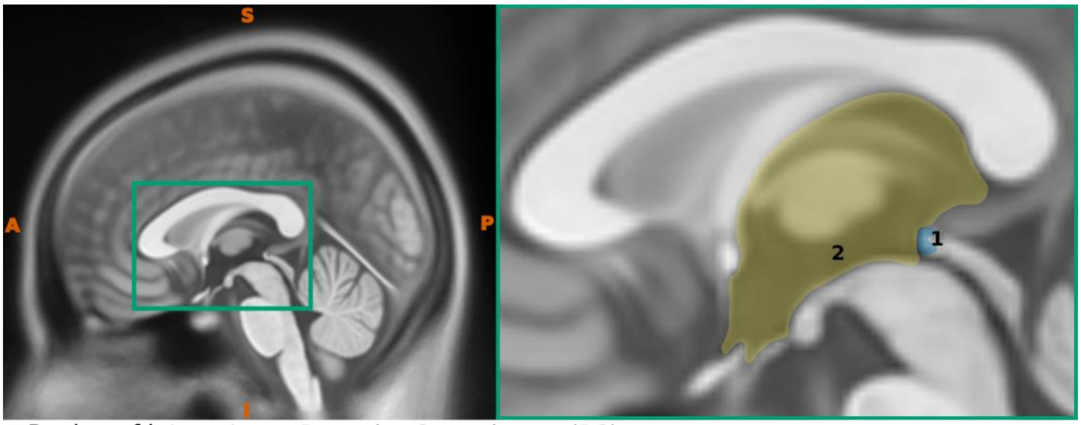


□Region of interest **1.**□Posterior Commissure (PC) **2.**□3rd Ventricle

**Figure. 4** Sagittal view of the Posterior commissure. The left panel presents a midsagittal view with the region of interest (ROI) highlighted, whereas the right panel offers an enlarged view of the ROI, emphasizing critical anatomical structures

2. Select a point at the Anterior border of the Posterior Commissure (PC)

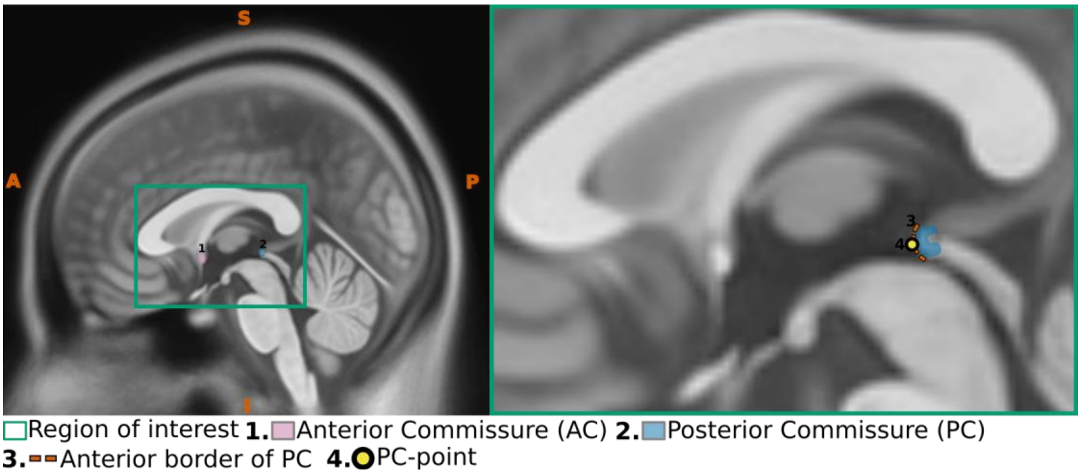


**Figure. 5** Sagittal view of the Posterior commissure. The left panel presents a midsagittal view with the region of interest (ROI) highlighted, whereas the right panel offers an enlarged view of the ROI, emphasizing critical anatomical structures

3. Use the corresponding axial plane to ensure the annotated point is accurately positioned along the midline

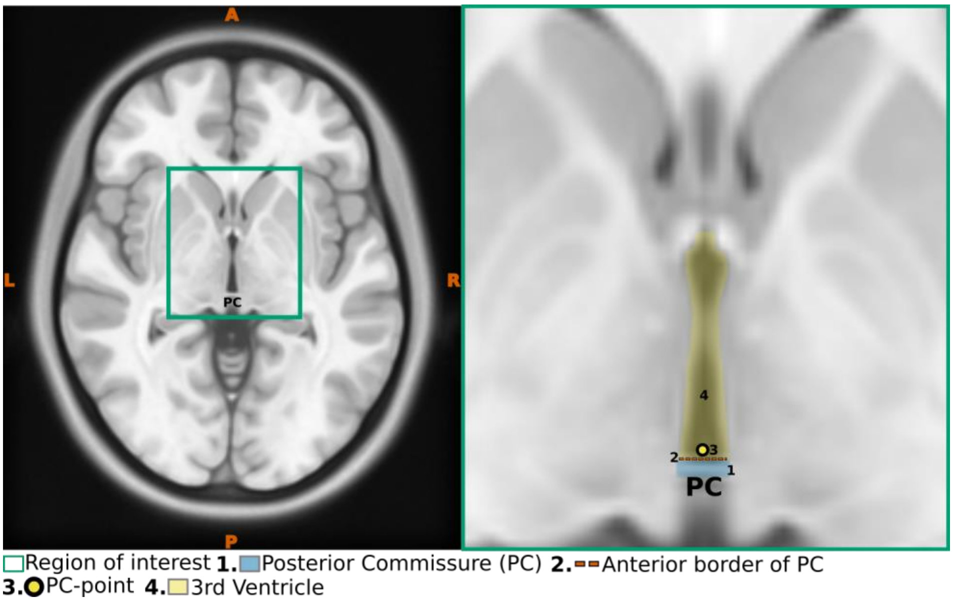


**Figure.6** Axial view of the Posterior commissure. The left panel presents a axial view with the region of interest (ROI) highlighted, whereas the right panel offers an enlarged view of the ROI, emphasizing critical anatomical structures

##### 1.3.Superior Pontine notch:

1. Identify the superior pontine notch
2. Place the first midline point in the superior pontine notch at the junction where the midbrain and pons meet

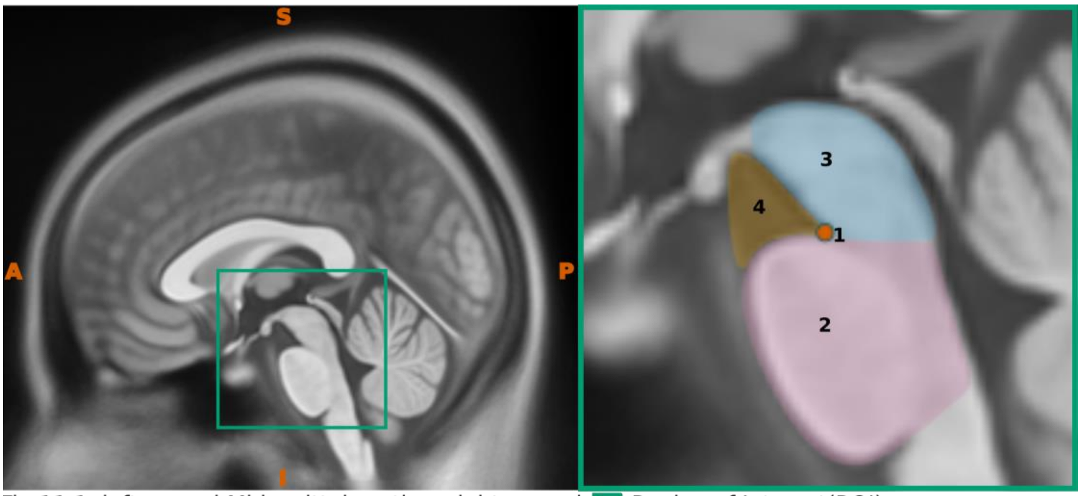


Fig.11.1, left pannel:Midsagittal section, right pannel: Region of Interest(ROI)
**1.** Superior Pontine notch **2.** Pons **3.** Midbrain **4.** Interpeduncular fossa

**Figure. 7** Sagittal view demonstrating the superior pontine notch. The left panel depicts a midsagittal perspective highlighting the region of interest (ROI), while the right panel provides a magnified view of the ROI, clearly emphasizing critical anatomical structures and specifically highlighting the superior pontine notch

3. Use the corresponding axial slice to ensure the point is correctly positioned in the midline, where the two (left and right) peduncles join to form the interpeduncular angle

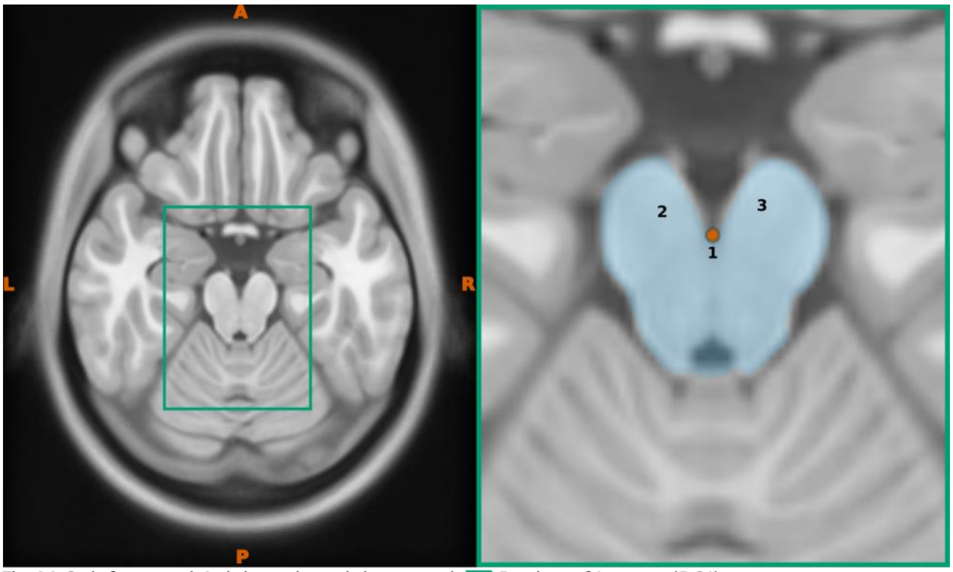


Fig.11.2, left pannel:Axial section, right pannel: Region of Interest(ROI)
**1.** Superior Pontine notch **2.** Left cerebral peduncle **3.** Right cerebral peduncle

**Figure. 8** Axial view demonstrating the superior pontine notch. The left panel depicts a axial perspective highlighting the region of interest (ROI), while the right panel provides a magnified view of the ROI, clearly emphasizing critical anatomical structures and specifically highlighting the superior pontine notch

##### 1.4. Fastigium or Apex of the Roof of fourth ventricle:

1. Identify the fastigium, also known as the apex of the roof of the fourth ventricle
2. Place the second midline point in the fastigium

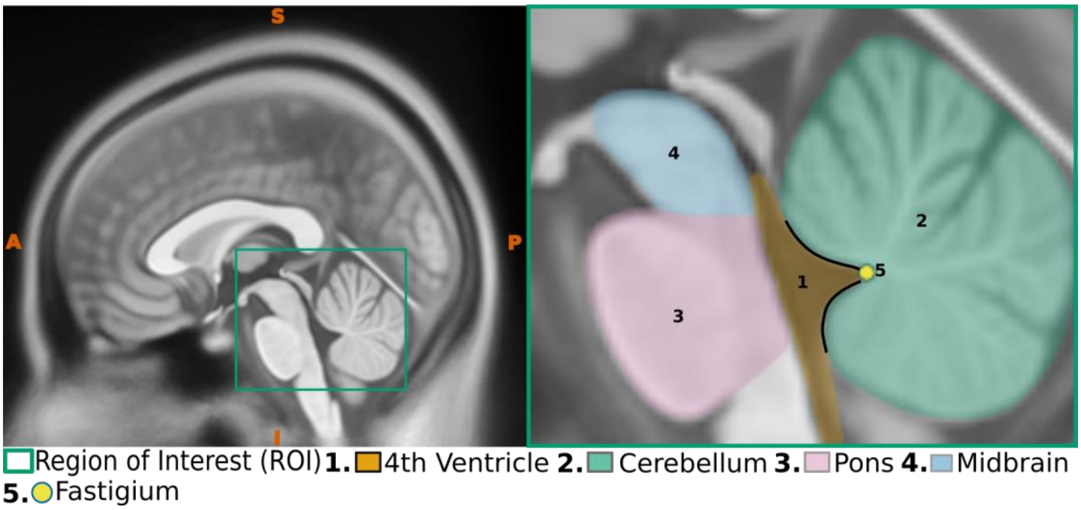


**Figure. 9** Sagittal view demonstrating the fastigium. The left panel depicts a midsagittal perspective highlighting the region of interest (ROI), while the right panel provides a magnified view of the ROI, clearly emphasizing critical anatomical structures and specifically highlighting the fastigium

3. Use the corresponding axial slice to ensure the point is accurately placed along the midline in the fastigium

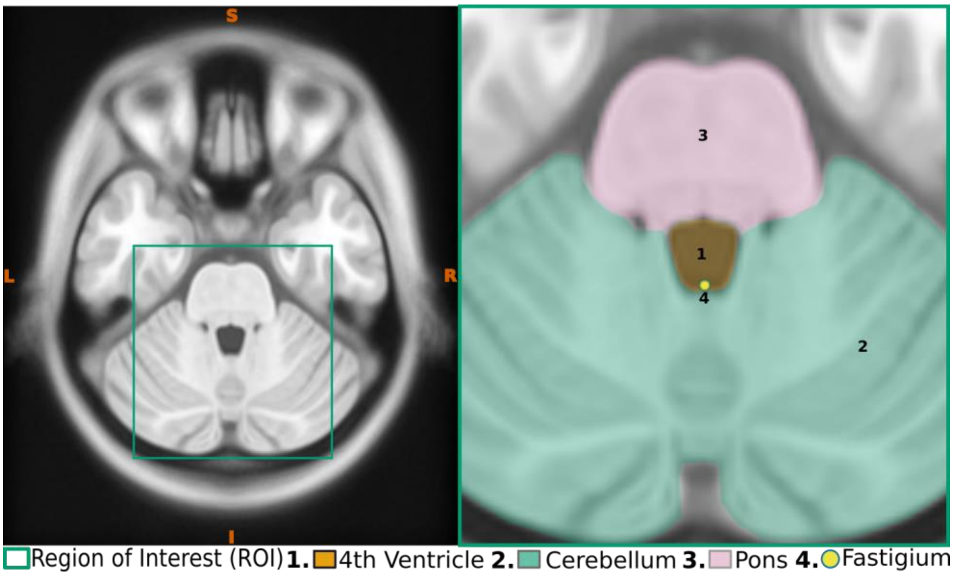


**Figure. 10** Axial view demonstrating the fastigium. The left panel depicts a axial perspective highlighting the region of interest (ROI), while the right panel provides a magnified view of the ROI, clearly emphasizing critical anatomical structures and specifically highlighting the fastigium

#### 1.5. Anterior and Posterior Points in the Falx Cerebri

1. Identify the corpus callosum.
2. Draw a line at the highest point of the upper border of the corpus callosum.

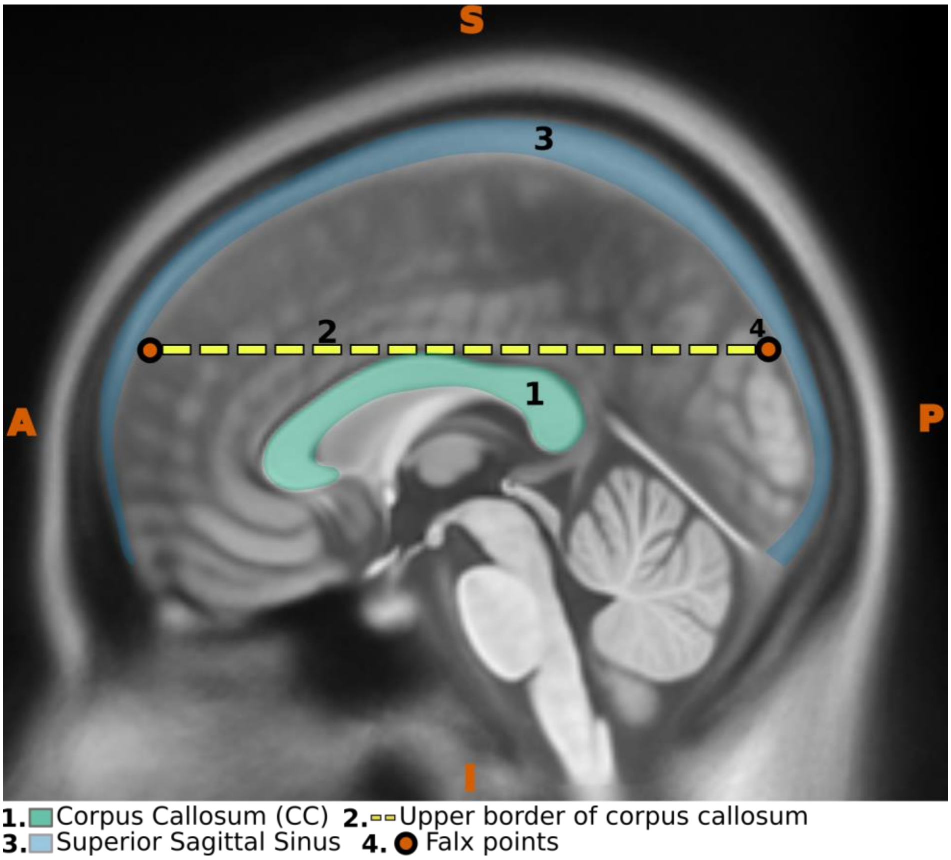


**Figure. 11** Sagittal view illustrating the corpus callosum with a horizontal line marking its superior-most boundary

3. Use the corresponding axial plane to place two midline points (the 3rd and 4th points): one at the anterior end of the falx cerebri (refer to Figure.12) and another at the posterior end, where the falx cerebri splits to enclose the superior sagittal sinus

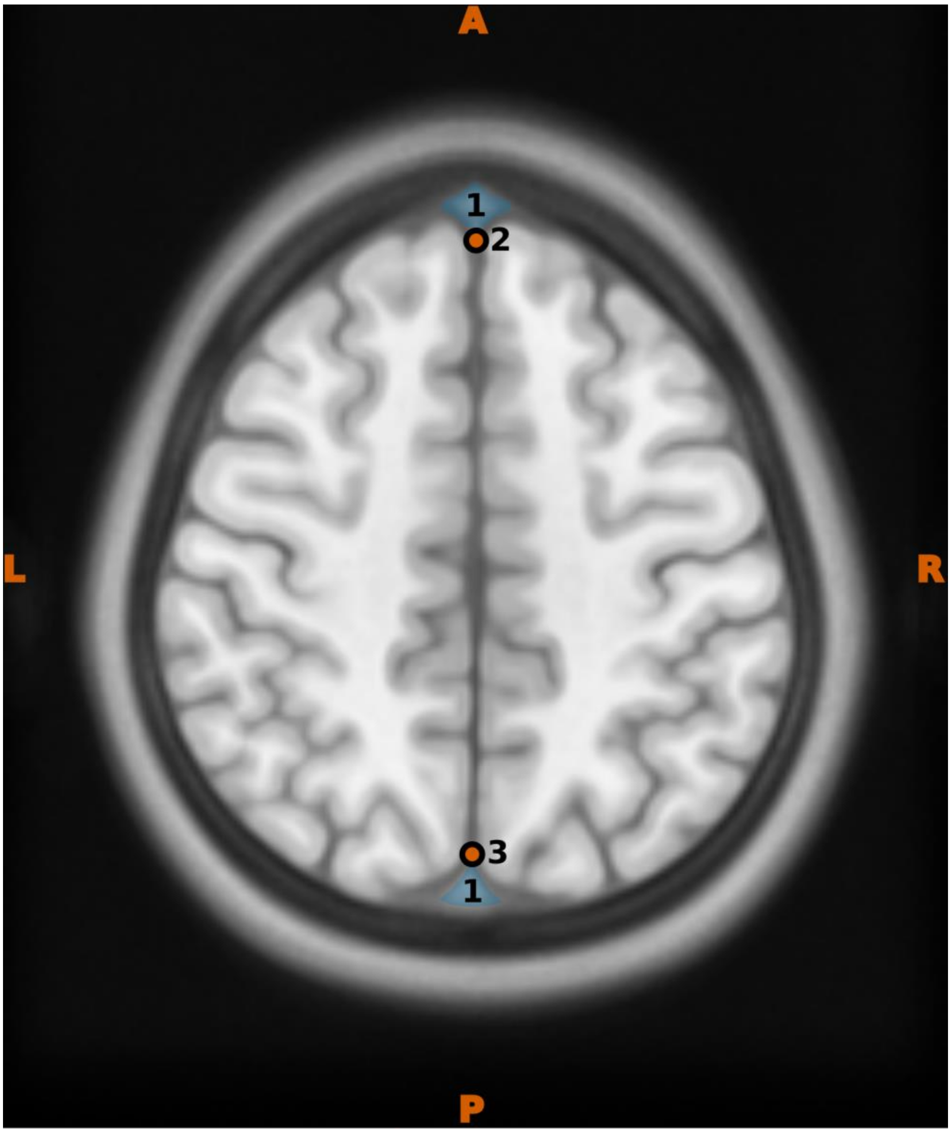


**Figure. 12** Axial view demonstrating the anterior and posterior falx cerebri at the superior-most level of the corpus callosum

#### 2.Verification of AC_PC alignment

After realigning the structural image to the AC-PC plane, it is essential to verify that the image is properly aligned. This involves checking for any rotations in the X, Y, and Z Axes, commonly referred to as pitch, roll, and yaw.

##### 2.1.Verifying pitch:

1.After the successful realignment, the Anterior Commissure (AC) and the Posterior Commissure (PC) should be at the same level. Refer to Figure. 13 For the differences before and after AC-PC alignment

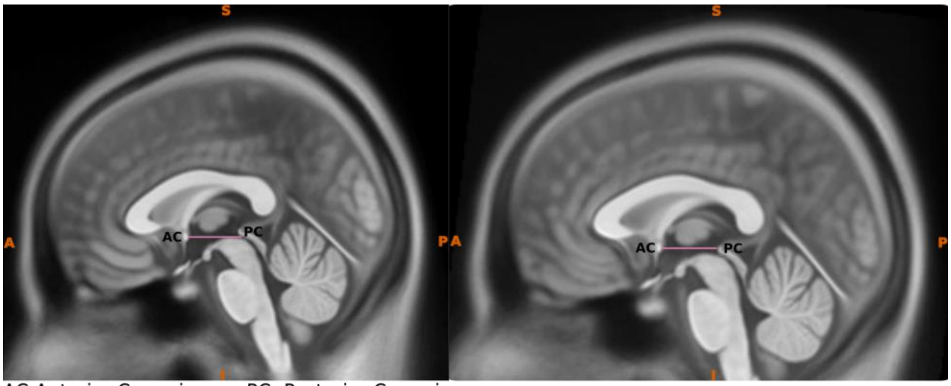

AC-Anterior Commissure, PC- Posterior Commissure

**Figure. 13** Verification of Pitch. From left to right, the first panel displays the midsagittal plane with the positions of the AC and the PC following improper realignment, highlighting noticeable misalignment. The right panel shows the same plane after successful realignment, with the AC and PC properly aligned at the expected anatomical level.

2.The presence of the Anterior Commissure (AC) and the Posterior Commissure (PC) in the same axial slice will create the characteristic keyhole appearance of the third ventricle. Refer to Figure.14 for the difference before and after the AC-PC alignment, and note the keyhole appearance after the AC-PC alignment (Figure 14 right panel)

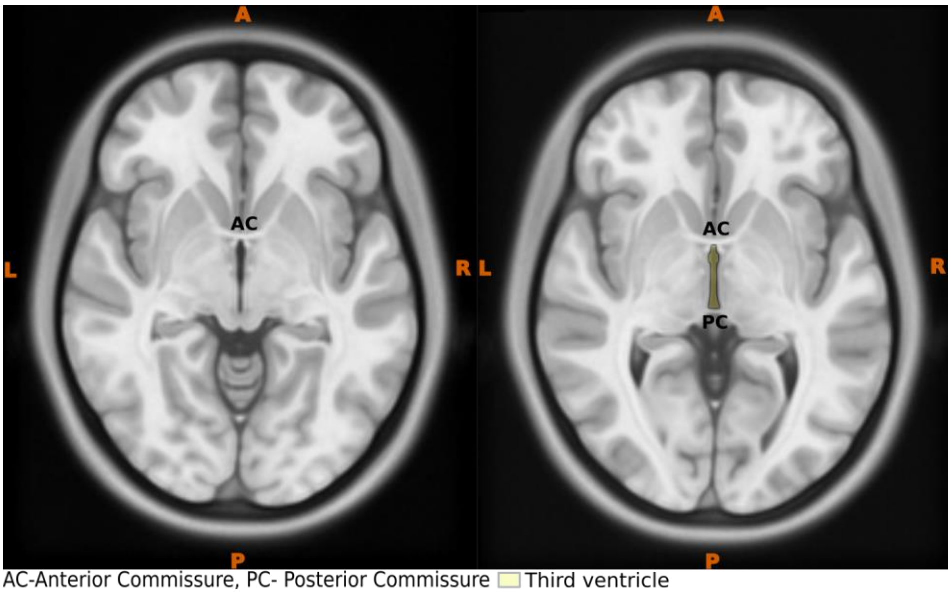

AC-Anterior Commissure, PC- Posterior Commissure Third ventricle

**Figure. 14** Verification of Pitch . Axial planes illustrating the appearance of the AC and PC before (left panel) and after (right panel) successful AC-PC realignment. In the right panel, the simultaneous presence of both AC and PC within the same slice confirms proper realignment. Additionally, the characteristic keyhole shape of the third ventricle further supports accurate anatomical alignment.

#### 2.2.Verifying roll:

1. Roll verification can be performed by examining the appearance of the eyeballs in axial images. In a correctly realigned scan, the eyeballs should appear symmetrical and evenly shaped within the selected axial slice. In Figure 15, note the uneven appearance of the eyeballs in the

left panel, indicating misalignment, compared to the right panel, where their even, symmetrical appearance confirms successful AC-PC realignment.

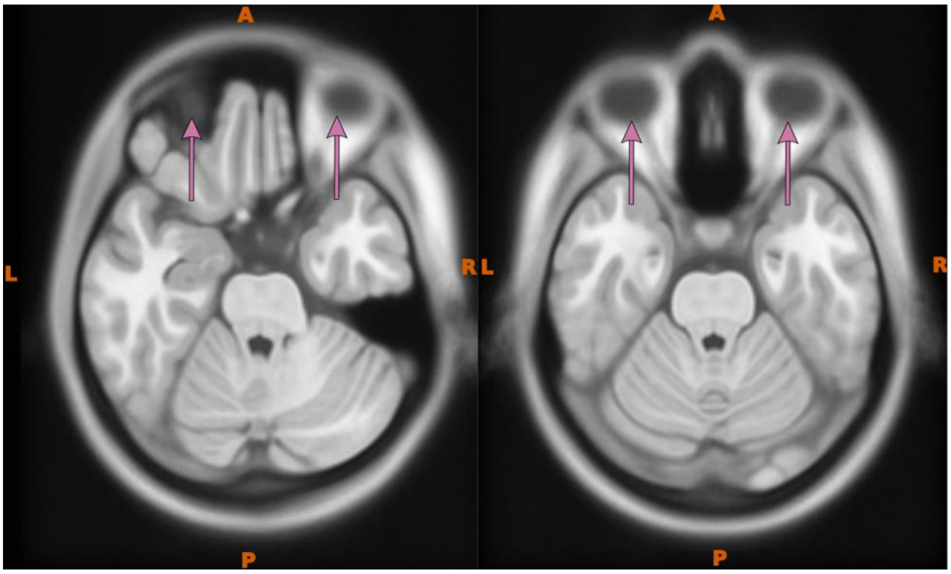


**Figure 15**: Verification of Roll. Axial planes illustrating the appearance of the eyeballs before (left panel) and after (right panel) successful AC-PC realignment.

#### 2.3.Verifying yaw:

1. We can verify the yaw by placing the vertical crosshair. The vertical line should align with the longitudinal fissure or midline of the brain in the axial plane. Refer to Figure. 16 left Panel where the longitudinal fissure and vertical line(yellow line) are not properly aligned before the AC-PC alignment. In contrast, Right Panel demonstrates proper alignment of the longitudinal fissure with the vertical line(yellow line), indicating successful midline alignment

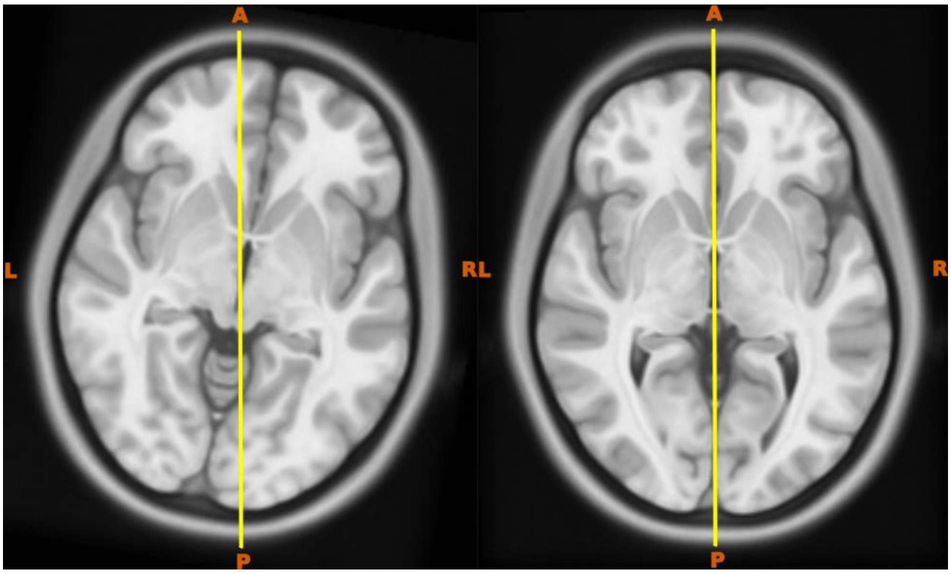


**Figure. 16** Verification of Yaw. Axial planes illustrating midline deviations before (left panel) and after (right panel) successful AC-PC realignment. Note how the midline shift in the left panel is corrected in the right panel, confirming proper alignment

### 3. Comprehensive validation of precursor guideline

#### 3.1.1. Data preparation

Two structural T1-weighted images were randomly selected from the open source ABRIM (Jansen et al., 2024) dataset: one representing healthy young adults (18-35 years) and another representing healthy aged (60-75). In addition, we randomly selected a representative T1-weighted image from a patient with Parkinson's Disease from the POM dataset. The three selected images were cropped to remove the large neck region using a robust field of view (FOV) method, and subsequently underwent bias correction (Zhang et al., 2001) utilizing the fsl_anat tool from the FSL software package (Jenkinson et al., 2012; Tian et al., 2016). Following this, the bias-corrected images were aligned to the AC-PC plane using the proposed guideline by the first author (AS). Once aligned, these images were rotated to three different angles. These rotation angles were determined from 226 ABRIM subjects, using the linear registration matrix associated with those subject's transformation to the MNI 152 space. Among the 266 rotations, three key values minimum, mean, and maximum were selected for image rotation. These angles were then randomly sign-flipped, resulting in the minimum value shifting to the mean position and the mean value shifting to the minimum. The resulting rotations, Rmin, Ravg, and Rmax, were then used to generate rotated images with known rotation parameters (Tables 1A and B). The rotated images were then subjected to intensity normalization using the percentile clipping method (i.e. removing the bottom and top 2%) see Figure.17 for the full data preparation pipeline.

Table 1: Summary of rotation angle values extracted from the Abrim dataset. The extreme values Table1A (minimum, mean, and maximum) were randomly sign-flipped and renamed as Rmin, Ravg, Rmax, according to new (minimum, mean, and maximum) values Table 1B .

**Table 1 A**

| | x | y | z |
|---|---|---|---|
| count | 226 | 226 | 226 |
| **mean** | **10.0** | **-0.0** | **0.0** |
| std | 4.0 | 1.0 | 1.0 |
| **min** | **-8.0** | **-4.0** | **-2.0** |
| 25% | 8.0 | -1.0 | -0.0 |
| 50% | 10.0 | -0.0 | 0.0 |
| 75% | 12.0 | 0.0 | 1.0 |
| **max** | **25.0** | **4.0** | **3.0** |

**Table 1 B**

| | x | y | z |
|---|---|---|---|
| Rmin | -10.0 | 0.0 | 0.0 |
| Ravg | 8.0 | -4.0 | 2.0 |
| Rmax | 25.0 | 4.0 | -3.0 |

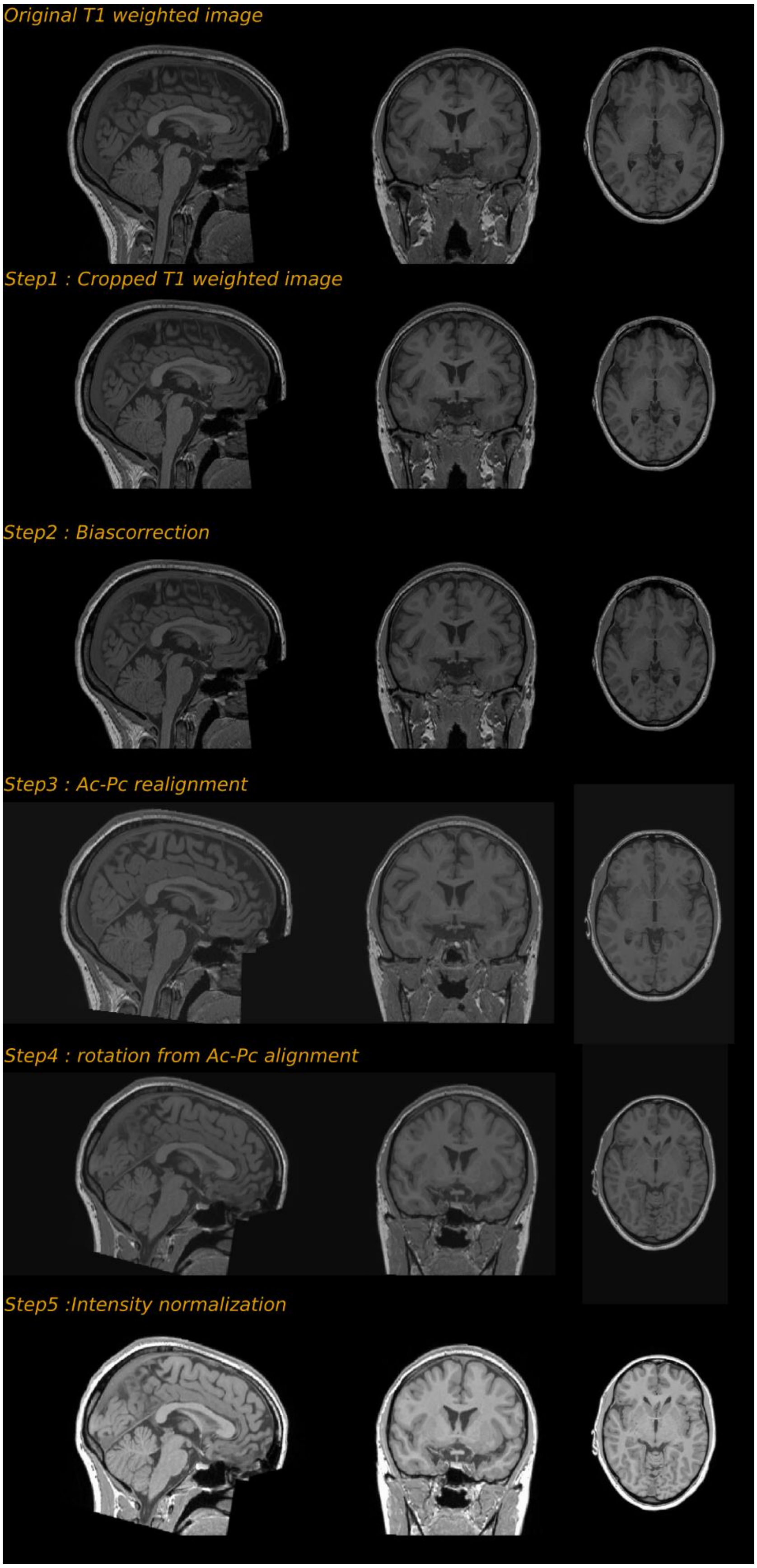


**Figure. 17** Preprocessing Stages for Data Preparation. This figure illustrates the key preprocessing steps involved in preparing the data for the evaluation process.

##### 3.1.2. Experimental procedures

The evaluation process comprised two stages. In the first stage, two expert and four non-expert volunteers conducted the realignment manually using the proposed guidelines. The experts were practicing neurosurgeons who perform manual AC-PC realignment daily. They conducted the realignment twice, first without, then with the guideline. The non-experts had no prior anatomical knowledge or experience with AC-PC realignment. They conducted the realignment with the proposed guideline only. All manual realignments were performed using 3D Slicer 5.0.3 (Fedorov et al., 2012). In the second stage, the realignment was executed using four existing automated methods.

The first author (AS) also manually realigned the three rotated images to the AC-PC plane following the proposed guidelines. This was done to establish a baseline performance of the guideline under similar rotation and imaging conditions, which is important because different rotation angles could differentially affect the visibility of the landmarks. The spatial locations of the borders of the AC and PC, and four midline points established by the volunteers and automated methods were compared to the baseline in terms of Euclidean distances. Additionally, image rotation angles and the Root Mean Squared Deviation (RMSD), derived from the rotation matrix, were compared against the baseline. Angular and spatial location errors were computed using a custom Python script, while the RMSD was calculated following the method outlined by Jenkinson,1999 (Jenkinson, 1999).

#### 3.2 Expert performance with and without the guideline

Two experts were tasked with realigning three rotated images ($R_{min}$, $R_{avg}$, and $R_{max}$) to the AC-PC Plane. First, the experts realigned the images without the aid of our guideline. They adhered only to the minimal requirements of the AC-PC transformation module in 3D Slicer software, which involves selecting the AC and PC points along with at least three midline points to create an AC-PC transformation. Next, the experts performed the AC-PC realignment using the proposed guideline. To ensure a balanced assessment, the order of the rotation conditions was reversed between the experts. We evaluated the experts' performance by measuring the RMSD, rotation angles, and Euclidean distances between the spatial locations of the AC, PC, and midline relative to the baseline.

##### 3.2.1. AC and PC location errors

Experts consistently demonstrated Euclidean distance errors of less than 1 mm across all rotation conditions, regardless of the presence of the guideline. Specifically, for $R_{min}$, the differences in AC and PC showed errors below 0.6 mm, which is smaller than the voxel size (0.8 mm). Similarly, the AC–PC distance error also remained below 0.6 mm. The average diameters of the commissures (3 mm for the AC and 2 mm for the PC in the sagittal plane), and the average AC-PC distance (24 mm) as reported by Schaltenbrand (Schaltenbrand et al., 1977).

For $R_{avg}$, experts achieved slightly better accuracy for the AC when using the guideline, whereas better results for the PC were observed without it. In both cases, the AC-PC distance error remained below 0.3 mm. Under the $R_{max}$ condition, performance was generally better without the guideline, with the exception of a single PC instance where the error reached 0.87 mm. Overall, experts demonstrated consistently high accuracy across all conditions, with no significant differences in performance between the guideline and no-guideline scenarios. These findings suggest that the proposed guideline closely aligns with the AC-PC selection strategies employed by expert neurosurgeons, highlighting strong similarities between established clinical conventions and the guideline's recommendations

Table 2. Comparison of Errors in AC-PC Location and AC-PC Distance. This table illustrates the errors (in mm) in AC-PC location and AC-PC distance, comparing results between experts and baseline. The comparison is made both with and without the use of guidelines

| | *Rmin* | | | *Ravg* | | | *Rmax* | | |
|---|---|---|---|---|---|---|---|---|---|
| | **Without guideline** | | | | | | | | |
| | **ac** | **pc** | **ac-pc distance** | **ac** | **pc** | **ac-pc distance** | **ac** | **pc** | **Ac-pc distance** |
| Expert 1 | 0.42 | 0.55 | 0.57 | 0.39 | 0.46 | 0.13 | 0.11 | 0.87 | 0.48 |
| Expert 2 | 0.56 | 0.20 | 0.26 | 0.75 | 0.71 | 0.14 | 0.21 | 0.48 | 0.49 |
| **mean** | **0.49** | **0.38** | **0.42** | **0.57** | **0.58** | **0.14** | **0.16** | **0.68** | **0.48** |
| | **With guideline** | | | | | | | | |
| Expert 1 | 0.32 | 0.58 | 0.31 | 0.50 | 0.76 | 0.34 | 0.44 | 0.84 | 0.98 |
| Expert 2 | 0.45 | 0.24 | 0.26 | 0.32 | 0.94 | 0.09 | 0.26 | 0.83 | 0.44 |
| **mean** | **0.38** | **0.41** | **0.29** | **0.41** | **0.85** | **0.22** | **0.35** | **0.84** | **0.71** |

##### 3.2.2. Midpoint location errors

The analysis of midpoint location error (i.e., Euclidean distances) was conducted exclusively under the guideline condition. Without the guideline, experts tended to select random points along the midline, resulting in a lack of consistency and correspondence. The differences in identifying the spatial locations for midpoint 1 are predominantly within 1 voxel (0.8mm) difference, with two exceptions where the errors reach up to 1 mm for $R_{\min}$ and 0.82 mm for $R_{\max}$. A similar trend is observed for midpoint 2, where most of the errors are less than 1 voxel difference, except for $R_{\min}$, which yielded an error of 0.93 mm. The errors increase for midpoints 3 and 4. For midpoint 3, the largest errors are observed for $R_{\max}$ (1.56 mm and 2.06 mm), while the minimal error occurs for $R_{\text{avg}}$ (0.79 mm and 1.05 mm). For $R_{\min}$, the error range is between 1.05 mm and 1.55 mm. The differences for midpoint 4 are highest for $R_{\max}$ (1.75 mm and 2.33 mm) and lower in $R_{\text{avg}}$ (0.31 mm and 0.64 mm). For $R_{\min}$, the errors are 0.39 mm and 1.20 mm.

While the experts demonstrated millimeter accuracy in identifying Midpoints 1 and 2, a slight deviation was observed in the identification of Midpoints 3 and 4. Thus, Accurately identifying specific points within thin, translucent structures such as the falx cerebri proves challenging, even for experts.

Table 3 Comparison of Errors in Midpoint locations. This table illustrates the errors (in mm) in Spatial locations of four midpoints. comparing results between experts and baseline

| | Midpoint 1 | | | Midpoint 2 | | | Midpoint 3 | | | Midpoint 4 | | |
|---|---|---|---|---|---|---|---|---|---|---|---|---|
| | Rmin | Ravg | Rmax | Rmin | Ravg | Rmax | Rmin | Ravg | Rmax | Rmin | Ravg | Rmax |
| Expert 1 | 0.34 | 0.53 | 0.82 | 0.93 | 0.26 | 0.51 | 1.05 | 0.79 | 2.06 | 0.39 | 0.31 | 2.33 |
| Expert 2 | 1.00 | 0.67 | 0.35 | 0.60 | 0.35 | 0.74 | 1.55 | 1.05 | 1.56 | 1.20 | 0.64 | 1.75 |
| **mean** | **0.67** | **0.60** | **0.58** | **0.76** | **0.30** | **0.62** | **1.30** | **0.92** | **1.81** | **0.80** | **0.48** | **2.04** |

#### 3.3. Non-expert performance with the guideline

Three rotated images ($R_{min}$, $R_{avg}$, $R_{max}$) were rearranged in four unique orders using permutation, resulting in four subgroups, each containing three images. four non-experts were tasked with realigning these rotated images to the AC-PC plane using the proposed guideline. The non-experts received both PDF and video versions of the guideline at least two hours before the experiment and were instructed to read the guideline and watch the video instructions. The experiment commenced once the non-experts were ready to perform the alignment. They were allowed to review the instructional video or the PDF guideline during the experiment. To evaluate the performance of the non-experts, we compared the spatial locations of the borders of the AC and the PC, And four midline points, as well as the rotation angles and RMSD, to the baseline.

##### 3.3.1. AC and PC location errors

In the study of the spatial locations of the AC and the PC, non-experts demonstrated proficient performance under minimal rotation $R_{min}$. The maximum errors recorded were 0.80 mm for AC, 0.50 mm for PC, and 0.75 mm for the AC-PC distance, all within a one voxel (0.8 mm) difference. However, under average rotation, $R_{avg}$, the peak value increased to 1.09 mm for AC, 1.15 mm for PC, and 1.33 mm for the AC-PC distance. With further increased rotation as in $R_{max,}$ the maximum errors observed were 2.16 mm for AC, 1.20 mm for PC, and 1.63 mm for the AC-PC distance. Despite a few minor deviations, the use of guidelines enabled non-experts to achieve accuracy nearly on par with that of the experts.

Table 4. Comparison of Errors in AC-PC Location and AC-PC Distance. This table illustrates the errors (in mm) in AC-PC location and AC-PC distance, comparing results between Non-experts and baseline.

| | Rmin | | | Ravg | | | Rmax | | |
|---|---|---|---|---|---|---|---|---|---|
| | ac | pc | ac-pc distance | ac | pc | ac-pc distance | ac | pc | ac-pc distance |
| Non-Expert 1 | 0.56 | 0.50 | 0.75 | 1.09 | 0.34 | 0.25 | 0.52 | 0.75 | 0.62 |
| Non-Expert 2 | 0.80 | 0.28 | 0.15 | 0.76 | 0.67 | 0.44 | 2.16 | 0.28 | 1.63 |
| Non-Expert 3 | 0.34 | 0.46 | 0.44 | 0.68 | 1.15 | 1.33 | 0.51 | 0.34 | 0.41 |
| Non-Expert 4 | 0.33 | 0.29 | 0.07 | 0.46 | 0.64 | 0.01 | 0.15 | 1.20 | 0.61 |
| **mean** | **0.51** | **0.38** | **0.35** | **0.75** | **0.70** | **0.51** | **0.84** | **0.64** | **0.82** |

##### 3.3.2. Midpoint location errors

The non-experts demonstrated superior performance in identifying Midpoints 1 and 2 under rotation conditions of $R_{min}$, $R_{avg}$, with most errors being under 1 mm. However, two outliers were noted in $R_{avg}$, where errors of 1.84 mm and 1.97 mm were recorded for Midpoints 1 and 2, respectively. In $R_{max}$, the errors for Midpoint 1 ranged between 0.26 mm and 1.32 mm, while for Midpoint 2, the errors varied from 0.71 mm to 1.32 mm. The error range for Midpoints 3 and 4 showed a notable increase, particularly under more challenging rotation conditions. For Midpoint 3, under the $R_{min}$, condition, errors ranged from 0.37 mm to 2.87 mm. In the $R_{avg}$, condition, errors spanned from 0.91 mm to 2.69 mm, excluding an outlier of 28.68 mm from the mean calculation. Under $R_{max}$, the errors varied significantly, with one instance showing a deviation as large as 7.12 mm. Similarly, for Midpoint 4, under the $R_{min}$, condition, errors ranged from 0.92 mm to 2.17 mm. In the $R_{avg}$, condition, errors fell between 0.76 mm and 3.36 mm, again excluding an outlier of 23.63 mm from the mean. Under $R_{max}$, errors ranged from 2.87 mm to 6.10 mm.

In summary, while non-experts demonstrated near expert-level accuracy in identifying Midpoints 1 and 2, they showed significant deviations for Midpoints 3 and 4, especially under increased rotation conditions such as $R_{avg}$, $R_{max}$. This highlights the challenges in accurately identifying specific points within the falx cerebri, even with the aid of guidelines.

Table 5. Comparison of Errors in Midpoint Locations (outliers excluded). This table illustrates the errors (in mm) in Spatial locations of four midpoints. comparing results between Non-experts and baseline. Mean values exclude identified outliers

| | Midpoint 1 | | | Midpoint 2 | | | Midpoint 3 | | | Midpoint 4 | | |
|---|---|---|---|---|---|---|---|---|---|---|---|---|
| | Rmin | Ravg | Rmax | Rmin | Ravg | Rmax | Rmin | Ravg | Rmax | Rmin | Ravg | Rmax |
| Non-Expert 1 | 0.26 | 0.61 | 1.32 | 0.96 | 0.21 | 0.71 | 1.64 | 28.68 | 2.33 | 1.11 | 23.63 | 2.87 |
| Non-Expert 2 | 0.56 | 0.68 | 0.62 | 0.99 | 0.46 | 1.10 | 2.87 | 2.69 | 7.12 | 2.17 | 3.36 | 6.10 |
| Non-Expert 3 | 0.47 | 1.84 | 1.07 | 0.97 | 1.97 | 1.32 | 0.37 | 0.98 | 2.27 | 1.13 | 1.56 | 3.18 |
| Non-Expert 4 | 0.46 | 0.67 | 0.26 | 0.96 | 0.25 | 0.78 | 2.01 | 0.91 | 0.97 | 0.92 | 0.76 | 3.95 |
| **mean** | **0.44** | **0.95** | **0.82** | **0.97** | **0.72** | **0.98** | **1.72** | **1.53** | **3.17** | **1.33** | **1.89** | **4.02** |

#### 3.4. Comparisons against automated realignment

Three automated realignment approaches were compared against the proposed guideline (i.e. the baseline established in section 3.1.2): AC-PC realignment using “acpcdetect” (Ardekani & Bachman, 2009), template registration using FSL (Jenkinson et al., 2012), and realignment based on point-warping using ANTs (Avants et al., 2008) + Slicer 3D (Fedorov et al., 2012).

The fully automated AC-PC realignment (acpcdetect) tool integrated into the Automatic Registration Toolbox (ART) realigns the structural MR images to the Schaltenbrand version of the AC-PC plane (Ardekani & Bachman, 2009). It employs a three-stage approach to realign individual MR images to the AC-PC plane. In the first stage, the midsagittal plane is estimated, and the image is rotated to align with this plane (Ardekani et al., 1997) In the second stage, AC and PC are automatically identified using a model-based algorithm. A transformation is then computed to realign the images to the AC-PC plane (Ardekani & Bachman, 2009).The third stage focuses on stabilizing this standard orientation by automatically identifying eight additional landmarks. A subsequent transformation is calculated to align these landmarks as closely as possible with predetermined target locations (Ardekani & Alzheimer's Disease Neuroimaging, 2022). Throughout all three stages, a rigid body registration method is employed to achieve the alignment.

For the fully automated AC-PC alignment method, no specific data preparation was necessary other than ensuring that the input data was in a short or unsigned short NIfTI format (Ardekani & Bachman, 2009). As for the manual realignments, we assessed the performance of this fully automated method by analyzing errors in rotation angles and calculating root mean square deviation against the baseline. Although the tool outputs the spatial locations of the AC and the PC, their convention (Schaltenbrand-center of ac to center of pc) differs from the one we adopted for the baseline (Clinical-posterior border of ac to anterior border of pc). Therefore, a direct quantitative comparison was not possible. However, we qualitatively assessed the spatial locations of the AC and PC( figures 18,19,20).

The template registration method, which is applied in the pre-processing pipeline of the WU-Minn Human Connectome Project (HCP) (Glasser et al., 2013), involves a rigid-body registration to a standard template image using 6 degrees of freedom (DOF).The rigid-body transformation matrix is derived from an initial 12 DOF affine registration to the template image. This approach preserves the shape and size of the individual subject (Glasser et al., 2013). A significant drawback of this approach is that its effectiveness is highly dependent on the performance of the underlying registration algorithms (Dadar et al., 2018) and the quality of the template image used.

As template registration critically depends on the quality of the template, the evaluation of this method was conducted in two distinct stages. First, we followed the HCP preprocessing protocol, which involves rigidly aligning the native structural image to the MNI152-2007 T1 template (Glasser et al., 2013). In the second stage, we applied MNI AC-PC alignment, where the MNI152-2007 T1 template is realigned to the AC-PC plane according to our specified guidelines. The AC-PC aligned template was then utilized for rigid body registration, with realignment performed using FSL's FLIRT (Greve & Fischl, 2009; Jenkinson et al., 2002; Jenkinson & Smith, 2001) and Apply-Warp (Andersson JLR, 2010; Glasser et al., 2013; Jenkinson et al., 2002; Jenkinson et al., 2012). The primary outputs from this approach were the transformed image and the transformation matrix. The absence of spatial landmark points necessitated a focus on evaluating the rotation error and the root mean squared deviations

(RMSD) exclusively. Consequently, these two metrics were the primary indicators of the method's accuracy and performance.

The point-warping method offers a sophisticated approach to aligning MRI images to AC-PC space (Horn et al., 2017; Pallavaram et al., 2015).This process also unfolds in two primary stages. Initially, the spatial coordinates of the AC, PC, and midline points are identified within a template image These landmark points are then subjected to a nonlinear transformation, which warps them onto the native structural image (Horn et al., 2017). Using these warped landmark points, the desired AC-PC line and midplane are established within the native structural image. Subsequently, the transformation matrix is computed to precisely realign the image to the AC-PC space (Horn & Kuhn, 2015; Oxenford et al., 2022). This method offers the advantage of performing realignment on the native structural image based on subject-specific anatomy. Nonetheless, the efficacy of this realignment is significantly influenced by the nonlinear algorithm employed within the registration software (Klein et al., 2009; Liu & Dawant, 2014), as well as the various algorithms utilized in the subsequent processing stages.

When applying the point-warping method, defining key landmark points within the chosen T1 template space, specifically MNI 152 2009b (Fonov et al., 2009),is essential. Here, these landmarks were identified using two distinct approaches: (1) as proposed by Horn et al in LeadDBS (Horn & Kuhn, 2015; Horn et al., 2017) and (2) according to our protocol proposed here. Once identified, the landmark points were non-linearly warped to the individual brain using ANTs SyN registration and point registration techniques (Avants et al., 2008). Following this, the landmarks were used to define the AC-PC line and the midplane. These definitions were then utilized to create the transformation using AC-PC transformation module in Slicer 3D software (Fedorov et al., 2012).The primary outputs of point warping methods include the spatial coordinates of the AC, PC, and midline points. Additionally, these methods produce both the transformed image aligned to the AC-PC axis and the corresponding transformation matrix. In the points registration method proposed by (Horn et al., 2017). the AC and PC landmarks also serve as midline points, alongside an additional landmark in the falx cerebri. Consequently, we could only assess the spatial locations of the AC and PC, the AC-PC distance, rotation angles, and root mean square Deviation. conversely, the points warping method using the proposed guideline allows for a more comprehensive comparison. It enables the evaluation of the spatial locations of the AC, PC, and midpoints, as well as rotation and root mean square deviation.

The automated realignment approaches were implemented in different software packages (FSL and ART), whereas the guideline-based procedure was performed in 3D Slicer. Because each tool uses its own coordinate convention and outputs transformation matrices in different formats, additional processing was required to make the results comparable. Specifically, all transformation matrices produced by acpcdetect and FSL were converted to ITK format using the C3D affine tool within ITK (Yoo et al., 2002). No conversion was needed for 3D Slicer outputs, as they are already provided in ITK format. This harmonization step standardized the transformation matrices across methods. We then derived rotation-angle errors and computed the root mean squared deviation (RMSD) from the rotation matrices for all approaches relative to the baseline

We observed a sign inversion in the *Y* and *Z* rotation components when comparing rotation angles derived from the acpcdetect transformation matrix. To investigate this discrepancy, we applied the original acpcdetect matrix to the rotated image to test whether it reproduced the ACPC-aligned NIfTI output generated by acpcdetect. This comparison revealed a mismatch between the reported matrix and the exported image: applying the original matrix did not produce an ACPC-aligned image, whereas the NIfTI output produced directly by acpcdetect was correctly ACPC-aligned. Notably, manually flipping the signs of the *Y* and *Z* components of the original matrix yielded an image that was properly ACPC-aligned and consistent with the acpcdetect NIfTI output.

For transparency, we therefore report both the angular error and RMSD values computed using the original matrix and the *Y/Z* sign-corrected matrix. In the main manuscript, we report results corresponding to the matrix that reproducibly generates an ACPC-aligned image consistent with the acpcdetect output (i.e., the *Y/Z*-corrected matrix).

Table 6. RMSD values (mm) derived from two acpcdetect rotation matrices (the original output matrix and the Y–Z sign-flipped variant),quantifying the deviation of the acpcdetect transformation from the baseline.

| | *Rmin* | *Ravg* | *Rmax* | *Mean* |
|---|---|---|---|---|
| acpcdetect -original | 4.72 | 7.40 | 8.68 | 6.93 |
| acpcdetect- YZ flipped | 4.42 | 2.46 | 3.04 | 3.31 |

Table 7. Rotation angles (degrees) for all three axes extracted from two acpcetect matrix variants (the original output matrix and the Y–Z sign-flipped variant) and compared with the baseline. Absolute differences relative to the baseline are reported in brackets.

| | *Rmin* | | | *Ravg* | | | *Rmax* | | |
|---|---|---|---|---|---|---|---|---|---|
| | ***X*** | ***Y*** | ***Z*** | ***X*** | ***Y*** | ***Z*** | ***X*** | ***Y*** | ***Z*** |
| ***Baseline*** | ***-11.65*** | ***-0.35*** | ***0.09*** | ***8.54*** | ***3.48*** | ***2.26*** | ***27.27*** | ***-0.74*** | ***-4.58*** |
| acpcdetect -original | -15.96(4.31) | 2.02(2.37) | -0.65(0.74) | 6.04(2.50) | -3.57(7.05) | -1.02(3.29) | 24.26(3.01) | 1.97(2.71) | 4.14(8.72) |
| acpcdetect-YZ flipped | -15.96(4.31) | -2.02(1.66) | 0.65(0.56) | 6.04(2.50) | 3.57(0.10) | 1.02(1.24) | 24.26(3.01) | -1.97(1.23) | -4.14(0.44) |

##### 3.4.1. AC and PC location errors

The “acpcdetect” and template registration approaches did not allow for direct comparisons of the AC and PC location. The point-warping methods adopted from (Horn & Kuhn, 2015; Horn et al., 2017) and the one based on the proposed guideline exhibited comparable errors (with respect to the baseline defined in section 3.1.2), although the latter demonstrated slightly better performance. Specifically, the mean errors for the AC, PC, and AC-PC distances using the Horn et al. 2017 method were 1.17, 0.69 and 1.38 mm, respectively, while for guideline-based point-warping, these errors were 1.06, 0.56, and 0.82 mm. Despite these minor improvements, the differences are very small and negligible.

This outcome underscores the strong alignment between the conventions for defining AC-PC locations(posterior border of the AC to the anterior border of the PC) in the proposed guidelines and those employed in the method outlined by Horn et al (Horn & Kuhn, 2015; Horn et al., 2017).The similarities indicate consistency in approach, validating the reliability of the proposed guideline.

Table 8. Comparison of Errors in AC-PC Location and AC-PC Distance. This table illustrates the errors (mm) in AC-PC location and AC-PC distance, comparing results between nonlinear points warping method and baseline. The comparison is made both with Horn et al. method and our proposed guidelines method

| | Points warping method: Horn et al. | | | Points warping method: proposed guideline | | |
|---|---|---|---|---|---|---|
| | *ac* | *pc* | *ac-pc distance* | *ac* | *pc* | *ac-pc distance* |
| Rmin | 1.52 | 1.06 | 1.86 | 1.22 | 0.74 | 1.25 |
| Ravg | 0.91 | 0.22 | 0.78 | 0.91 | 0.70 | 0.21 |
| Rmax | 1.09 | 0.79 | 1.50 | 1.04 | 0.25 | 0.99 |
| **mean** | **1.17** | **0.69** | **1.38** | **1.06** | **0.56** | **0.82** |

##### 3.4.2. Midpoint location errors

In the point-warping method proposed by Horn et al., the authors selected the AC, PC, and an arbitrary point in the falx cerebri as midline markers to define the midplane. This approach differs from the landmarks we have chosen, making a direct comparison of midplane locations between the two methods not feasible. Therefore, we limited our comparison to the midpoint locations obtained from the guideline-based point-warping method and the baseline.

This approach demonstrated excellent correspondence for midpoints 1 and 2, with mean errors of less than 1 mm. However, midpoints 3 and 4 showed less accurate correspondence, with mean values of 16.64 mm and 16.97 mm, respectively. Although this deviation results in a larger error compared to the baseline, midpoints 3 and 4 are still accurately warped within the falx cerebri, ensuring precise selection of the midplane. Consequently, there were no adverse effects on the final realignment. In conclusion, while the point-warping method accurately warps points within the falx cerebri, it fails to preserve correspondence consistently.

Table 9. Comparison of Errors in Midpoint locations. This table illustrates the errors (mm) in four Midpoint locations, comparing results between nonlinear points warping method and baseline

| | Midpoint1 | Midpoint2 | Midpoint3 | Midpoint4 |
|---|---|---|---|---|
| Rmin | 0.99 | 1.06 | 15.34 | 26.17 |
| Ravg | 0.90 | 1.00 | 5.48 | 2.04 |
| Rmax | 0.28 | 0.57 | 29.11 | 22.69 |
| **mean** | **0.72** | **0.88** | **16.64** | **16.97** |

##### 3.4.4 .General Discussion on the Performance of Automated Methods

A comparison of the three automated realignment methods against the validated baseline realignment revealed that non-linear warping of key landmark points to subject-specific images using the ANTs SyN registration (Avants et al., 2008), followed by image realignment using the transformation derived from the warped points in 3D Slicer, yielded the most accurate alignment outperforming both fully automated (auto acpcdetect) and template-based registration methods. The effectiveness of this method is attributed to the nonlinear point warping technique's superior ability to accurately identify key landmarks such as the AC and PC with high precision. This level of accuracy is consistent with findings from previous work by (Liu & Dawant, 2014). Additionally, our results further demonstrate the method's accuracy in identifying other critical midline points 1 and 2. Although the spatial location of Midline points 3 and 4 were deviated they were accurately warped within the falx cerebri, resulting in no significant impact on midplane definition. These deviations likely reflect local deformation effects inherent to non-linear registration. The guideline-based points warping method demonstrated slightly better performance compared to the method proposed by (Horn et al., 2017) . However, the differences between the two are minimal and can be considered negligible.

In contrast, the suboptimal performance of the template registration method that uses the MNI152 template as a reference image for rigid body registration, is notable. The extreme errors observed with this method can be attributed to differences in how the AC-PC space is defined compared to the Talairach and Schaltenbrand conventions. Unsurprisingly, these errors are significantly reduced when the template is realigned to the clinically utilized version of the Schaltenbrand AC-PC plane. emphasizing the necessity for careful template selection based on specific application requirements.

To the best of our knowledge, the auto acpcdetect is the only open-source software that provides fully automated transformation of structural MR images to the Schaltenbrand version of the AC-PC plane. This tool is especially valuable for researchers working in deep brain stimulation (DBS) or anyone needing to realign multiple datasets to this specific version of the AC-PC plane. Our validation of the auto acpcdetect technique reveals significant deviations between the autodetect method and the baseline, indicating the algorithm's suboptimal performance (fig 18,19,20).

Closer analysis reveals that this model-based approach consistently fails to identify the PC in all three cases by a substantial margin. While the location of the AC is not perfect, it remains relatively close to the actual position. However, this discrepancy ultimately leads to an improper definition of the AC-PC line, which compromises the overall accuracy of the alignment.

Figure 18: Visual Comparison of the Automated Algorithm Performance Versus the Baseline Developed Using the Proposed Manual Guideline Under the Maximum Rotation Condition (Rmax). The left panels display sagittal sections highlighting the region of interest (ROI), while the right panels present enlarged views of the ROI for detailed comparison. As illustrated in panels A to C: A. Center of the anterior commissure (AC) identified by the automated AC-PC detection algorithm (blue dot) versus the actual center of AC (yellow) in the same slice. B. Center of the posterior commissure (PC) identified by the automated algorithm (blue dot) versus the actual center of PC (yellow). C. Schaltenbrand-version AC-PC line derived from the automatically detected AC and PC (blue) compared to the AC-PC line (reddish orange) derived from true centers of AC and PC (yellow)

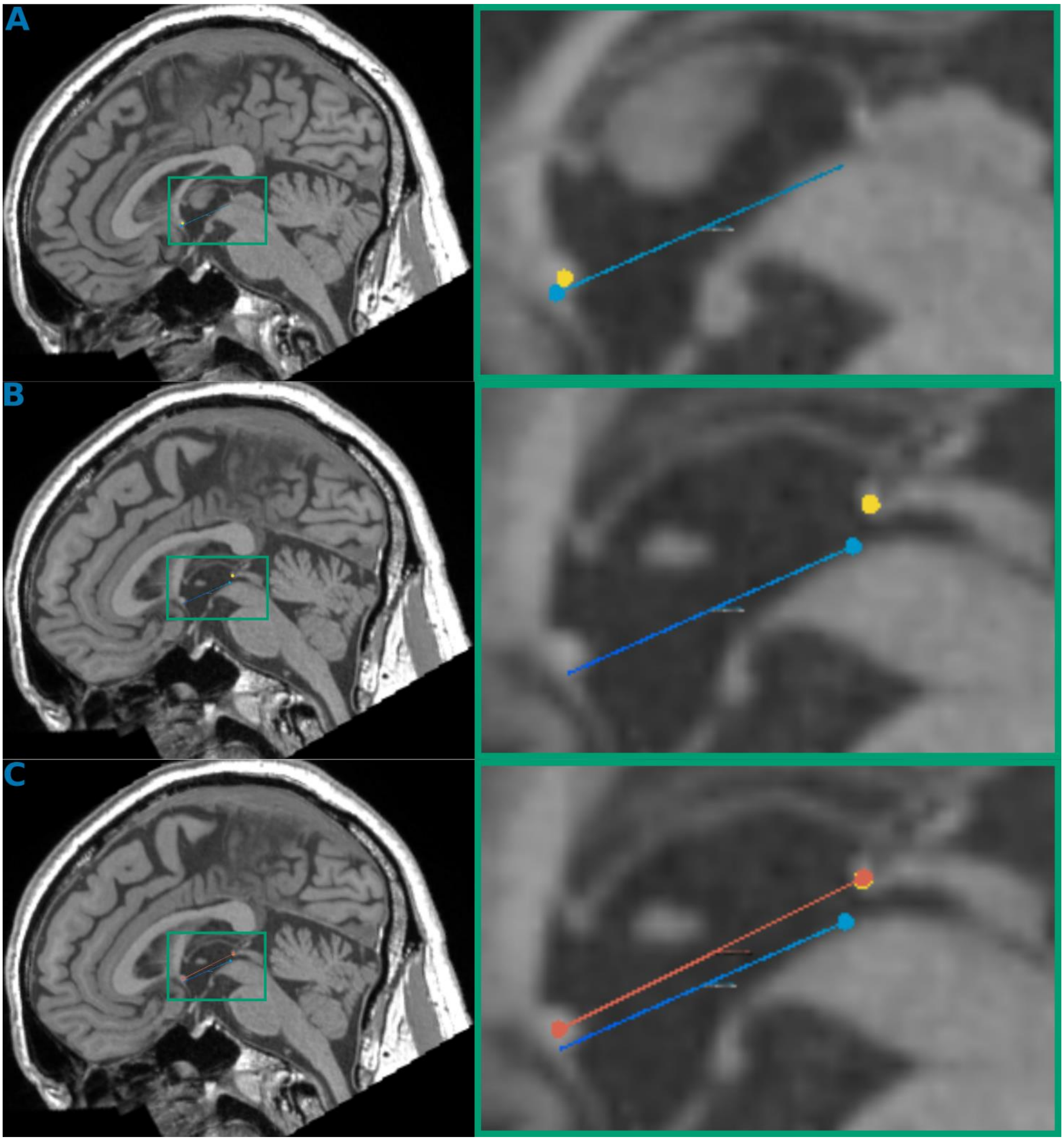

Figure 19: Visual Comparison of the Automated Algorithm Performance Versus the Baseline Developed Using the Proposed Manual Guideline Under the Minumum Rotation Condition (Rmin). The left panels display sagittal sections highlighting the region of interest (ROI), while the right panels present enlarged views of the ROI for detailed comparison. As illustrated in panels A to C: A. Center of the anterior commissure (AC) identified by the automated AC-PC detection algorithm (blue dot) versus the actual center of AC (yellow) in the same slice. B. Center of the posterior commissure (PC) identified by the automated algorithm (blue dot) versus the actual center of PC (yellow). C. Schaltenbrand-version AC-PC line derived from the automatically detected AC and PC (blue) compared to the AC-PC line (reddish orange) derived from true centers of AC and PC (yellow)

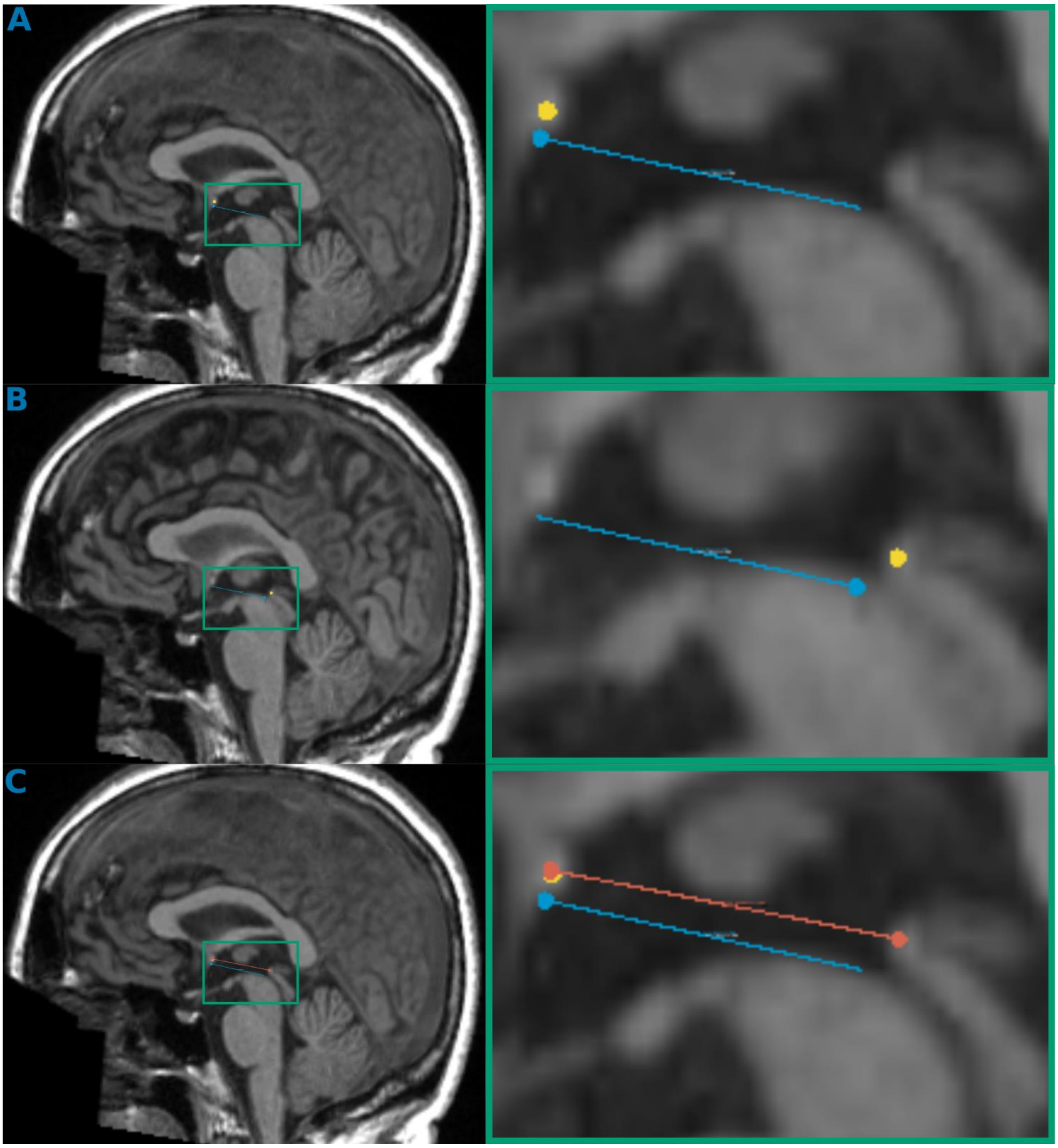

Figure 20: Visual Comparison of the Automated Algorithm Performance Versus the Baseline Developed Using the Proposed Manual Guideline Under the Average Rotation Condition (Ravg). The left panels display sagittal sections highlighting the region of interest (ROI), while the right panels present enlarged views of the ROI for detailed comparison. As illustrated in panels A to C: A. Center of the anterior commissure (AC) identified by the automated AC-PC detection algorithm (blue dot) versus the actual center of AC (yellow) in the same slice. B. Center of the posterior commissure (PC) identified by the automated algorithm (blue dot) versus the actual center of PC (yellow). C. Schaltenbrand-version AC-PC line derived from the automatically detected AC and PC (blue), only visible in panels A and B. compared to the AC-PC line (reddish orange) derived from true centers of AC and PC (yellow)

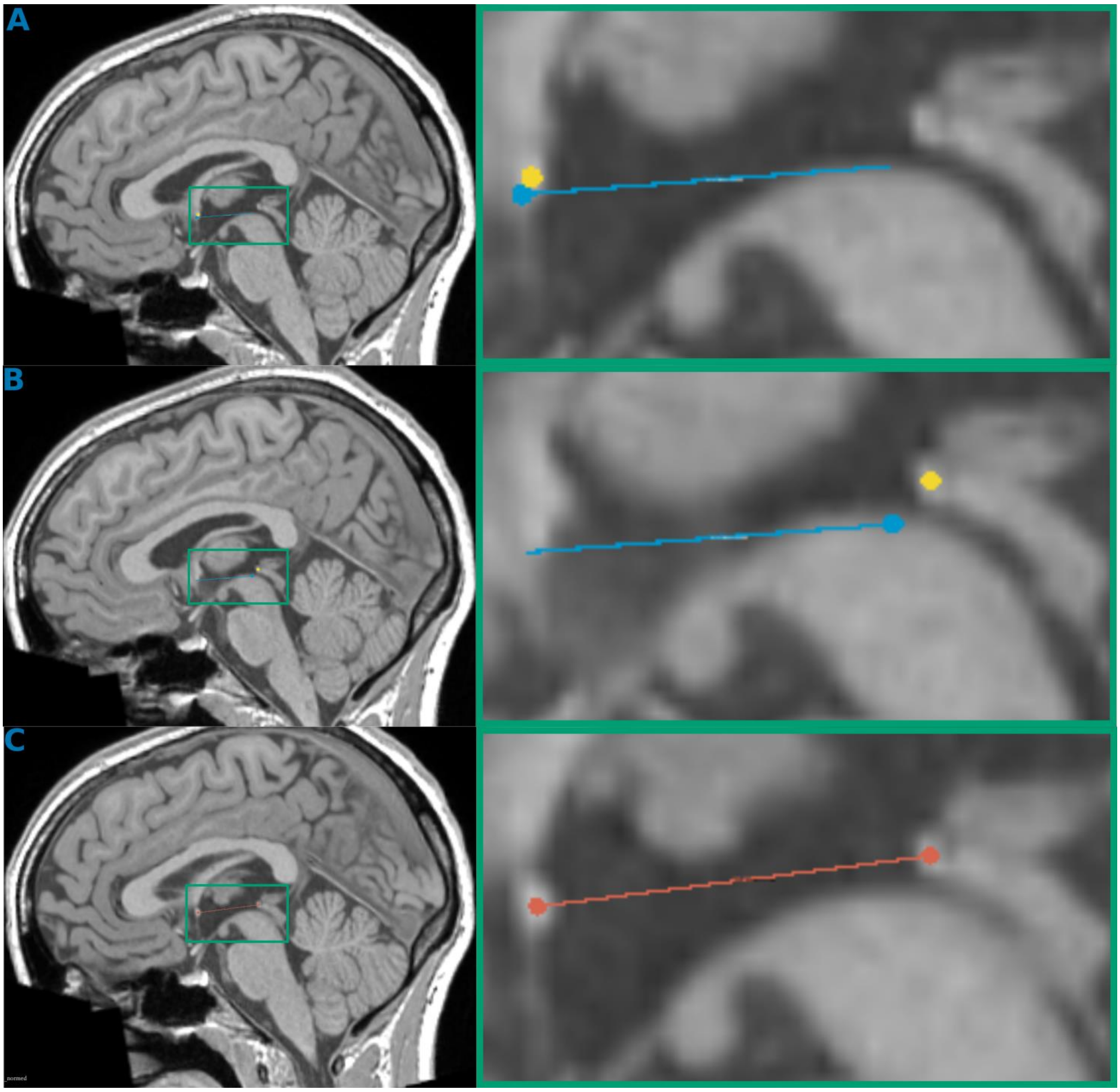

#### 4. Guideline Amendments

Based on a detailed analysis of the experimental results and feedback from both expert and non-expert observers, three amendments were made to the precursor guideline. Below are the justifications for each change and the corresponding modifications implemented in the guidelines:

##### Amendment 1 – Final guideline step 2.2.3: AC-PC line

While the results from manual observers were generally accurate, small deviations did occur. Due to the close proximity of the AC and PC structures, even minor discrepancies can lead to significant rotational errors in the final realignment. [Figure 21 -24]. Therefore, accurately defining the AC-PC line is critical. To address this issue, we have introduced an additional step into the final guideline specifically, Step 2.3.3. This step emphasizes the importance of precise AC-PC line identification, supported by visual aids to enhance accuracy and consistency.

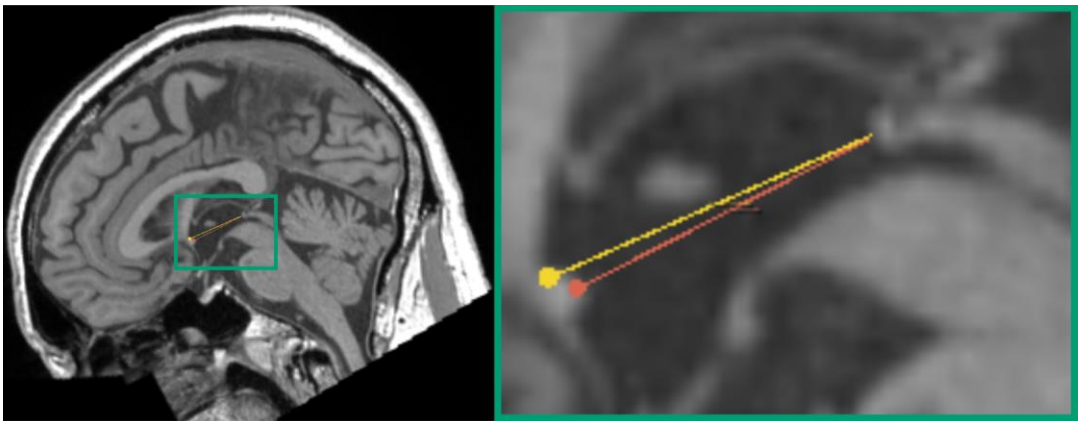

Figure 21: Comparison of AC-PC Line Definitions. The left panel presents a midsagittal section demonstrating the variation between the two AC-PC lines within the region of interest. The right panel provides a magnified view highlighting the differences between the baseline AC-PC line (reddish-orange) and the AC-PC line determined by Nonexpert-2 (yellow). Notice the discrepancy in identifying the AC , which consequently leads to a larger angular difference.

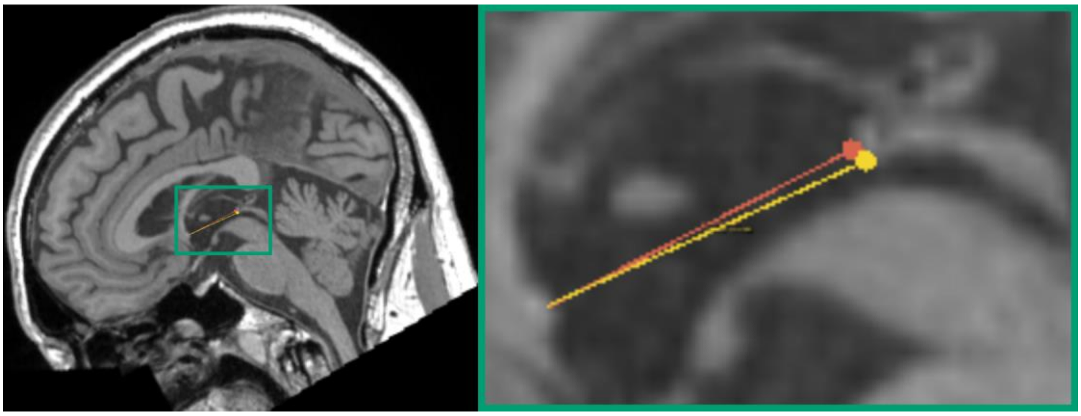

Figure 22: Comparison of AC-PC Line Definitions. The left panel presents a midsagittal section demonstrating the variation between the two AC-PC lines within the region of interest. The right panel provides a magnified view highlighting the differences between the baseline AC-PC

line (reddish-orange) and the AC-PC line determined by Nonexpert-4 (yellow). Notice the discrepancy in identifying the PC , which consequently leads to a larger angular difference.

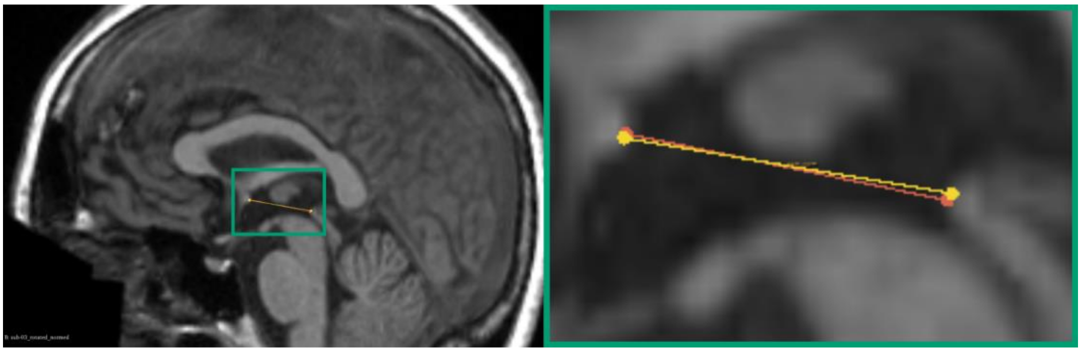

Figure 23: Comparison of AC-PC Line Definitions. The left panel presents a midsagittal section demonstrating the variation between the two AC-PC lines within the region of interest. The right panel provides a magnified view highlighting the differences between the baseline AC-PC line (reddish-orange) and the AC-PC line determined by expert-1 (yellow)

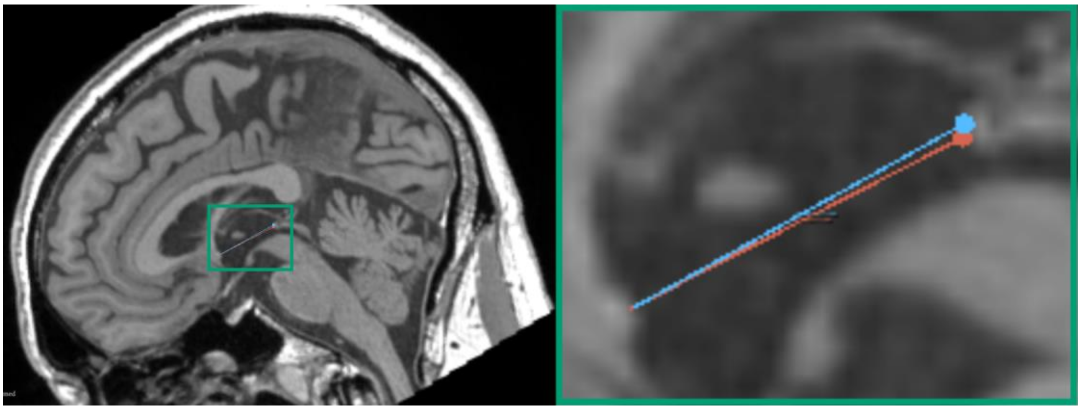

Figure 24: Comparison of AC-PC Line Definitions. The left panel presents a midsagittal section demonstrating the variation between the two AC-PC lines within the region of interest. The right panel provides a magnified view highlighting the differences between the baseline AC-PC line (reddish-orange) and the AC-PC line determined by expert-1 (yellow)

##### Amendment 2 -final guideline Step 2.6: Identifying midpoints 3 and 4

Although all manual observers correctly identified midpoint locations 3 and 4, including Non-Expert 1, who caused notable deviations of 28.68 mm and 23.63 mm by skipping a step in the precursor guideline (Specifically step 1.5 (2)), they all encountered difficulties in maintaining correspondence. For instance, they struggled to place a landmark point precisely at the junction between the falx cerebri and the superior sagittal sinus in the axial plane at the highest level of the corpus callosum. Although this minor degree of inaccuracy is expected due to the inherent challenges of tracing delicate, sheet-like structures in the brain using MR images (Liu &

Dawant, 2015), we aim to provide additional clarification to improve the identification of midpoints 3 and 4 in the precursor guideline. To address this, we revised the guideline to include clearer instructions for identifying these two midpoints within the falx cerebri. By utilizing visual aids, we have broken down the instructions into three sub-steps (2.6.1-2.6.3 in final guideline), enhancing clarity and improving the accuracy of landmark identification.

##### Amendment 3 - final guideline 2.3: all three verification criteria

Another significant refinement was made to the verification criteria of the precursor guideline. In the precursor guideline, we initially adopted verification criteria from Ramirez et al. (Ramirez et al., 2014) to validate AC-PC realignment. While these criteria were effective at identifying large deviations, our results and feedback from manual observers indicated that they may lack sensitivity to smaller misalignments, particularly around the X-axis, which is crucial for accurate AC-PC alignment.

To improve sensitivity to smaller errors, we revised the three verification criteria for more precise AC-PC alignment detection. In the precursor guideline, verification of pitch or rotation around the X-axis involved visually confirming that the AC and PC were positioned at the same level on both sagittal and axial planes. In the sagittal plane, this was confirmed by drawing a straight line connecting the centres of AC to the PC, while in the axial plane, alignment was verified by ensuring the AC and PC appeared on the same slice, creating the characteristic "keyhole" appearance of the third ventricle (Ramirez et al., 2014). However, given that the AC is larger than the PC approximately 3 mm versus 2 mm in sagittal diameter each structure can span more than one slice in images with a 1 mm isotropic voxel resolution, As a result, evaluating a single slice may overlook subtle misalignments(Figure26). Therefore, we now recommend examining three consecutive slices in the axial plane to confirm precise AC-PC realignment and assessing the full diameter of both the AC and PC in the sagittal plane. This optimized approach improves detection of finer misalignments and supports more accurate realignment verification.

In the precursor guideline, roll verification depended on visually assessing eyeball levels (Ramirez et al., 2014), where visual aids illustrated balanced eyeballs as a cue for accurate AC-PC realignment and unbalanced eyeballs as an indicator of misalignment. While this visual cue is effective for detecting large errors, it may miss smaller, subtler misalignments that are more difficult to detect. To address this, we optimized the visual aids to illustrate changes associated with smaller errors, making the verification process more sensitive to minor discrepancies. Additionally, we incorporated the coronal plane in the evaluation (Hamilton et al., 2017), providing a more comprehensive perspective to detect and correct subtle alignment errors. This enhanced approach allows for finer accuracy in verifying roll alignment. We adopted a similar approach for verifying yaw in the precursor guideline, the axial view example showing a large Z-axis rotational deviation has been replaced with an example image highlighting smaller deviations, enabling the detection of more subtle misalignments (Hamilton et al., 2017; Ramirez et al., 2014)

#### 5.Examples of Minor rotational deviations

The verification criteria outlined in our initial guideline assist in visually validating the results of manual AC-PC realignment. However, the examples we provided are particularly sensitive to larger errors, potentially limiting their effectiveness in detecting subtle misalignments. To visually illustrate examples of smaller deviations around the X, Y, and Z axes, we first realigned the MNI152 T1-weighted image, with a 1mm isotropic voxel resolution, to the AC-PC plane following the precursor guideline. Using this AC-PC alignment as the baseline, we then applied incremental rotations to the image from -3 to +3 degrees (i.e., -3, -2, -1, 1, 2, 3) along each of the three axes.

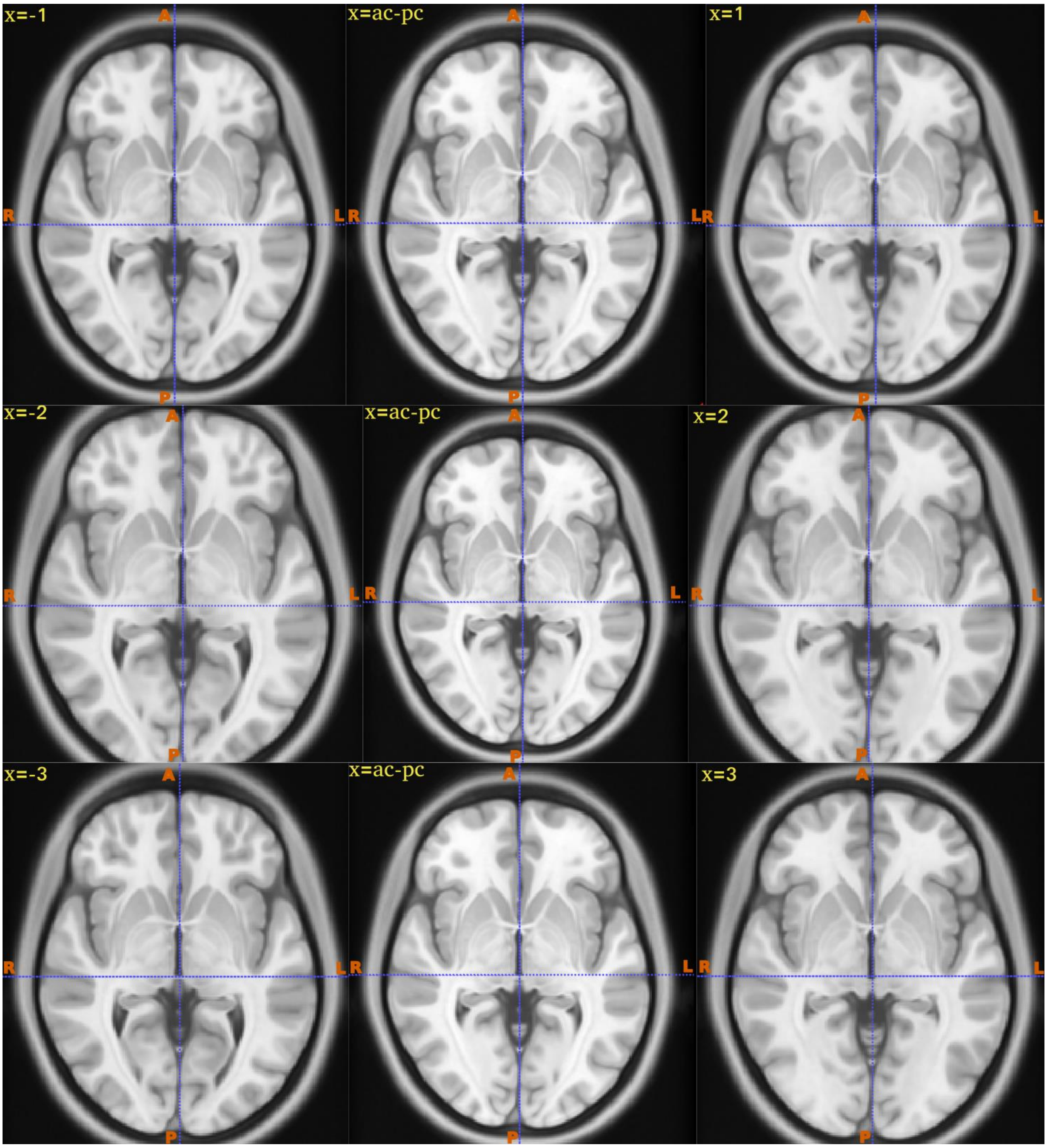


Figure 25: Axial plane images demonstrating rotational deviations around the X-axis relative to the correctly aligned AC-PC reference image. The left panels illustrates images with negative rotational deviations (-1°, -2°, -3°), the middle panels shows the precisely realigned AC-PC plane, and the right panels presents images with positive rotational deviations (+1°, +2°, +3°). Notably, despite these deviations, the AC-PC landmarks consistently appear in the same axial plane across all images. This consistency suggests that relying solely on the visual confirmation of AC and PC appearing in a single axial slice may not effectively detect subtle misalignments.

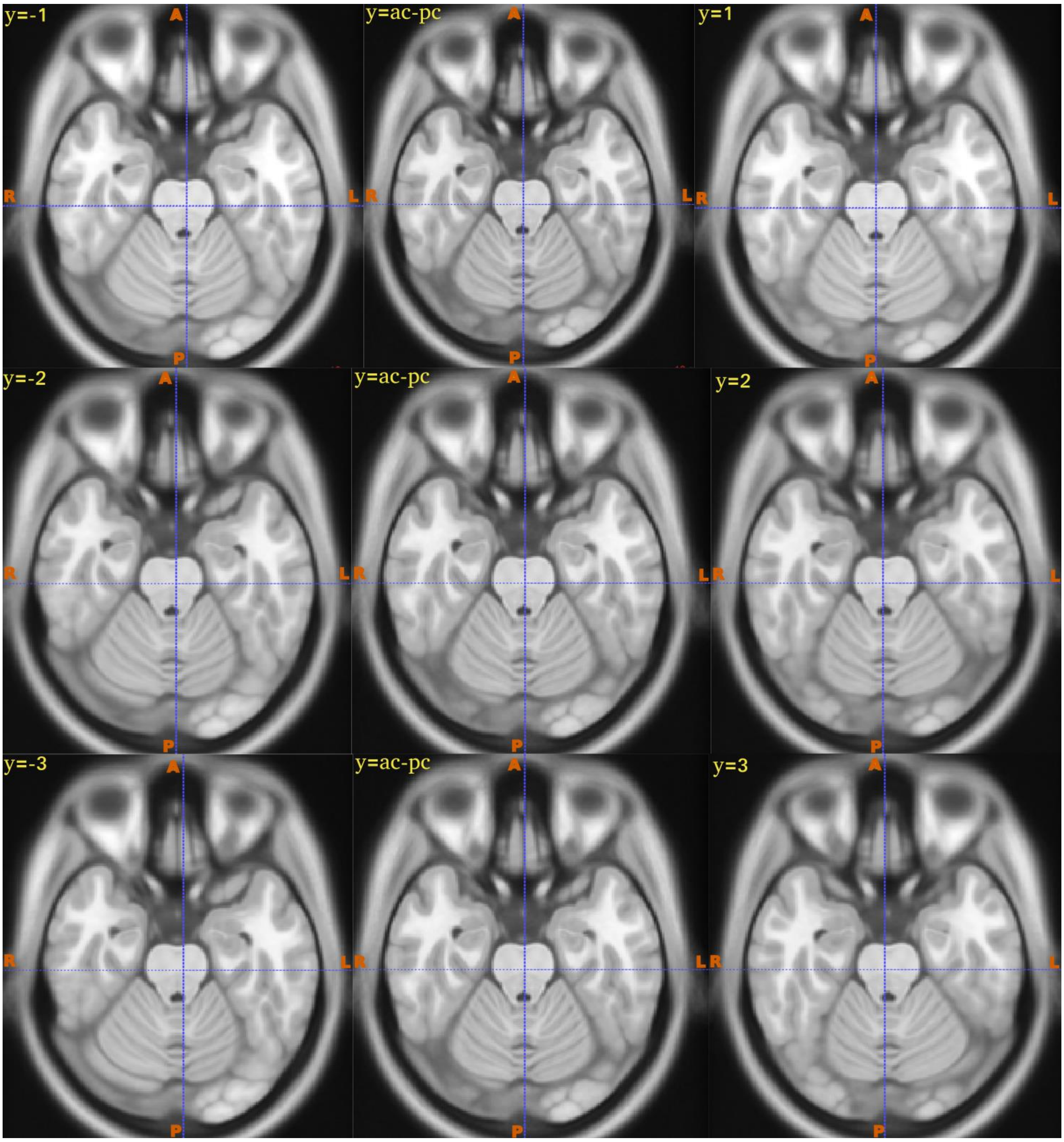


Figure 26: Axial planes illustrating rotational deviations around the Y-axis relative to the AC-PC realigned image. The left panels demonstrates negative rotational deviations (-1°, -2°, -3°), highlighting the visual effect where the right eyeball appears larger than the left. The middle panels represents the correctly realigned AC-PC plane, characterized by symmetry and equal appearance of both eyeballs. Conversely, the right panels illustrates positive rotational deviations (+1°, +2°, +3°), resulting in the left eyeball appearing larger than the right.

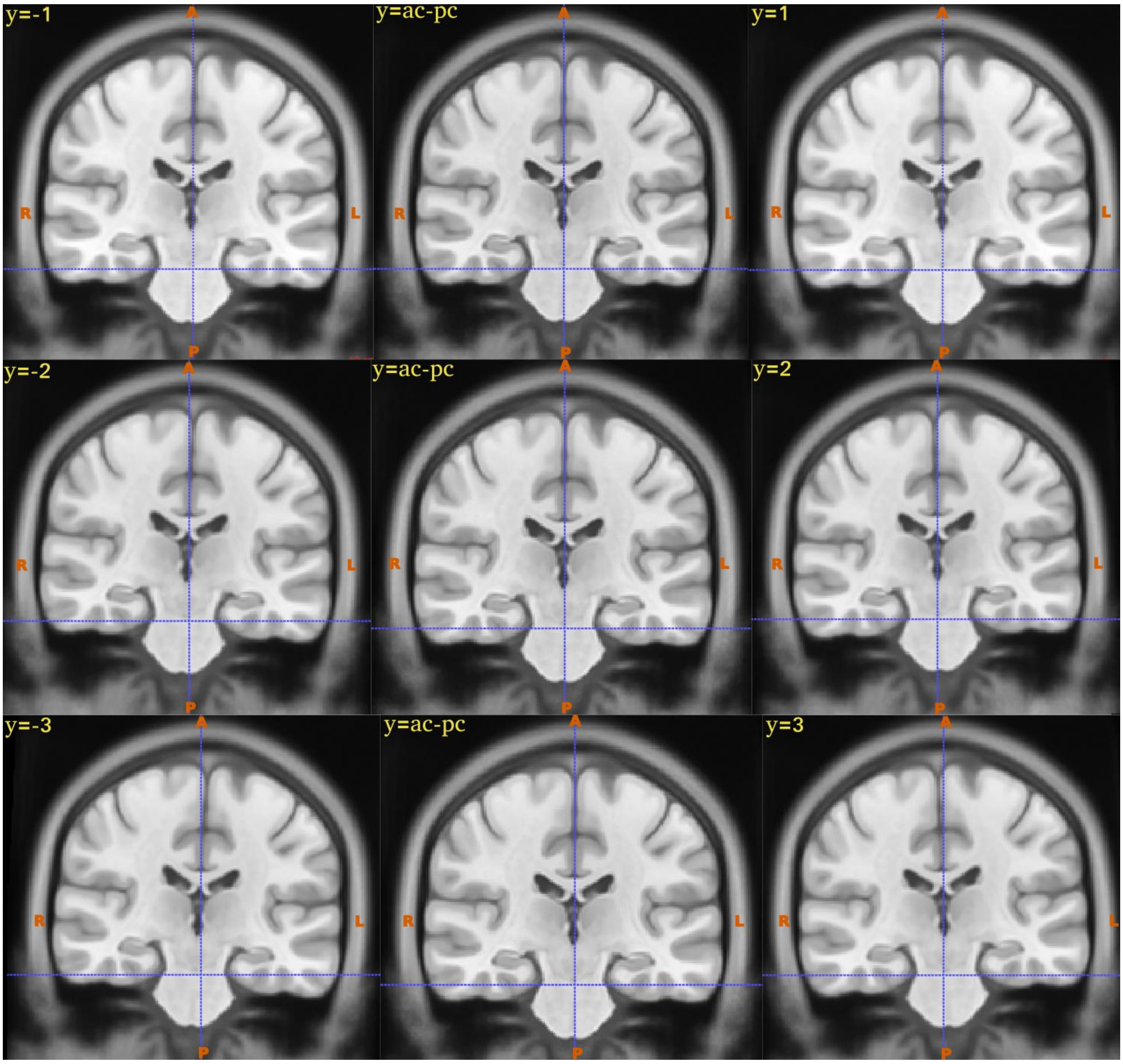


Figure 27 : Coronal planes illustrating rotational deviations around the Y-axis relative to the AC-PC realigned reference image. The left panel demonstrates negative rotational deviations (-1°, -2°, -3°), with the vertical crosshair shifted to the right of midline. The central panel depicts the accurately realigned AC-PC plane, with the vertical crosshair precisely positioned at the midline. The right panel illustrates positive rotational deviations (+1°, +2°, +3°), where the vertical crosshair shifts to the left of midline.

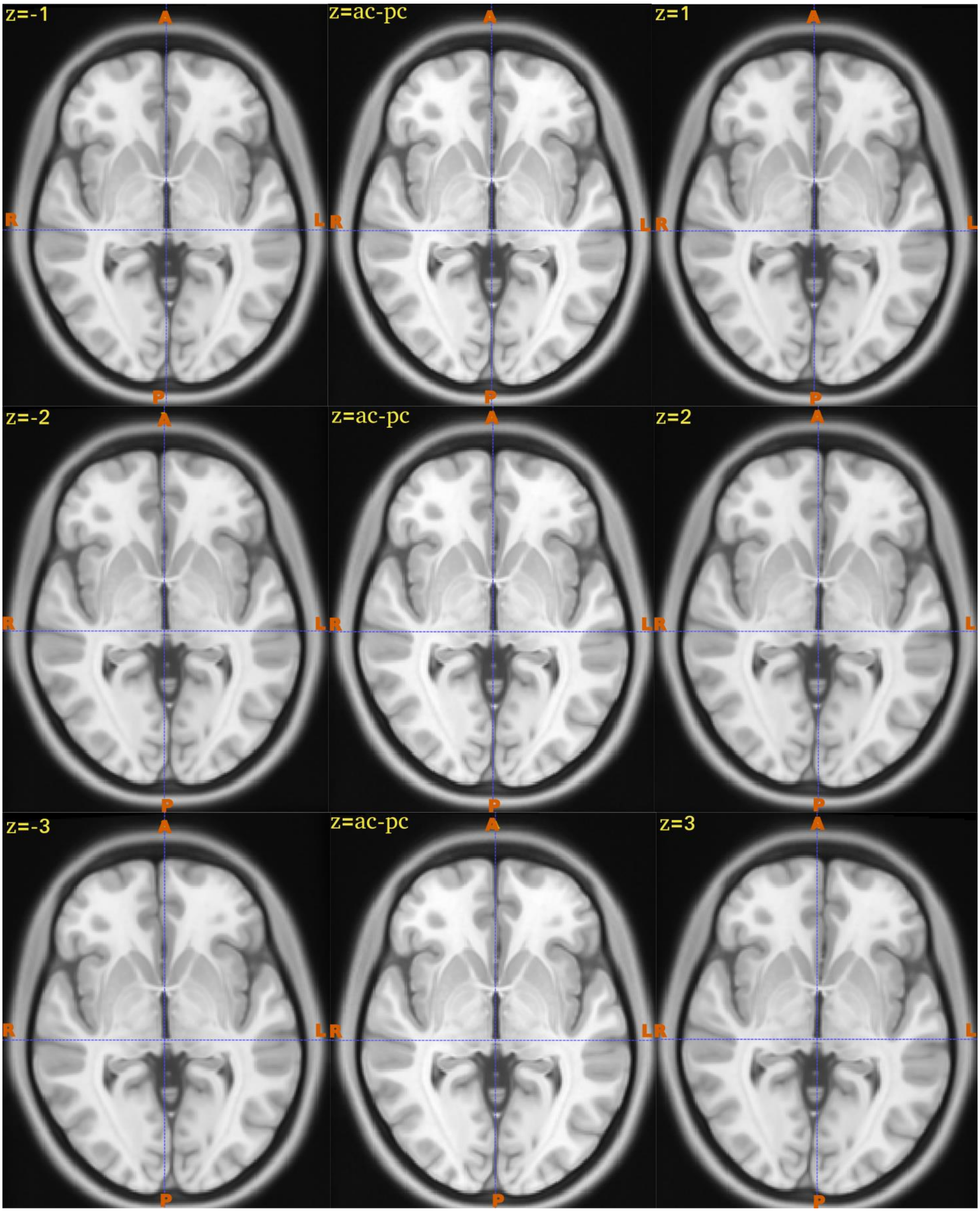


Figure 28: Axial planes illustrating rotational deviations around the Z-axis relative to the accurately aligned AC-PC plane. The left panels presents images demonstrating negative rotational deviations (-1°, -2°, -3°), characterized by the vertical crosshair shifting progressively to the left. The middle panels depicts the precisely realigned AC-PC plane, where the vertical crosshair is centered accurately along the midline. The right panels exhibits images illustrating positive rotational deviations (+1°, +2°, +3°), with the vertical crosshair progressively shifting to the right.

### 4.References


Andersson JLR, J. M., Smith S (2010). *Non-linear registration, aka spatial normalisation. FMRIB technical report TR07JA2*.

Ardekani, B. A., & Alzheimer's Disease Neuroimaging, I. (2022). A new approach to symmetric registration of longitudinal structural MRI of the human brain. *J Neurosci Methods*, *373*, 109563. https://doi.org/10.1016/j.jneumeth.2022.109563

Ardekani, B. A., & Bachman, A. H. (2009). Model-based automatic detection of the anterior and posterior commissures on MRI scans. *Neuroimage*, *46*(3), 677-682. https://doi.org/10.1016/j.neuroimage.2009.02.030

Ardekani, B. A., Kershaw, J., Braun, M., & Kanno, I. (1997). Automatic detection of the mid-sagittal plane in 3-D brain images. *IEEE Trans Med Imaging*, *16*(6), 947-952. https://doi.org/10.1109/42.650892

Avants, B. B., Epstein, C. L., Grossman, M., & Gee, J. C. (2008). Symmetric diffeomorphic image registration with cross-correlation: evaluating automated labeling of elderly and neurodegenerative brain. *Med Image Anal*, *12*(1), 26-41. https://doi.org/10.1016/j.media.2007.06.004

Dadar, M., Fonov, V. S., Collins, D. L., & Alzheimer's Disease Neuroimaging, I. (2018). A comparison of publicly available linear MRI stereotaxic registration techniques. *Neuroimage*, *174*, 191-200. https://doi.org/10.1016/j.neuroimage.2018.03.025

Fedorov, A., Beichel, R., Kalpathy-Cramer, J., Finet, J., Fillion-Robin, J. C., Pujol, S., Bauer, C., Jennings, D., Fennessy, F., Sonka, M., Buatti, J., Aylward, S., Miller, J. V., Pieper, S., & Kikinis, R. (2012). 3D Slicer as an image computing platform for the Quantitative Imaging Network. *Magn Reson Imaging*, *30*(9), 1323-1341. https://doi.org/10.1016/j.mri.2012.05.001

Fonov, V. S., Evans, A. C., McKinstry, R. C., Almli, C. R., & Collins, D. L. (2009). Unbiased nonlinear average age-appropriate brain templates from birth to adulthood. *Neuroimage*, *47*, S102. https://doi.org/https://doi.org/10.1016/S1053-8119(09)70884-5

Glasser, M. F., Sotiropoulos, S. N., Wilson, J. A., Coalson, T. S., Fischl, B., Andersson, J. L., Xu, J., Jbabdi, S., Webster, M., Polimeni, J. R., Van Essen, D. C., Jenkinson, M., & Consortium, W. U.-M. H. (2013). The minimal preprocessing pipelines for the Human Connectome Project. *Neuroimage*, *80*, 105-124. https://doi.org/10.1016/j.neuroimage.2013.04.127

Greve, D. N., & Fischl, B. (2009). Accurate and robust brain image alignment using boundary-based registration. *Neuroimage*, *48*(1), 63-72. https://doi.org/10.1016/j.neuroimage.2009.06.060

Hamilton, L. S., Chang, D. L., Lee, M. B., & Chang, E. F. (2017). Semi-automated Anatomical Labeling and Inter-subject Warping of High-Density Intracranial Recording Electrodes in Electrocorticography. *Front Neuroinform*, *11*, 62. https://doi.org/10.3389/fninf.2017.00062

Horn, A., & Kuhn, A. A. (2015). Lead-DBS: a toolbox for deep brain stimulation electrode localizations and visualizations. *Neuroimage*, *107*, 127-135. https://doi.org/10.1016/j.neuroimage.2014.12.002

Horn, A., Kuhn, A. A., Merkl, A., Shih, L., Alterman, R., & Fox, M. (2017). Probabilistic conversion of neurosurgical DBS electrode coordinates into MNI space. *Neuroimage*, *150*, 395-404. https://doi.org/10.1016/j.neuroimage.2017.02.004

Jansen, M. G., Zwiers, M. P., Marques, J. P., Chan, K. S., Amelink, J. S., Altgassen, M., Oosterman, J. M., & Norris, D. G. (2024). The Advanced BRain Imaging on ageing and Memory (ABRIM) data collection: Study design, data processing, and rationale. *PLOS ONE*, *19*(6), e0306006. https://doi.org/10.1371/journal.pone.0306006

Jenkinson, M. (1999). *Measuring Transformation Error by RMS Deviation.FMRIB technical report TR99MJ1*.

Jenkinson, M., Bannister, P., Brady, M., & Smith, S. (2002). Improved optimization for the robust and accurate linear registration and motion correction of brain images. *Neuroimage*, *17*(2), 825-841. https://doi.org/10.1016/s1053-8119(02)91132-8

Jenkinson, M., Beckmann, C. F., Behrens, T. E. J., Woolrich, M. W., & Smith, S. M. (2012). FSL. *Neuroimage*, *62*(2), 782-790. https://doi.org/https://doi.org/10.1016/j.neuroimage.2011.09.015
Jenkinson, M., & Smith, S. (2001). A global optimisation method for robust affine registration of brain images. *Med Image Anal*, *5*(2), 143-156. https://doi.org/10.1016/s1361-8415(01)00036-6
Klein, A., Andersson, J., Ardekani, B. A., Ashburner, J., Avants, B., Chiang, M. C., Christensen, G. E., Collins, D. L., Gee, J., Hellier, P., Song, J. H., Jenkinson, M., Lepage, C., Rueckert, D., Thompson, P., Vercauteren, T., Woods, R. P., Mann, J. J., & Parsey, R. V. (2009). Evaluation of 14 nonlinear deformation algorithms applied to human brain MRI registration. *Neuroimage*, *46*(3), 786-802. https://doi.org/10.1016/j.neuroimage.2008.12.037
Liu, Y., & Dawant, B. M. (2014). Automatic detection of the anterior and posterior commissures on MRI scans using regression forests. *Annu Int Conf IEEE Eng Med Biol Soc*, *2014*, 1505-1508. https://doi.org/10.1109/EMBC.2014.6943887
Oxenford, S., Roediger, J., Neudorfer, C., Milosevic, L., Güttler, C., Spindler, P., Vajkoczy, P., Neumann, W.-J., Kühn, A., & Horn, A. (2022). Lead-OR: A multimodal platform for deep brain stimulation surgery. *eLife*, *11*, e72929. https://doi.org/10.7554/eLife.72929
Pallavaram, S., D'Haese, P. F., Lake, W., Konrad, P. E., Dawant, B. M., & Neimat, J. S. (2015). Fully automated targeting using nonrigid image registration matches accuracy and exceeds precision of best manual approaches to subthalamic deep brain stimulation targeting in Parkinson disease. *Neurosurgery*, *76*(6), 756-765. https://doi.org/10.1227/NEU.0000000000000714
Ramirez, J., Scott, C. J., McNeely, A. A., Berezuk, C., Gao, F., Szilagyi, G. M., & Black, S. E. (2014). Lesion Explorer: a video-guided, standardized protocol for accurate and reliable MRI-derived volumetrics in Alzheimer's disease and normal elderly. *J Vis Exp*(86). https://doi.org/10.3791/50887
Schaltenbrand, G., Wahren, W., & Hassler, R. G. (1977). *Atlas for stereotaxy of the human brain* (2d, rev. and enl. ed.). Thieme.
Tian, W., Zhang, J., Chen, J., Liu, Y., Chen, X., & Wang, N. (2016). A quantitative study of intracranial hypotensive syndrome by magnetic resonance. *Clin Neurol Neurosurg*, *141*, 71-76. https://doi.org/10.1016/j.clineuro.2015.12.014
Yoo, T. S., Ackerman, M. J., Lorensen, W. E., Schroeder, W., Chalana, V., Aylward, S., Metaxas, D., & Whitaker, R. (2002). Engineering and algorithm design for an image processing Api: a technical report on ITK--the Insight Toolkit. *Stud Health Technol Inform*, *85*, 586-592. https://www.ncbi.nlm.nih.gov/pubmed/15458157
Zhang, Y., Brady, M., & Smith, S. (2001). Segmentation of brain MR images through a hidden Markov random field model and the expectation-maximization algorithm. *IEEE Trans Med Imaging*, *20*(1), 45-57. https://doi.org/10.1109/42.906424
